# CO_2_ reduction driven by a pH gradient

**DOI:** 10.1101/2020.03.02.973982

**Authors:** Reuben Hudson, Ruvan de Graaf, Mari Strandoo Rodin, Aya Ohno, Nick Lane, Shawn E. McGlynn, Yoichi M.A. Yamada, Ryuhei Nakamura, Laura M. Barge, Dieter Braun, Victor Sojo

## Abstract

All life on Earth is built of organic molecules, so the primordial sources of reduced carbon are a major open question in studies of the origin of life. A variant of the alkaline-vent theory suggests that organics could have been produced by the reduction of CO_2_ via H_2_ oxidation, facilitated by geologically sustained pH gradients. The process would be an abiotic analog—and proposed evolutionary predecessor—of the modern Wood-Ljungdahl acetyl-Co-A pathway of extant archaea and bacteria. The first energetic bottleneck of the pathway involves the endergonic reduction of CO_2_ with H_2_ to formate, which has proven elusive in low-temperature abiotic settings. Here we show the reduction of CO_2_ with H_2_ at moderate pressures (1.5 bar), driven by microfluidic pH gradients across inorganic Fe(Ni)S precipitates. Isotopic labelling with ^13^C confirmed production of formate. Separately, deuterium (^2^H) labelling indicated that electron transfer to CO_2_ did not occur via direct hydrogenation with H_2_. Instead, freshly deposited Fe(Ni)S precipitates appear to facilitate electron transfer in an electrochemical-cell mechanism with two distinct half-reactions. Decreasing the pH gradient significantly, or removing either H_2_ or the precipitate, yielded no detectable product. Our work demonstrates the feasibility of spatially separated, yet electrically coupled geochemical reactions as drivers of otherwise endergonic processes. Beyond corroborating the ability of early-Earth alkaline hydrothermal systems to couple carbon reduction to hydrogen oxidation through geologically plausible and biologically relevant mechanisms, these results may also be of significance for industrial and environmental applications, where other redox reactions could be facilitated using similarly mild approaches.

## Introduction

A dependable supply of reduced organic molecules is essential for any scenario of the origin of life. On the early Earth, one way in which this supply could have been attained was by reduction of CO_2_ with H_2_ (1–5). Geological models indicate that CO_2_ was at comparatively high concentrations in the ocean during the Hadean eon, whereas H_2_ was the product of serpentinization processes in the Earth’s crust and would have emanated as part of the efflux of alkaline hydrothermal vents (3, 6–8). Upon meeting at the vent-ocean interface, the model suggests, the reaction between the two dissolved gases would have produced hydrocarbons, which would in turn take roles in the transition from geochemistry to biochemistry (2, 9). Under standard conditions (1 atm, 25 °C, pH 7), the reaction between CO_2_ and H_2_ to produce formate (HCOO^−^) is thermodynamically disfavored, with ΔG^0^’ = +3.5 kJ mol^−1^ (10, 11). In ancient alkaline vents, however (Figure 1A), H_2_ was present in the OH^−^-rich effluent of the vent, which would have favored its oxidation to water (1). Conversely, CO_2_ would have been dissolved in the relatively acidic ocean, which facilitated protonation in its reduction to HCOO^−^ (1). Assisted by Fe(Ni)S minerals precipitated at the interface, a pH gradient of more than 3 units should be enough to increase the viability of the reaction by ∼180 mV, thereby rendering the reaction favorable (12, 13).

**Figure 1.**
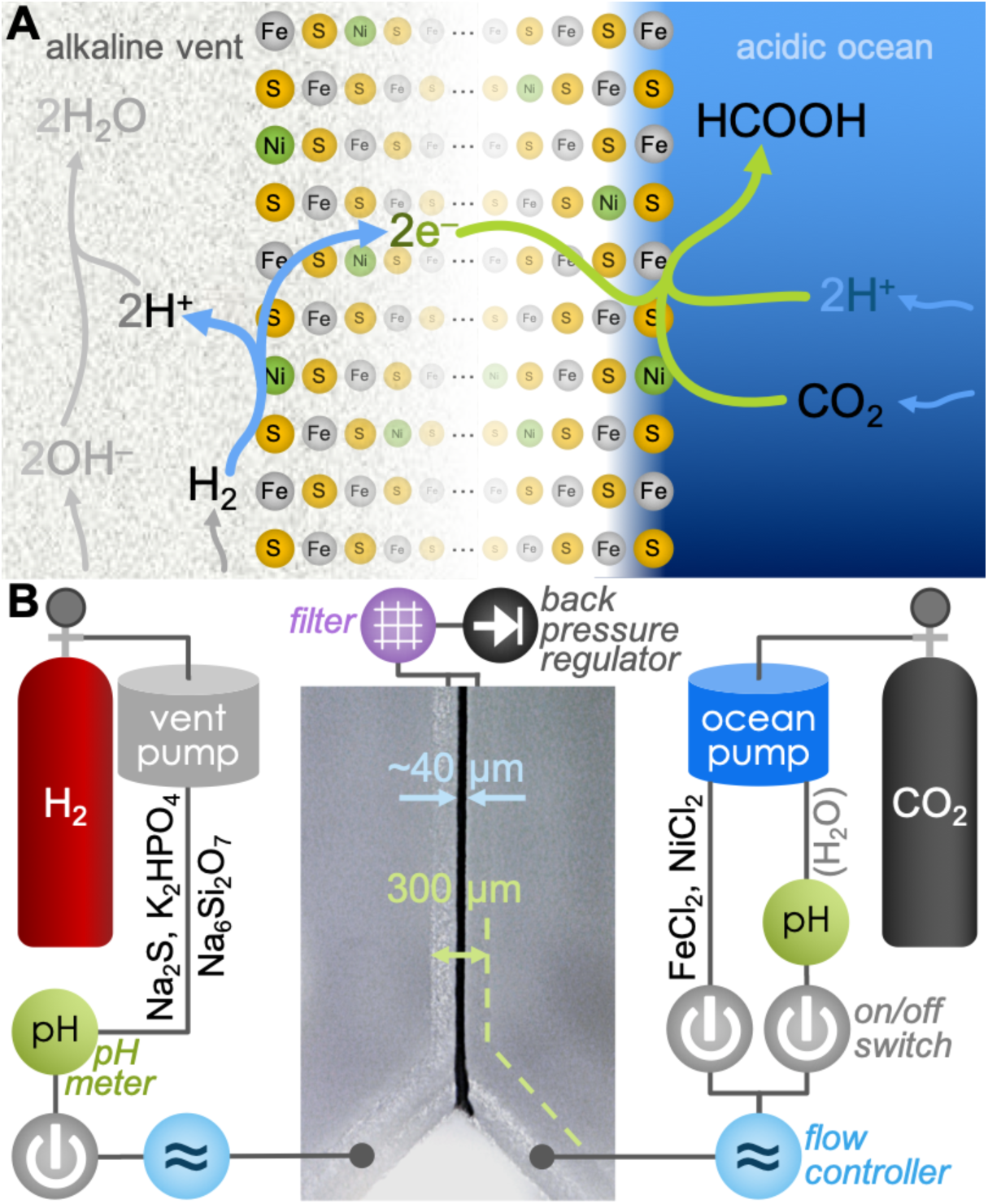
Proposed mechanism of pH-driven CO_2_ reduction with H_2_ across a semi-conducting Fe(Ni)S barrier, and reactor schematic. (**A**) Under alkaline-vent conditions the oxidation of H_2_ (left) is favored by the alkaline pH, due to the presence of hydroxide ions (OH^−^) that react exergonically in the production of water. Released electrons travel across the µm-to-m-thick Fe(Ni)S network (center) to the more oxidizing, acidic solution on the ocean side. There they meet dissolved CO_2_ and a relatively high concentration of protons (H^+^), favoring the production of formic acid HCOOH (or formate HCOO^−^). This electrochemical system enables the overall reaction between H_2_ and CO_2_, which is not observed in batch-solution reaction conditions. (**B**) Diagram of the reactor, with embedded micrograph of a reaction run with precipitate at the interface (further details in the main text and Figure S1).

While there were multiple sources of reduced carbon on the early Earth that may have led to rich chemistries (14–18), the scenario above is especially appealing because of its resemblance to the Wood-Ljungdahl (WL) acetyl Co-A pathway of carbon fixation (2, 19). Highlighting its relevance to studies of the emergence of life, the WL process is the only one of six known biological carbon-fixation pathways that releases energy overall instead of consuming it, and variations of it are present in extant members of both Archaea (methanogens) and Bacteria (acetogens) (2, 5, 9, 10, 20). The first step of the pathway is the reduction of CO_2_ with H_2_ to produce formate (HCOO^−^, or its dehydrated electronic equivalent, CO). This reaction is endergonic, so multiple members of both Archaea and Bacteria use a pH gradient across the cell membrane to power the otherwise unfavorable step (11, 21, 22). In the absence of cellular coupling mechanisms such as electron bifurcation or chemiosmosis, however, this first endergonic step is a key energetic bottleneck of the WL pathway and a major open question for studies of the origin of biological carbon fixation.

Here we show the abiotic indirect reduction of CO_2_ to formate (HCOO^−^) with H_2_, driven by a microfluidic pH gradient across freshly deposited Fe(Ni)S precipitates, via a mechanism that resembles—and may have been the evolutionary predecessor of—the decoupled electron flow of the WL pathway.

## Results

### Precipitation of Fe(Ni)S at the interface in a gas-driven Y-shaped microfluidic reactor

Aiming to coincide with fluid compositions in previous work (4, 23, 24), we prepared an alkaline-vent simulant fluid containing Na_2_S (100 mM), K_2_HPO_4_ (10 mM) and Na_2_Si_3_O_7_ (10 mM) in deaerated water. The counterpart ocean-side fluid contained FeCl_2_ (50 mM) and NiCl_2_ (5 mM). The two fluids were introduced at the tips of a Y-shaped borosilicate microfluidic reactor (Figure 1B). Ambient pressures of H_2_ and CO_2_ have proven insufficient to drive CO_2_ reduction (24, 25), so instead of attempting to dissolve either gas by bubbling prior to the reaction, here we used gas pressure-driven microfluidic pumps. The alkaline fluid was pushed with H_2_ at 1.5 bar, with the ocean analog being pushed with CO_2_ at the same pressure (see reactor schematic in Figure 1B and photograph in Figure S1). We split each reactor run into two consecutive stages, the first for deposition of the Fe(Ni)S precipitates at the interface between the two fluids, and the second (referred to as “post-precipitation” below) for attempting reaction between CO_2_ and H_2_ (or other reagents, as detailed where relevant). Confluence of the ocean and vent analogs during 15 to 60 s yielded a 30 to 60 µm-wide precipitate at the interface between the two fluids, visible under a digital 200x optical microscope (Figure 1B, center).

### Detection of formate and confirmation with ^13^C isotopic labeling

Following formation of the precipitate, and to prevent further precipitation from clogging the microfluidic channels (25), in the second (post-precipitation) stage we switched the ocean fluid to pure deaerated H_2_O, pushed by CO_2_ (Figure 1B, right). The vent simulant was left as before (Na_2_S, K_2_HPO_4_, and Na_2_Si_3_O_7_, pushed by H_2_). Using in-line pH meters, we determined the pH of the ingoing fluids at the point of entry (shortly before going into the reactor; Figure 1B), finding the ocean simulant at pH 3.9 and the vent simulant at pH 12.3. At a flow rate of 5 µL/min for each input, the residence time of fluids within the central channel was a maximum of 3.3 s, so we allowed the system to flow for at least 2 min before collecting output. We then collected a single (i.e. mixed) output of the reactor over the following 50 min, and later analyzed it by nuclear magnetic resonance (NMR) spectroscopy. Under these conditions, we detected formate (HCOO^−^) by both ^1^H and ^13^C NMR (Table 1, experiment #1), at a concentration of 1.5 µM. Singlet peaks in the ^1^H (8.42 ppm, Figure 2A) and ^13^C (165.8 ppm, Figure S2) NMR spectra matched samples of pure (>98%) formic acid (Figure S3). Keeping the ocean solution free of metal salts from the start—i.e., in the absence of the precipitate—gave no detectable product (see S.I.). Running both the precipitation and reaction stages with isotopically enriched (99% ^13^C) ^13^CO_2_ (Table 1, experiment #2) gave a stronger singlet in the ^13^C spectrum (165.8 ppm, Figure 2B), and yielded the expected splitting of the formyl singlet into a doublet in the ^1^H spectrum (*J* = 195 Hz), due to the ^1^H–^13^C coupling in the formyl group (Figure 2C).

**Figure 2.**
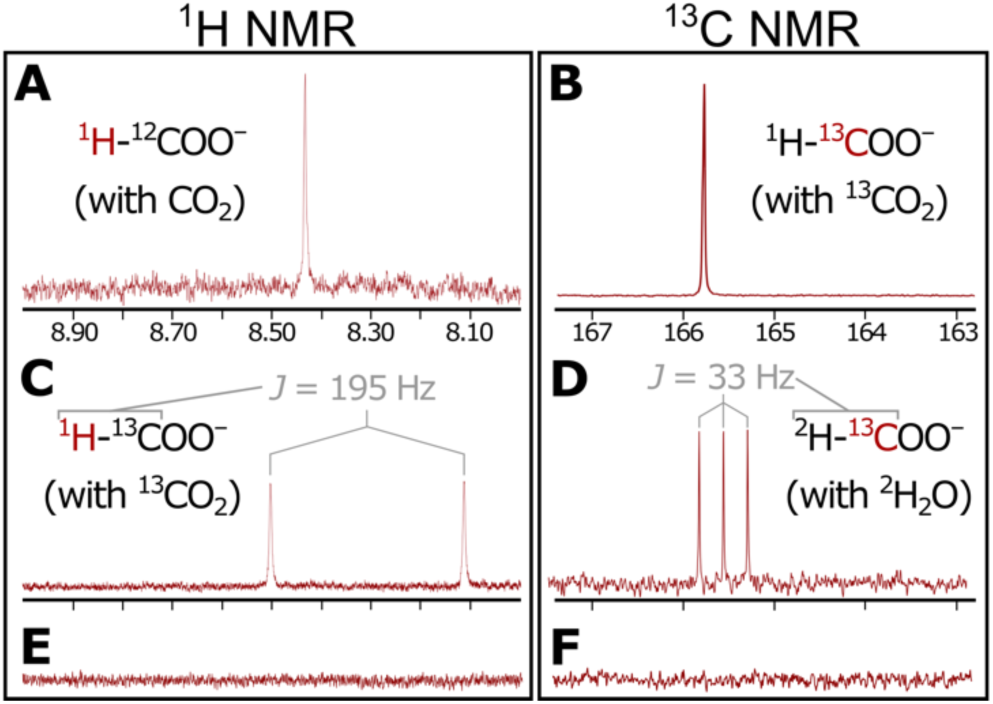
^1^H and ^13^C NMR spectra of produced formate. Singlets in the ^1^H (**A**) and ^13^C (**B**) NMR spectra demonstrate the production of formate in our system. (**C**) With isotopically labeled ^13^CO_2_, coupling between ^1^H and ^13^C produced an expected doublet in the ^1^H spectrum. (**D**) Replacing regular water with deuterated ^2^H_2_O on the ocean side yielded a triplet in the ^13^C spectrum due to coupling between the ^2^H and ^13^C nuclei. (**E**, **F**) No formate was detected upon replacing H_2_ with N_2_ as the vent-driving gas. Corresponding entries in Table 1 are A:1, B:2, C:2, D:5. E:3, F:3.

**Table 1.**
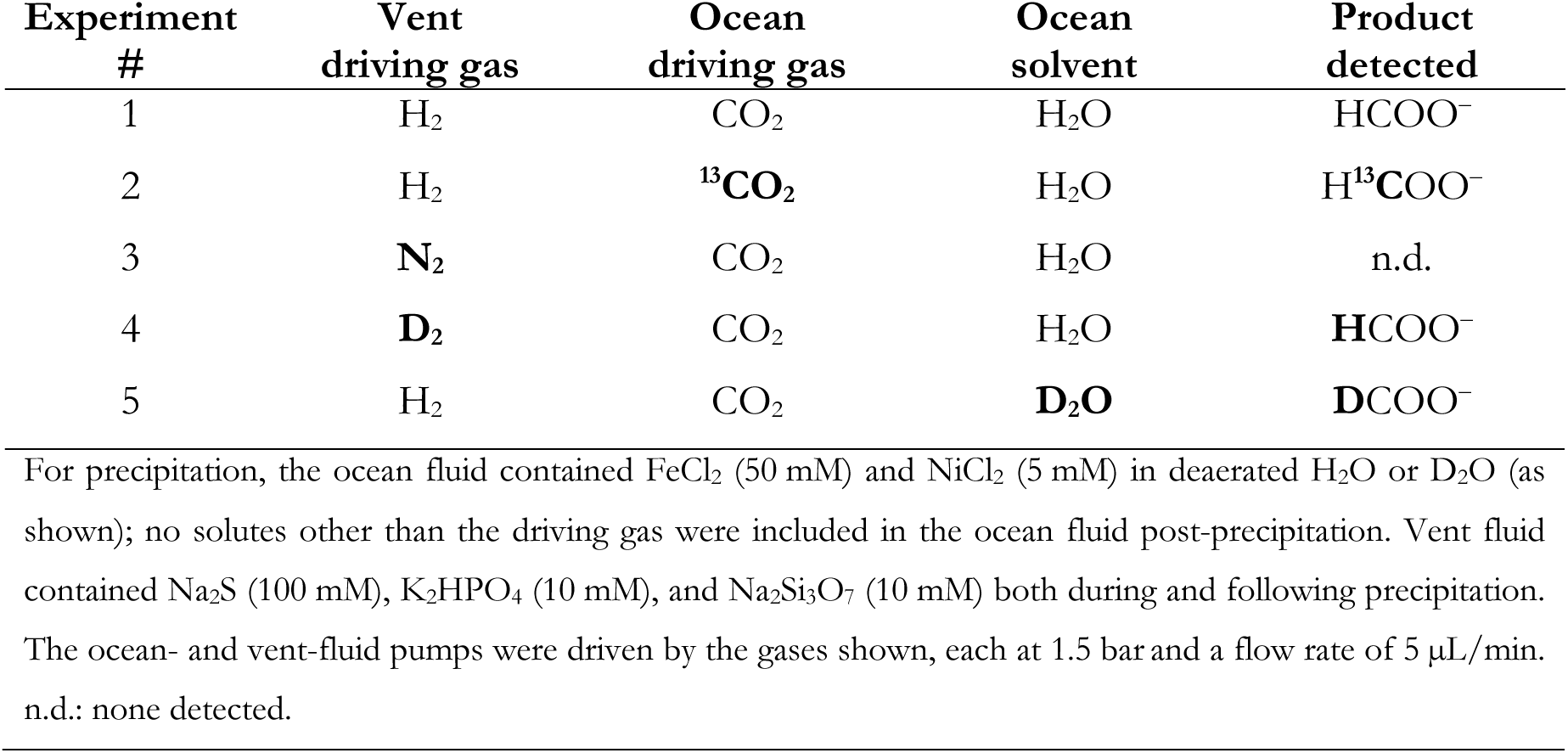
Mechanistic analysis of CO_2_ reduction^a^.

H_2_ appears to be necessary for CO_2_ reduction: with the vent-side fluid driven by N_2_ instead of H_2_ (i.e. in the absence of H_2_ both during and following precipitation), we observed no reduction products (experiment #3 and Figure 2E-F).

### Evidence for an indirect pH-dependent electrochemical mechanism of CO_2_ reduction driven by H_2_ oxidation

In search of mechanistic insight, we conducted deuterium-labeling (^2^H, or D) experiments (Table 1, experiments #4-5), using the isotopic variants throughout the entire experiment. Regardless of whether we used unlabeled H_2_ (experiment #1) or D_2_ (experiment #4) to drive the vent-side pump, we observed only non-isotopically labeled formate (HCOO^−^) in the efflux, suggesting that CO_2_ reduction may be occurring exclusively on the ocean side. Conversely, with D_2_O used in place of regular H_2_O on the ocean side, and with unlabeled H_2_ driving the vent-side pump (experiment #5), we detected only DCOO^−^ (as evidenced by a triplet in ^13^C NMR, *J* = 33 Hz, and the lack of any other discernible peaks; Figure 2D). This further confirms that CO_2_ reduction here matched the isotopic composition of the ocean side, not the vent side.

We next explored the role of the inherent pH gradient of the simulated submarine alkaline hydrothermal system. The successful reductions reported in Table 1 proceeded with an initial ocean-simulant pH of 3.9, whereas the vent analog was at pH 12.3. Upon mixing, this initial ΔpH of 8.4 units would have inevitably dropped, but we have previously shown that pH gradients of multiple units hold successfully over time across microfluidic scales, particularly in the presence of a precipitate at the interface (23). We aimed to determine whether such a pH gradient is required in our reduction system to facilitate the oxidation of H_2_ on the alkaline side and the reduction of CO_2_ on the acidic side (Figure 1A). Following precipitation under the same conditions as above for experiment #1 in Table 1 (repeated in Table 2), we evaluated the effects of a number of changes to the pH and composition of each of the two fluids (Table 2). Replacement of the vent simulant with pure H_2_O driven by H_2_ afforded no product (Table 2, experiment #6). Leaving the Na_2_S, K_2_HPO_4_ and Na_2_Si_3_O_7_ in the vent solution and H_2_ as the driver gas, but acidifying the vent solution with HCl to pH 3.9 and pH 7.0 likewise resulted in no detectable formate production (Table 2, experiments #7 and #8, respectively). Adding 100 mM Na_2_CO_3_ to the ocean fluid while still using CO_2_ as the driving gas (Table 2, experiment #9) raised the ocean-side pH to 9.8; we detected no product under these conditions. Removing silicate from the vent side after precipitation still yielded formate (Table 2, experiment #10), as did removing both silicate and phosphate to have only Na_2_S (Table 2, experiment #11). With only K_2_HPO_4_ in the vent side post-precipitation we detected only trace amounts of formate (below our limit of quantification of 0.37 µM; see S.I. Methods), possibly due to the insufficiently alkaline pH of 9.1 (Table 2, experiment #12). The more alkaline K_3_PO_4_ raised the pH to 12.1 and led to production of considerably more formate (Table 2, experiment #13). These results simultaneously confirm the role of the pH gradient and show that a continuous supply of aqueous sulfide is not required in our system (this however does not preclude the possibility of precipitate-bound sulfide acting as a stoichiometric reductant in addition to H_2_). Removing Ni from the ocean precipitation fluid (experiment #14) produced only small amounts of formate (below our limit of quantification of 0.37 µM). Conversely, replacing Fe to have Ni as the only metal in the ocean precipitation fluid (NiCl_2_ 55 mM, experiment #15) yielded 1.4 µM formate, pointing toward a crucial role for Ni within the precipitates. Removing both FeCl_2_ and NiCl_2_ from the ocean fluid expectedly did not produce a precipitate and did not result in the production of detectable formate (experiment #16).

**Table 2.**
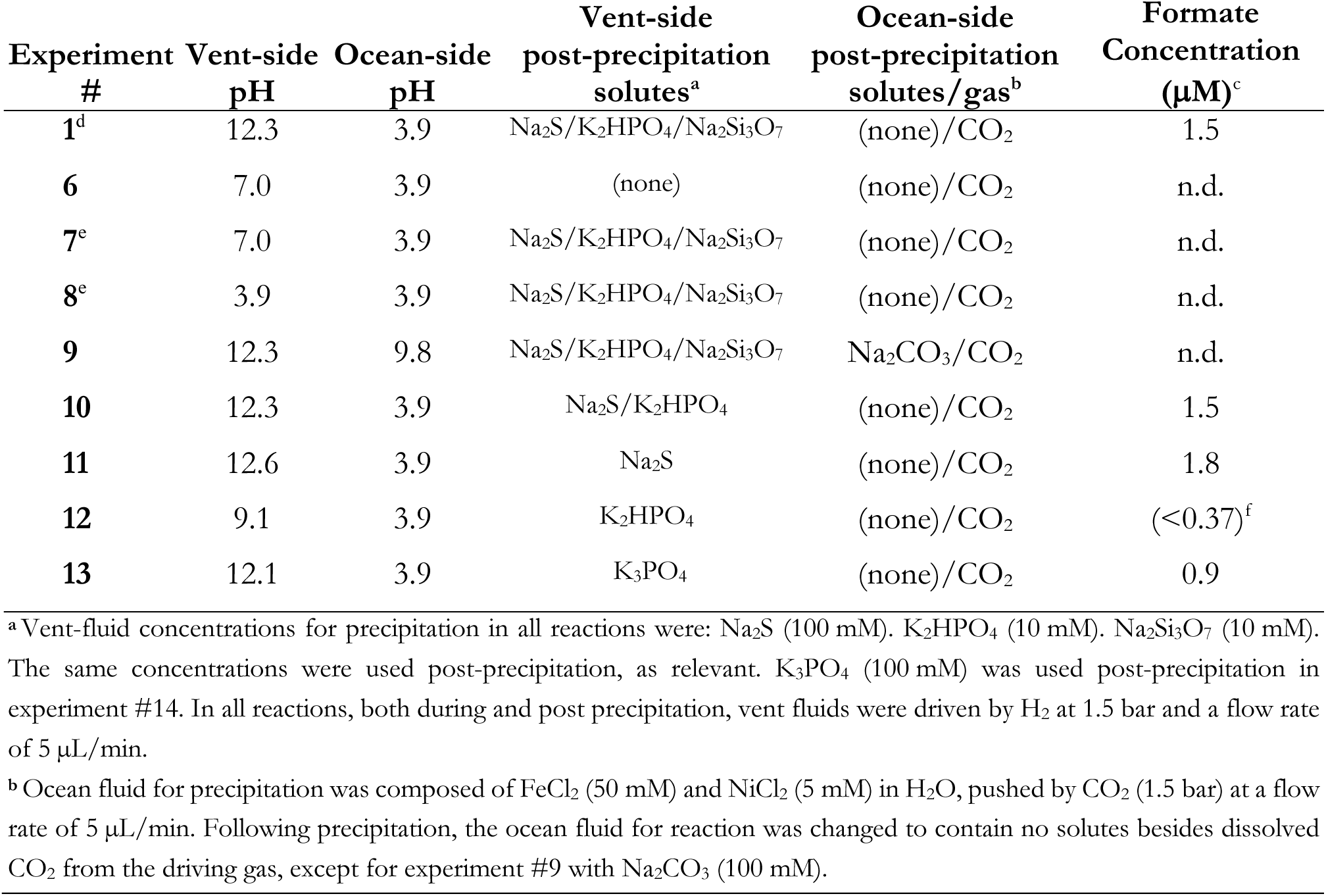
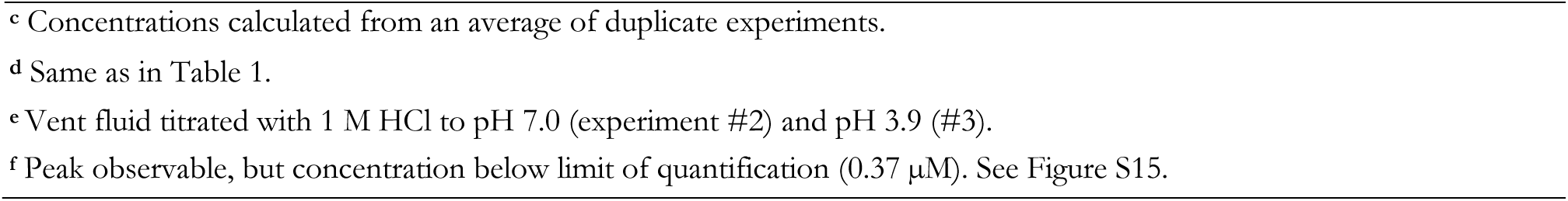
Exploration of the role of the microfluidic pH gradient across the mineral precipitate in CO_2_ reduction.

| Experiment # | Vent-side pH | Ocean-side pH | Vent-side post-precipitation solutes<sup>a</sup> | Ocean-side post-precipitation solutes/gas<sup>b</sup> | Formate Concentration ($\mu M$)<sup>c</sup> |
| --- | --- | --- | --- | --- | --- |
| 1<sup>d</sup> | 12.3 | 3.9 | $Na_2S/K_2HPO_4/Na_2Si_3O_7$ | (none)/$CO_2$ | 1.5 |
| 6 | 7.0 | 3.9 | (none) | (none)/$CO_2$ | n.d. |
| 7<sup>e</sup> | 7.0 | 3.9 | $Na_2S/K_2HPO_4/Na_2Si_3O_7$ | (none)/$CO_2$ | n.d. |
| 8<sup>e</sup> | 3.9 | 3.9 | $Na_2S/K_2HPO_4/Na_2Si_3O_7$ | (none)/$CO_2$ | n.d. |
| 9 | 12.3 | 9.8 | $Na_2S/K_2HPO_4/Na_2Si_3O_7$ | $Na_2CO_3/CO_2$ | n.d. |
| 10 | 12.3 | 3.9 | $Na_2S/K_2HPO_4$ | (none)/$CO_2$ | 1.5 |
| 11 | 12.6 | 3.9 | $Na_2S$ | (none)/$CO_2$ | 1.8 |
| 12 | 9.1 | 3.9 | $K_2HPO_4$ | (none)/$CO_2$ | (<0.37)<sup>f</sup> |
| 13 | 12.1 | 3.9 | $K_3PO_4$ | (none)/$CO_2$ | 0.9 |

<sup>a</sup> Vent-fluid concentrations for precipitation in all reactions were: $Na_2S$ (100 mM), $K_2HPO_4$ (10 mM), $Na_2Si_3O_7$ (10 mM). The same concentrations were used post-precipitation, as relevant. $K_3PO_4$ (100 mM) was used post-precipitation in experiment #14. In all reactions, both during and post precipitation, vent fluids were driven by $H_2$ at 1.5 bar and a flow rate of $5\ \mu L/min$ .
<sup>b</sup> Ocean fluid for precipitation was composed of $FeCl_2$ (50 mM) and $NiCl_2$ (5 mM) in $H_2O$ , pushed by $CO_2$ (1.5 bar) at a flow rate of $5\ \mu L/min$ . Following precipitation, the ocean fluid for reaction was changed to contain no solutes besides dissolved $CO_2$ from the driving gas, except for experiment #9 with $Na_2CO_3$ (100 mM).
<sup>c</sup> Concentrations calculated from an average of duplicate experiments.
<sup>d</sup> Same as in Table 1.
<sup>e</sup> Vent fluid titrated with 1 M HCl to pH 7.0 (experiment #2) and pH 3.9 (#3).
<sup>f</sup> Peak observable, but concentration below limit of quantification (0.37 μM). See Figure S15.

NMR spectra and further analyses for all experiments discussed here are presented in S.I. Figures Figure S2–Figure S19. Images of the precipitates before and after a number of the reactions can be seen in Figure S20.

## Discussion

We report the abiotic pH-driven reduction of CO_2_ to formate with H_2_ in a minimalistic microfluidic replicant of an alkaline hydrothermal system. Removing the pH gradient entirely or significantly reducing it—either by acidifying the vent fluid or alkalinizing the ocean fluid—yielded no detectable formate. With the acidic-ocean side fixed at pH 3.9, we observed greater formate yields with increasing vent-side pH. Respectively for experiments 7, 12, 13, 1 and 11 in Table 2, as the vent-side pH rose (7.0 < 9.1 < 12.1 < 12.3 < 12.6), so did the average formate concentration measured (undetected < below 0.37 µM < 0.9 µM < 1.5 µM < 1.8 µM).

Removing the precipitate entirely also yielded no product. Without nickel in the ocean precipitation fluid, the yield dropped below our limit of quantification, suggesting that Ni is a crucial part of the precipitate for the reduction mechanism operating here. Given that Ni can typically serve as a more efficient classical and ionic hydrogenation catalyst than Fe (26), the requirement for Ni within the precipitate is not surprising. Although we do not invoke such a hydrogenation mechanism here, the electrochemical mechanism that we propose in Figure 1A would rely on the metal center for similar catalytic functions as those required during classical or ionic hydrogenations.

Removing H_2_ entirely by replacing it with N_2_ as the vent-driving gas—while still keeping Na_2_S in the vent fluid—similarly led to no detectable product. Conversely, we do observe successful formate production with Na_2_S absent from the vent fluid post-precipitation and H_2_ as vent-driving gas (Experiment #14). Together, these results suggest that H_2_ is a necessary reagent in our mechanism and that sulfide is not sufficient as a reducing agent. It is also noteworthy that experiment #11 with only Na_2_S in the post-precipitation vent fluid yielded a higher concentration of formate than experiment #1, despite containing the same concentration of sulfide; we attribute this result to the higher vent-side pH in the absence of K_2_HPO_4_ and Na_2_Si_3_O_7_. In all, and while we cannot presently affirm that the precipitates are acting only as catalysts, our results suggest that H_2_ is the main electron donor, that a large pH gradient is necessary for its oxidation, and that sulfide is insufficient (and might not be required) as an electron donor.

We have used 1.5 bar pressures of both H_2_ and CO_2_, which are nominally not too dissimilar from the ambient-pressure bubbling employed in similar earlier work that did not yield any detectable reduced carbon (24, 25). In those experiments, the two gases were dissolved into the fluids prior to the reactions. Recent calculations suggest that atmospheric pressures of H_2_ and CO_2_ are only marginally insufficient to drive the reaction to formate (25). We therefore infer that the positive results reported in this work are most plausibly due to the advantages of the gas-driven system used here, which likely result in higher dissolved gas concentrations than the atmospheric-pressure bubbling performed previously (24, 25). Chiefly, the ability to keep the gases consistently within the system, as well as more reliably maintaining the fluids anoxic, may suffice as an explanation for the positive results observed here in contrast with previous attempts.

### Alternative mechanisms for the reduction of CO_2_ linked to H_2_ oxidation

Several CO_2_-reduction mechanisms are possible for a laminar-flow system such as the one that we have used. We briefly explore a number of them below and discuss why we deem them less likely than the electrochemical mechanism suggested in Figure 1A. Further details are presented in the S.I. (Figures Figure S21–Figure S26).

Perhaps the most chemically intuitive—albeit least biochemically homologous—mechanisms of carbon reduction possible here are direct hydrogenations (Figures Figure S21–Figure S23), in which hydrogen from H_2_ would be transferred directly to CO_2_ (27–31), either as atomic hydrogen (classical hydrogenation) or as hydride (ionic hydrogenation). Most simply, the product in such a mechanism should match the isotopic signature of the H_2_/D_2_ vent gas. Instead, the produced formate matches only the isotopic make-up of the ocean-side water, regardless of the composition of the vent-side gas or water. In these direct hydrogenation mechanisms, adsorbed hydrogen species could in principle exchange with the surrounding fluid, such that we lose the original isotopic signature (32). However, any such process would imply migration of significant amounts of fluid across the precipitate. The substantial ensuing mixing of fluids should have given a scrambled H/D formyl signal, which we do not observe, all but ruling out the direct-hydrogenation mechanisms.

Alternatively, the hydrogen atoms in the resultant formate may not derive directly from H_2_. Instead, the mechanism may proceed via redox cycling of an edge or corner Fe or Ni atom (M^2+^ ⇄ M^0^) wherein a metal is first reduced by H_2_ (leaving 2 protons to dilute away), and subsequently the metal transfers the acquired electrons on to CO_2_, with accompanying proton abstraction from the local aqueous environment (Figures Figure S24–Figure S26). Noteworthy for any mechanisms that rely on proton transfer, none of our experiments contained acids added to the ocean side. The acidic pH of 3.9 that we report was achieved solely by dissolution of CO_2_ in water. Thus, all ocean-side protons must derive from the dissociation of carbonic acid via:

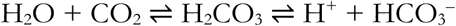

When our reaction is conducted with D_2_O (experiment #5) as the ocean solvent, we find only deuterated formate (DCOO^−^) in the efflux, suggesting that CO_2_ reduction did not occur on the vent side (which had normal H_2_ as the feed gas and normal H_2_O as the solvent, and so would have yielded unlabeled HCOO^−^ as the product). This is confirmed in experiment #4 with D_2_ as vent-driving gas, which yielded only unlabeled HCOO^−^. Several scenarios for such a localized redox cycling are possible (Figures Figure S24–Figure S26), but because all require co-location, none could feasibly offer the exclusively ocean-side isotopic signature that we observe.

As an alternative to the above catalytic mechanisms (classical and ionic hydrogenations, or localized redox cycling), a non-catalytic (i.e. stoichiometric) precipitate oxidation could be driving CO_2_ reduction. However, the lack of reaction in the aqueous sulfide-containing but H_2_-free experiment #3 suggests that there is a requirement for a continuous supply of electrons from H_2_—not from dissolved sulfide or the precipitate itself.

Together with the strong pH dependence of the reactions shown in Table 2, these observations suggest that the reduction of CO_2_ here proceeds via a de-coupled electrochemical mechanism in which electrons from H2 oxidation on the alkaline-vent side are shuttled across the Fe(Ni)S precipitates toward CO_2_ on the acidic-ocean side (Figure 1A). There, the reacting CO_2_ picks up a proton from the local water, yielding formate with the isotopic signature of the ocean side.

### Venturi pull inside hydrothermal pores would bring ocean fluids into the vent system

We show that production of organics could have occurred on the ocean side of an ancient alkaline hydrothermal-vent system. This raises the question of whether these organics would simply dilute away into the ocean before they could take any biochemical role (33), and it is conspicuously unlike the WL pathway (25), in which carbon reduction occurs inside the cell. Because the microscopic structure of hydrothermal vents is typically highly porous and reticulated (8, 34), we hypothesize that ocean fluids could have been actively pulled into the microfluidic system due to Venturi-effect suction. Once inside, the ocean’s carbonic fluids could react with electrons being transferred across the vent’s catalytic minerals, and fresh precipitation could also occur further in as the two fluids met. We have developed a computational finite-elements simulation to explore this prediction (see S.I. and Figure S27 for details). Results show that a system of a size similar to that of our 300 µm-wide reactor would indeed lead to the type of microfluidic confluence of reagents that we have shown here and elsewhere (23, 24, 35). Notably, such an effect is not limited to submarine alkaline vents, and will likely occur within porous hydrothermal systems at any location and depth, lending itself to multiple geochemical scenarios for the emergence of life.

### Batch vs. flow reactors

An alternative one-pot batch (rather than microfluidic) system for the reduction of CO_2_ with H_2_ has been reported recently (36), elaborating on previous results (37, 38). Using more highly reduced minerals and higher pressures (10 bar H_2_) than the ones employed here, the batch system generates significantly higher concentrations of formate, as well as multiple further reduction products (including acetate, methanol and pyruvate). However, the close resemblance to biological mechanisms—such as the pH-driven CO_2_ reduction demonstrated here—added to the possibility of thermal-driven concentration increases (39–41), makes microfluidic reactors attractive, in spite of the presently low yields in our flow system. The two types of system thus provide complementary compelling evidence for organics arising under anoxic alkaline hydrothermal-vent conditions, and both must be explored further as potential sources for life’s first molecules.

## Conclusions

We report the pH-driven reduction of CO_2_ with H_2_, in the first step of a geologically plausible analog—and proposed evolutionary predecessor (2)—of the Wood-Ljungdahl acetyl-CoA pathway for carbon fixation.

These results tie in with previous findings that alkaline vent-like pH gradients can be kept at the microscale across Fe(Ni)S precipitates (23), that these pH gradients can provide the necessary overpotentials to drive the reaction between H_2_ and CO_2_ (12), that low pressures of the two gases are insufficient for reaction (24, 25), that heat gradients in hydrothermal systems can drive concentration increases by thermophoresis and capillary flow at gas-liquid interfaces (39–41), and that redox and pH gradients can drive early bioenergetics (42, 43) and amino acid synthesis (44).

Both dissolved hydrogen gas and a significant pH gradient of the type and polarity found at alkaline hydrothermal vents appear to be necessary driving forces in our system. Further, our findings suggest an electrochemical mechanism whereby dissolved H_2_ oxidizes on the alkaline-vent side and its electrons travel through the catalytic precipitate network toward the acidic-ocean side, where they reduce dissolved CO_2_ to formate. To date, such an experimental confirmation has proven elusive (4, 24, 25). Since this is the first endergonic hurdle in the reduction of CO_2_ via the only energetically profitable CO_2_ fixation pathway known, our findings provide a geologically credible route to a plausible origin of carbon fixation at the emergence of life—on the early Earth and potentially on water-rock planetary bodies elsewhere.

## Methods

Minimal simulants of alkaline-vent and oceanic fluids were driven into respective tips of a Y-shaped borosilicate reactor. To maximize the amounts of gases dissolved, we used gas-pressure-driven microfluidic pumps (Dolomite Mitos P-pump). The ocean fluid was pushed with CO_2_ and the vent fluid with H_2_, both at 1.5 bar. The reaction was undertaken in an anaerobic N_2_-purged glovebox. Single (mixed) effluxes were collected and analyzed using ^1^H and ^13^C NMR. Quantification was achieved with ^1^H NMR. Further descriptions of experimental, analytical and computational methods are presented in the S.I.

## Author contributions

RH and VS designed the experiments, with contributions from all authors. RH, RdG, MSR and VS performed the experiments. RH performed NMR experiments. VS developed the computational simulation. All authors analyzed the results and wrote the manuscript.

## Acknowledgements

RH, LMB and VS acknowledge support from the NASA Maine Space Grant Consortium (SG-19-14 & SG-20-19). RH is supported by NSF Award 1415189 and acknowledges support from the Japan Society for the Promotion of Science (JSPS) (FY2016-PE-16047). SEM is supported by NSF Award 1724300. VS acknowledges support from JSPS (FY2016-PE-16721), the European Molecular Biology Organization (EMBO) (ALTF-1455-2015), the Institute for Advanced Study in Berlin, and the Gerstner Family Foundation.

## Supporting Information

### 1. Further Methods

#### 1.1. Reactor parts and assembly

The reactor consisted of the following parts (see Figure S1):

- **Pressure-driven microfluidic pumps:** Dolomite Mitos P-pump (Dolomite part #:3200175)
- **Pump lids:** Mitos P-pump 3-way chamber lid (to enable seamless transition between different fluids on both the ocean and vent sides) (Dolomite part #: 3200044)
- **Valves:** 2-way in-line valves (Dolomite part #: 3200087)
- **Microfluidic chips:** Micronit H-Microreactor (borosilicate) (Micronit part #: 00755)
- **Microfluidic chip holder:** Micronit Fluidic Connect 4515 chipholder (Micronit part #: FC_FC4515)
- **T connectors:** Flow splitter 1/16” OD T-shape (3 connections) (Micronit part #: 01628)
- **In-line pH meters:** Sensorex pH electrode (Sensorex part #: 970277) with Modified Flow Cell FC49K, 50uL, PVDF (Sensorex part #: 970282).
- **Optical microscope:** Opti-Tekscope Digital USB Microscope Camera – Advanced CMOS Sensor, True High Definition Macro 200x Zoom Imaging

**Figure S1.**
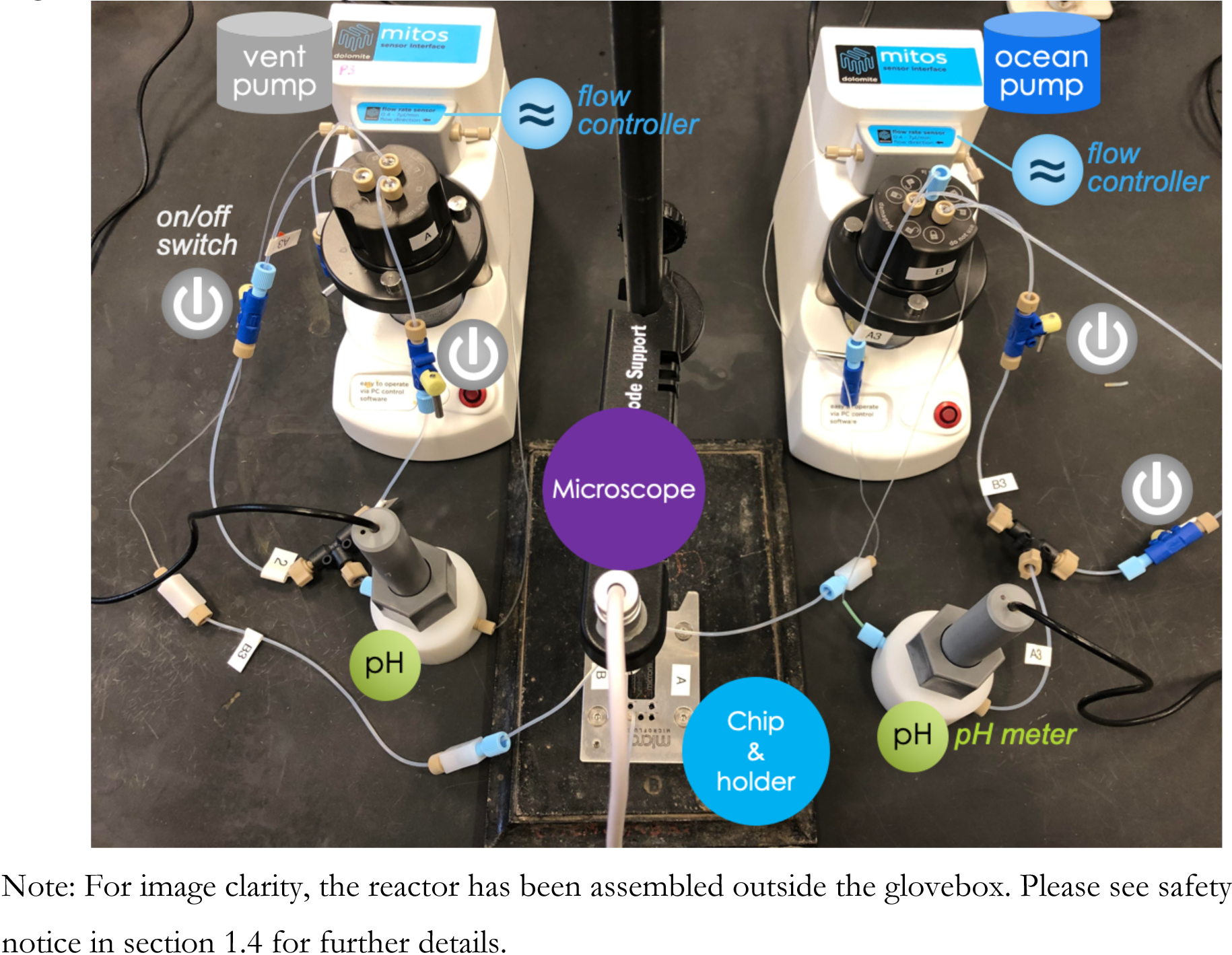
Reactor images.

The microfluidic reactor systems were assembled using borosilicate Y-junction chips (Micronit H-reactor). The channels in the system were half-pipes with a width of 300 ± 4.0 µm and a depth of 150 ± 5 µm. Simulants of ancient vent efflux and ocean fluids were entered at the two inputs of the Y-junction. A single, mixed output was collected. The fluid compositions, summarized in Table S1, were similar to those reported elsewhere (4, 24, 25). A general schematic of the reactor is provided in Figure 1B of the main text, and a photograph in Figure S1. A safety notice relating to work with pressurized H_2_ is presented below in section 1.4.

In order to maximize the amount of dissolved gas (ocean side: CO_2_; vent side H_2_), we designed the microfluidic system to operate on gas pressure-driven pumps (Dolomite Mitos P-pump). Prior to driving the ocean and vent simulant fluids through the microfluidic device, we subjected the chambers to 10 pressurization-depressurization cycles (5 bar) in order to fully replace the headspace gas with the desired H_2_ or CO_2_. The final pressurization was held at the desired driving pressure (1.5 bar) for 10 min, prior to allowing the fluids to flow into the reactor.

To establish parallel flow prior to setting the Fe(Ni)S precipitate, we flowed the vent fluid (Na_2_S, K_2_HPO_4_, Na_2_Si_3_O_7_ in water; driven by H_2_) alongside a metal-free ocean fluid (water; driven by CO_2_), each at a flow rate of 5 µL/min. Control samples were taken at this point, resulting in no detectable product (presented as experiment #16 in Table 2 of the main text). Upon establishment of the parallel flow, we introduced the metal-containing ocean fluid (FeCl_2_ and NiCl_2_ in water; driven by CO_2_). Invariably, within seconds of introducing the metal-containing ocean fluid, a black precipitate formed at the boundary of the laminar flow, which grew until the metal-containing ocean flow was replaced by the metal-free ocean flow (switched back to avoid the precipitate from growing to the point of occluding the channel). This switch from metal-containing to metal-free fluids typically occurred within 15-60 s of initial precipitate formation. Throughout the experiment, we maintained a flow rate of 5 µL/min for each the ocean-and vent-replicant fluids (10 µL/min overall combined flow).

Mimicking early-Earth anoxic conditions, both the vent efflux and ocean fluids were prepared in an N_2_-purged glovebox with degassed, distilled water. To degas, water was sparged with N_2_ for 30 min and subjected to three freeze-pump-thaw cycles. The gas-driven system we used is fully air-tight so, after preparation of the solutions in the anaerobic glovebox, the reaction was carried out outside. However, to provide an additional safety measure when working with pressurized H_2_, the vent-side pump was housed within the N_2_-purged glovebox throughout the experiment (see safety notice in section 1.4).

#### 1.2. Maximum estimated dissolved H_2_ and CO_2_ concentrations

It was not possible in our system to determine the exact concentrations of dissolved H_2_ and CO_2_ in the vent and ocean fluids, respectively. However, using Henry’s law, we estimate the maximum possible concentrations under our conditions to be 52 mM CO_2_ and 1.2 mM H_2_. Henry’s law constants were obtained from the National Institute of Standards and Technology’s WebBook at https://webbook.nist.gov, with ID=C124389 for CO_2_ and ID=C1333740 for H_2_.

#### 1.3. Residence times and flow speeds

Given that the volume of the common channel of the chip is reported by the manufacturers as 0.55 µL, and that the fluids have a combined 10 µL/min flow rate (equivalent to 0.1667 µL/s), we calculate a total residence time of 3.3 s. The length of the common channel is reported as 14.8 mm, which gives a linear speed of 4.48 mm/s.

#### 1.4. Safety notice

The Dolomite Mitos P-pumps are not designed for use with reactive gases (such as the highly combustible H_2_). While the electronics are sealed away from the pressure chamber, a failed seal coupled with a spark from the electronics could result in H_2_ combustion and rupture of the chamber. As a safety precaution, when using pressurized H_2_ we operated the vent-side pump within a nitrogen-purged glovebox. This way, even if a pump seal failed and the electronics sparked, there would not be enough ambient O_2_ to enable H_2_ combustion.

#### 1.5. Reagents and concentrations

- Iron (II) chloride tetrahydrate (99%) (Oakwood Chemicals).
- Nickel (II) chloride (98%) (Oakwood Chemicals).
- Sodium sulfide nonahydrate, ACS, (98.0% min) (Oakwood Chemicals).
- H_2_ gas (UHP) (Maine Oxy).
- CO_2_ gas (beverage grade: 99.95%) (Maine Oxy).
- N_2_ gas (UHP) (Maine Oxy).
- Carbon-^13^C Dioxide (99 Atom % ^13^C) (Isotec/Sigma Aldrich).
- Deuterium (99.8 Atom %) (Isotec/Sigma Aldrich).
- K_2_HPO_4_ (ACS reagent > 98%) (Sigma Aldrich).
- K_3_PO_4_ (ACS reagent > 98%) (Sigma Aldrich).
- Na_2_CO_3_ (ACS reagent > 98%) (Sigma Aldrich).

**Table S1.** Ocean and vent analog default fluid compositions

| Ocean-analog fluid |  | Vent-analog fluid |  |
| --- | --- | --- | --- |
| FeCl<sub>2</sub> | 50 mM | Na<sub>2</sub>S | 100 mM |
| NiCl<sub>2</sub> | 5 mM | K<sub>2</sub>HPO<sub>4</sub> | 10 mM |
|  |  | Na<sub>2</sub>Si<sub>3</sub>O<sub>7</sub> | 10 mM |
| CO<sub>2</sub> | 1.5 bar | H<sub>2</sub> | 1.5 bar |

Note: Once the precipitate was formed, the FeCl<sub>2</sub> and NiCl<sub>2</sub> were removed from the ocean-fluid input.

#### 1.6. NMR analysis

Formate was identified using ^1^H and ^13^C NMR, and quantified using ^1^H NMR. The post-chip effluent was centrifuged, of which 400 µL was added to an NMR tube with 100 µL of D_2_O. For quantification, acetone was used as an internal standard (Figures Figure S2 and Figure S3), added immediately prior to analysis. Quantification was achieved by integration of the formyl peak in comparison against the known concentration for the six methyl protons of the acetone peak. The identity and concentration of formic acid were confirmed by spiking the outflow solution with additional formic acid, and noting the growth of the existing ^1^H and ^13^C peaks (Figure S2) rather than the introduction of new peaks (24). To confirm quantification results, re-integration of the formyl ^1^H peak (8.42 ppm) in comparison with the acetone internal standard (2.22) followed after the known amount of additional formic acid was added. ^1^H NMR spectra were conducted with water suppression for 256 scans. ^13^C spectra were collected for 20,000-40,000 scans. Our limit of quantification for formate was 0.37 µM (see section 5). On a standard curve with acetone (0.6 µM) as an internal standard, concentrations of formate below this concentration deviated from linearity (0.037 µM & 0.0037 µM). NMR spectra were acquired on a Bruker 500 MHz NMR spectroscope.

### 2. ^13^C NMR spectrum of Experiment 1

**Figure S2.**
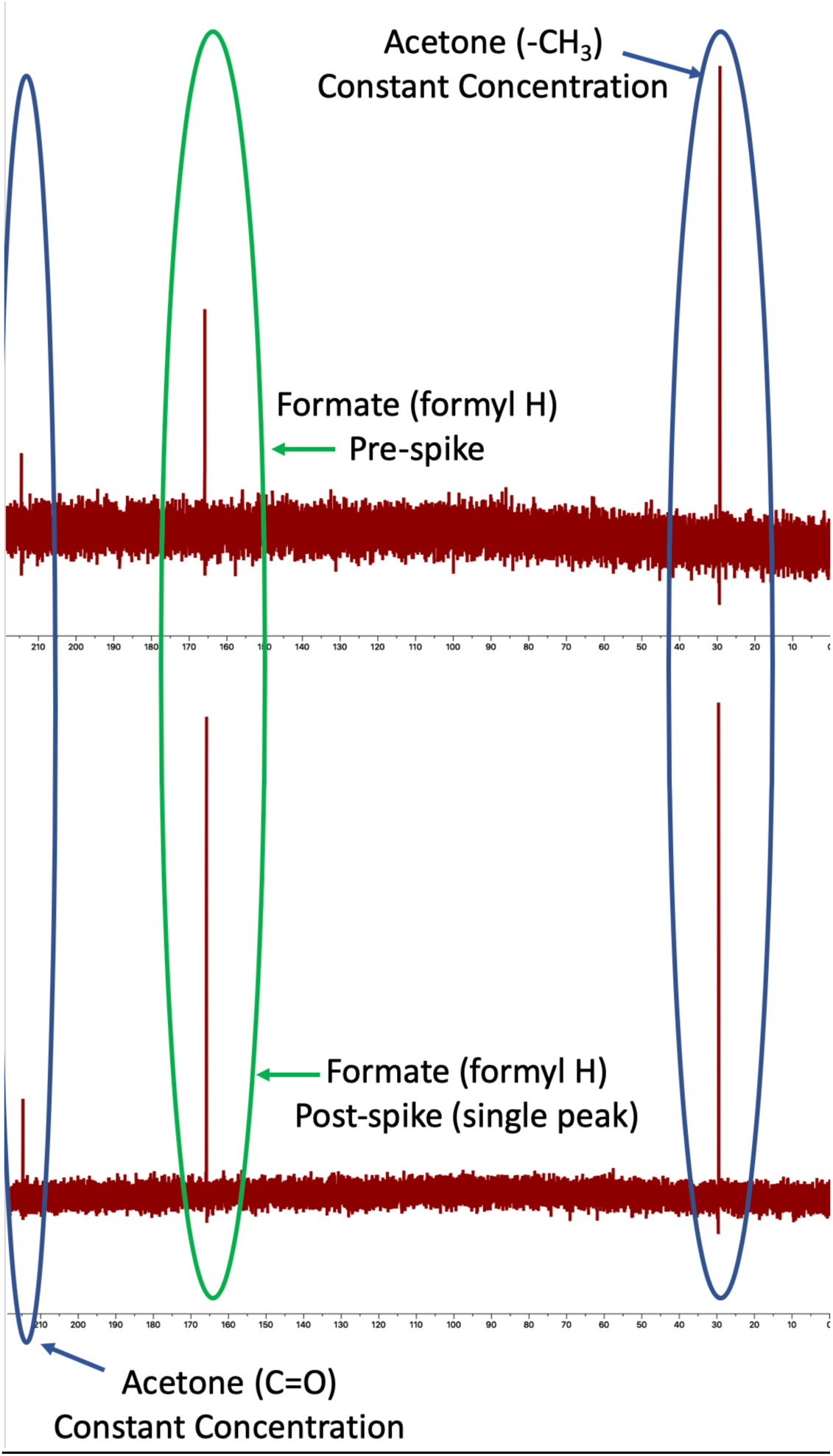
^13^C NMR spectrum of Experiment 1 efflux with and without formate spike. The tubes also contained acetone (CH_3_(CO)CH_3_). **Top:** ^13^C NMR spectrum of experiment 1 efflux. **Bottom:** ^13^C NMR spectrum of experiment 1 efflux after spiking with formate (increasing concentration by 2 µM). The growth of the formate peak (rather than introduction of a new peak) indicates that the original sample indeed contained formate.

### 3. NMR reference spectra

**Figure S3.**
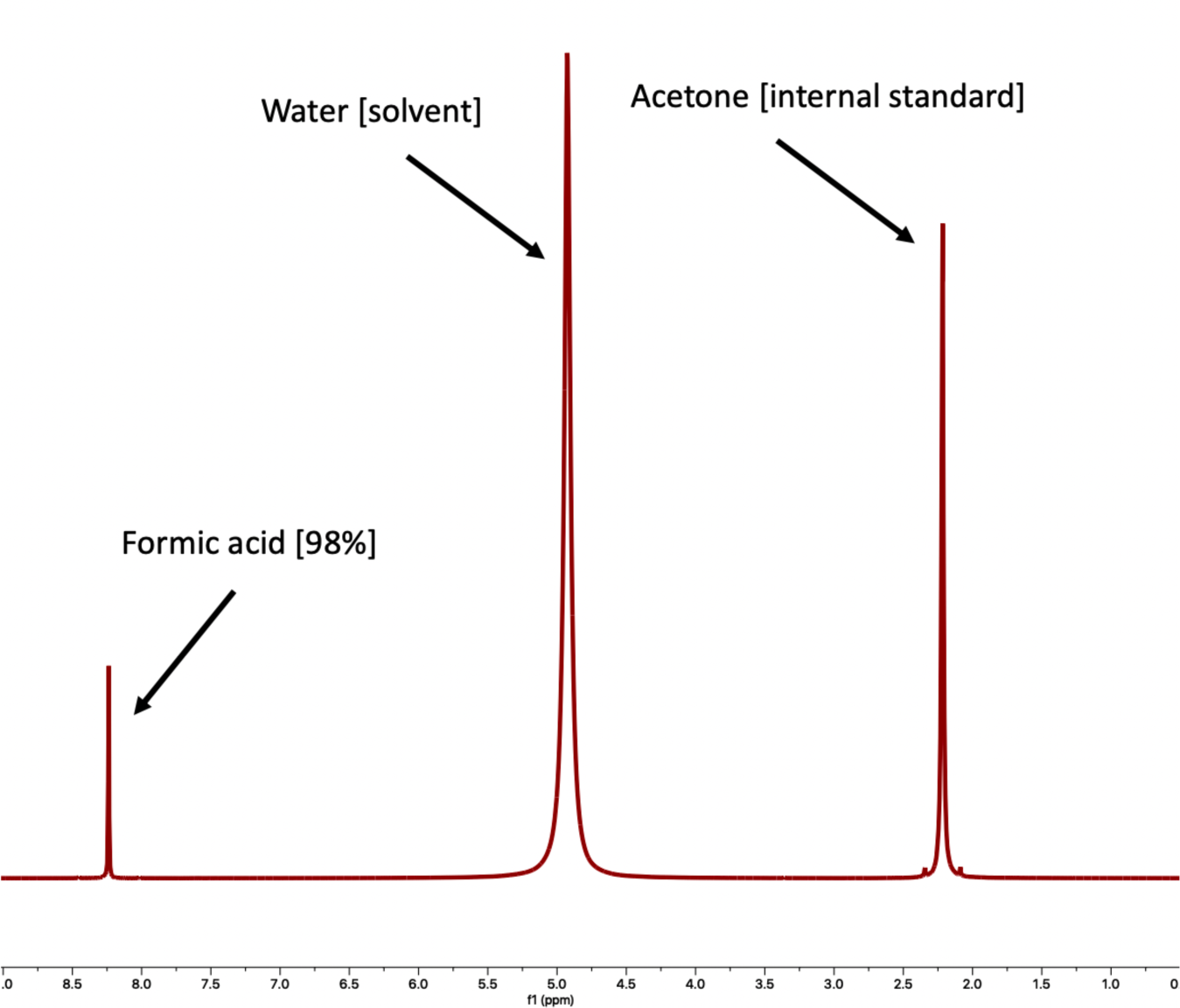
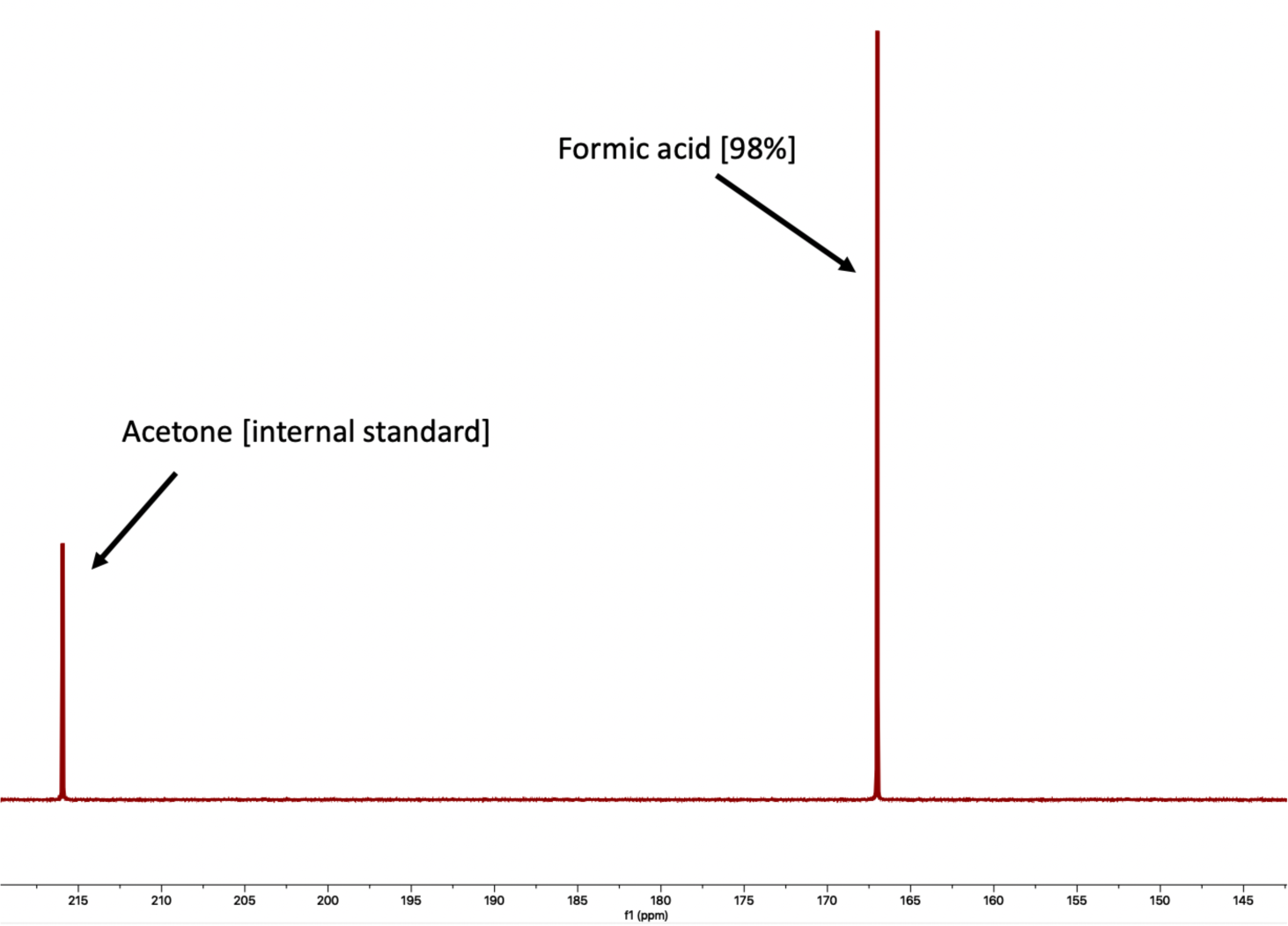
Reference spectra of formic acid and acetone. NMR spectra of formic acid (98%) along with acetone internal standard. **A)** ^1^H NMR spectrum. **B)** ^13^C NMR spectrum.

### 4. Further NMR details and quantification

**Figure S4.**
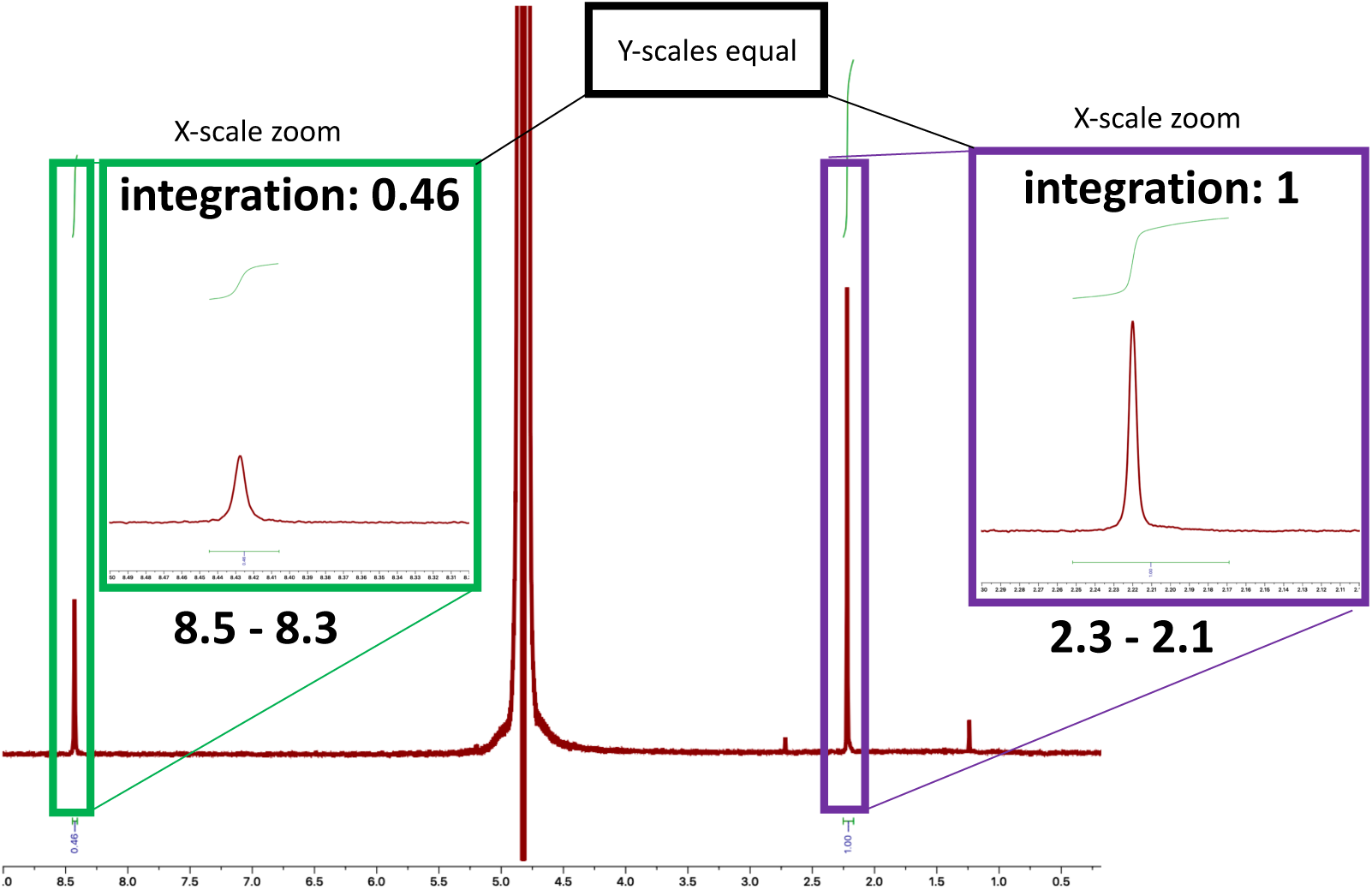
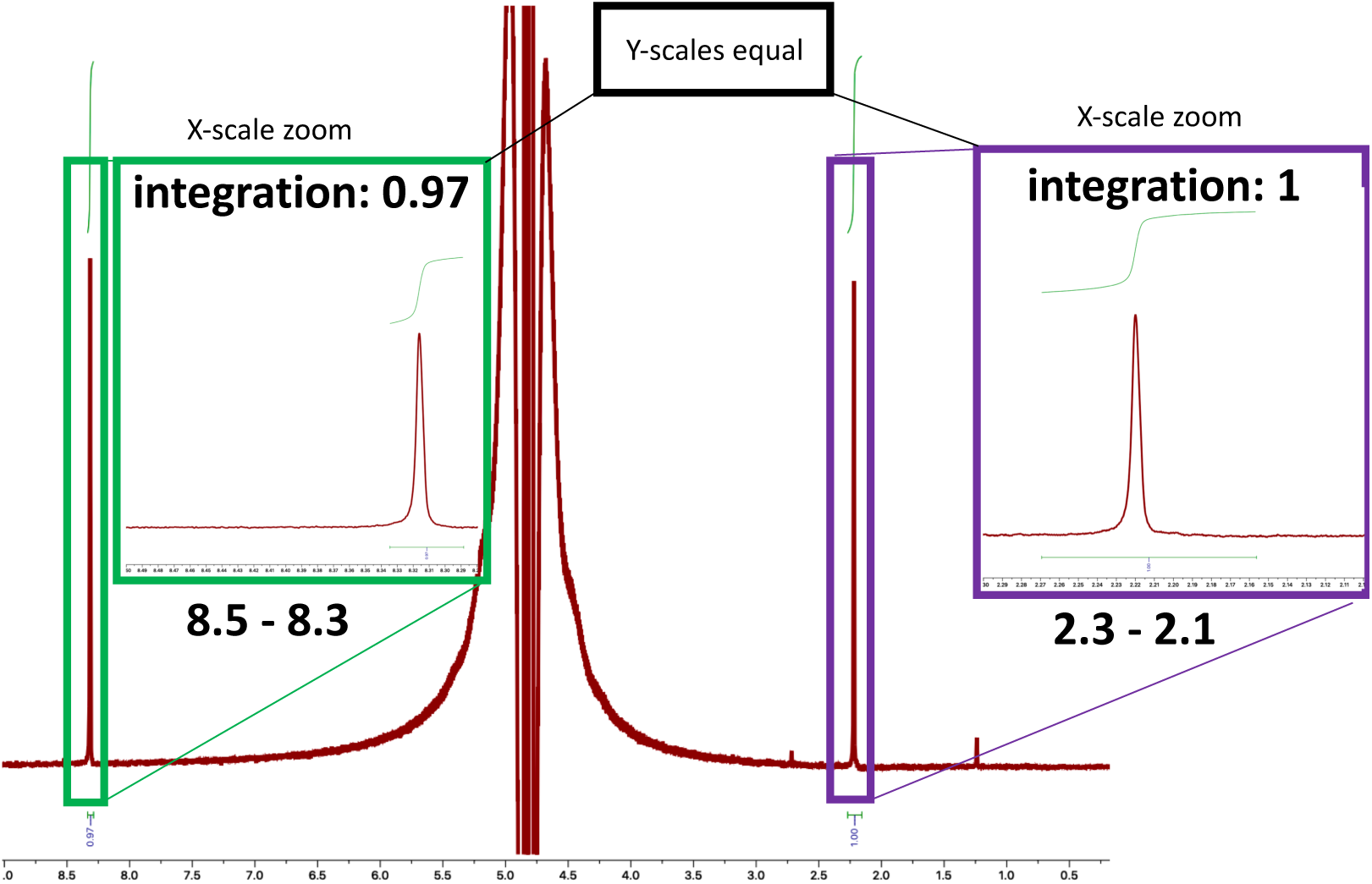
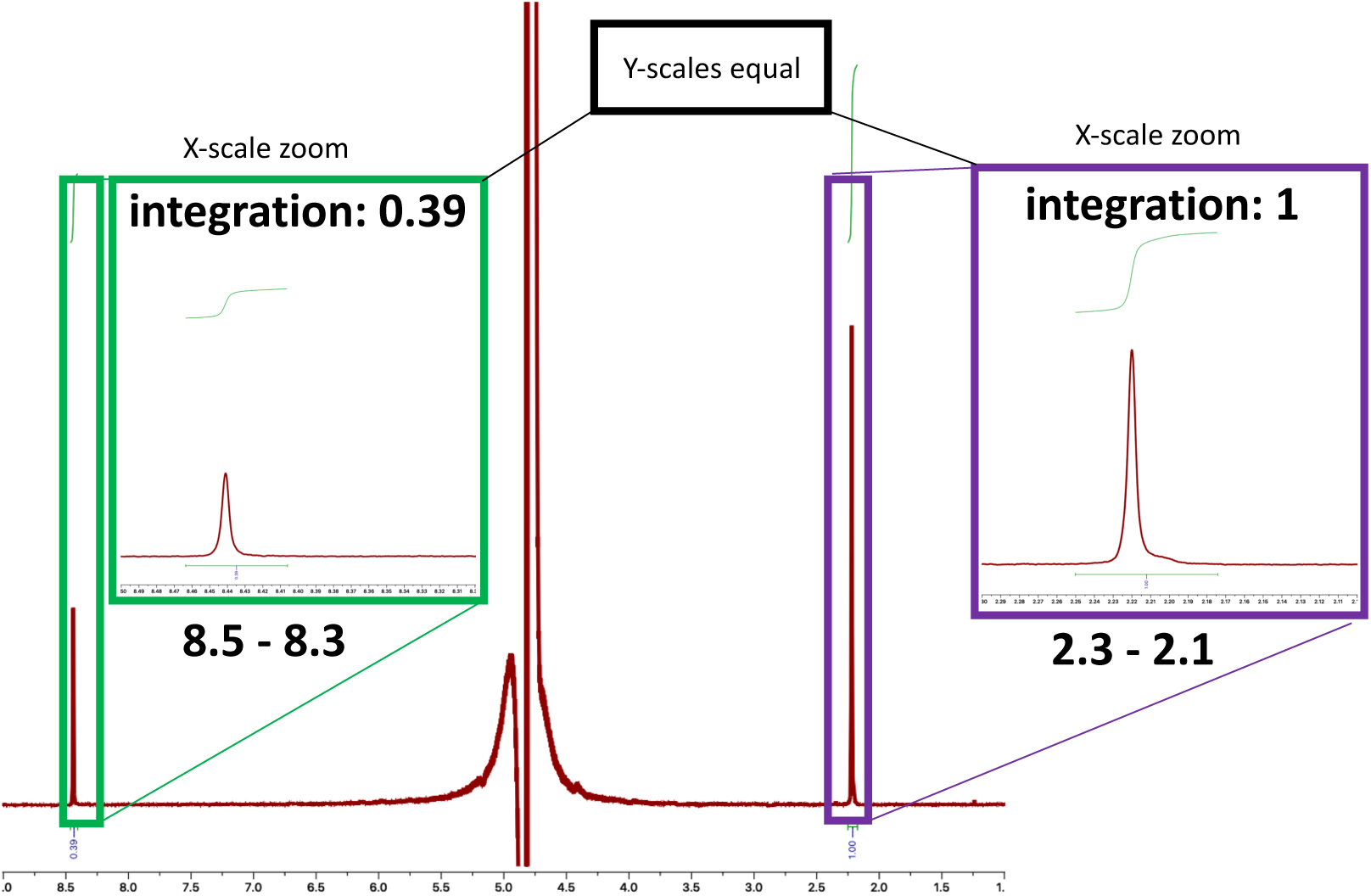
^1^H NMR spectra of Experiment 1 with 0.6 µM acetone as an internal standard. **A)** Integration of the acetone peak (1.00; 6 protons) relative to the formyl peak (0.46; 1 proton), indicates a 2 µM (1.656) concentration of formic acid. **B)** Subsequent spiking of the previous sample (experiment 1) with a standard solution formic acid (raising concentration by an additional 2 µM), doubled the relative integration of the formyl peak (0.97) compared to the acetone peak (1.00). After spiking with a standard solution of formic acid, growth of a single peak, rather than introduction of a new peak, confirms the presence of formic acid in the original sample. Likewise, the magnitude of growth of the formate peak closely corresponds with the originally calculated concentration of formate. Pre-spike calculated formate concentration: 1.7 µM. Post-spike (2.0 µM) calculated formate concentration: 3.5 µM. Therefore, using the post/pre spike values to back-calculate the concentration of formate offers a value of 1.5 µM (3.5 – 2.0 = 1.5 µM). **C)** Integration of the acetone peak (1.00; 6 protons) relative to the formyl peak (0.39; 1 proton), indicates a 1.404 µM concentration of formic acid. An average of the two experiment 1 samples indicates a 1.5 μM concentration of formic acid.

**Figure S5.**
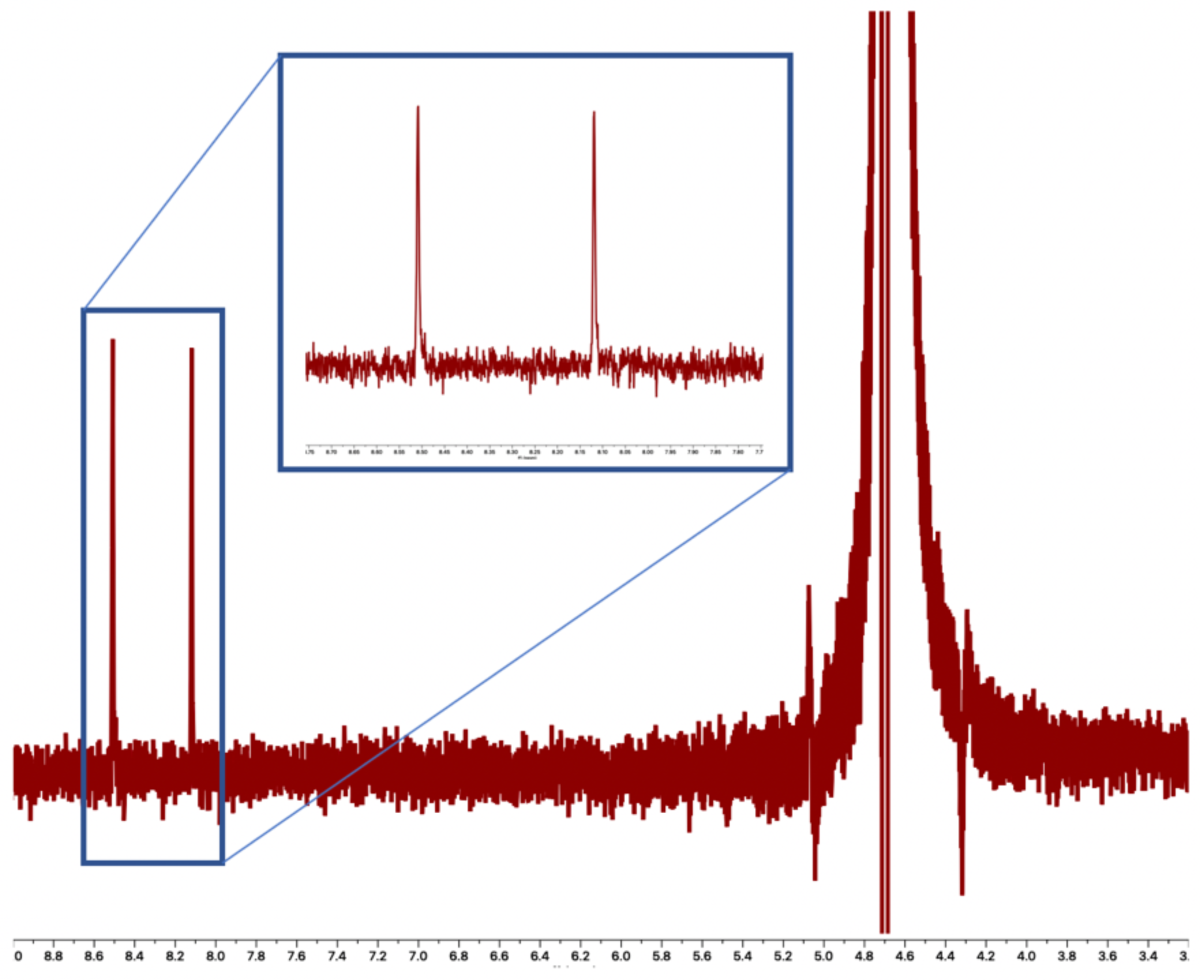
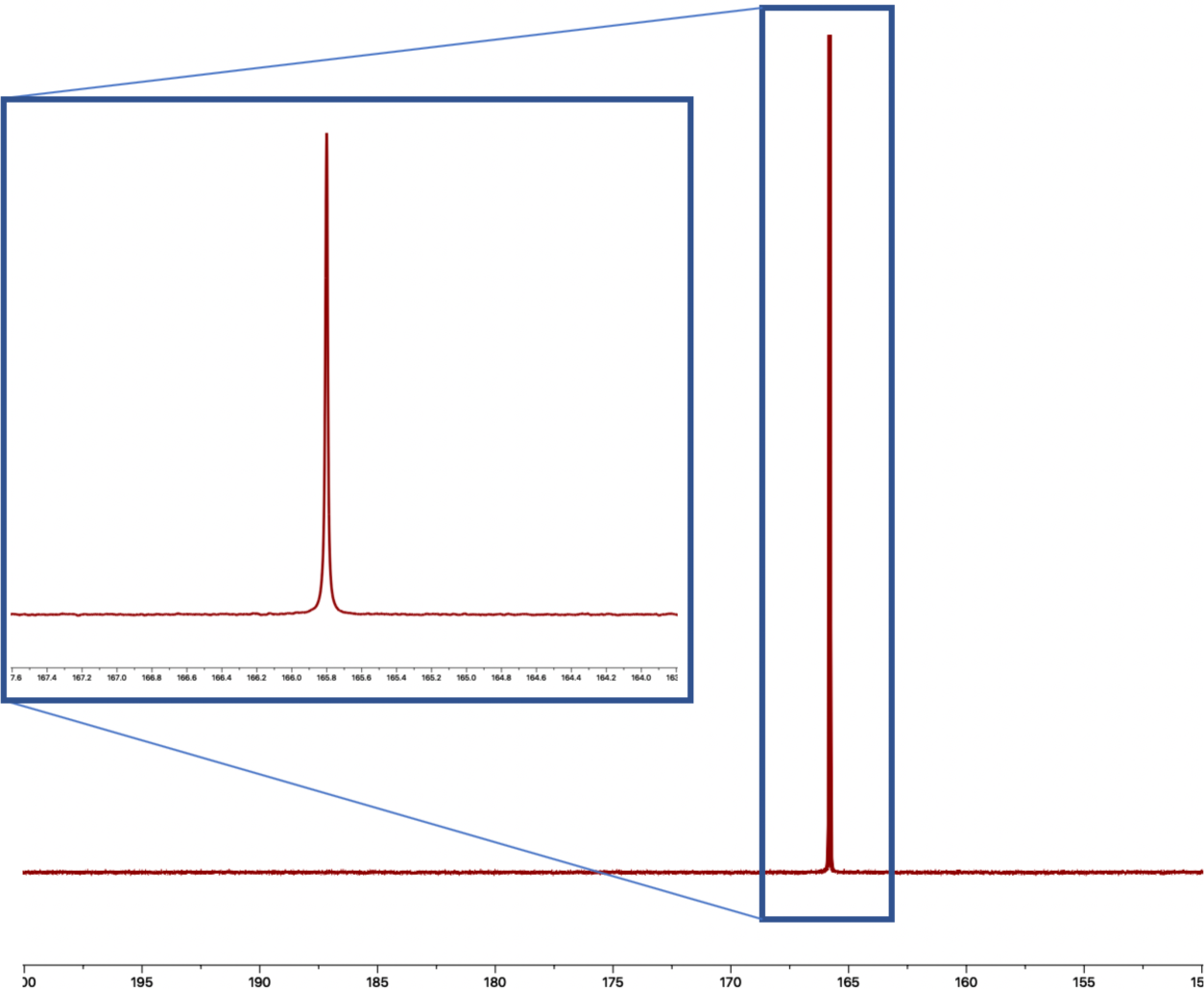
^1^H and ^13^C NMR spectra of Experiment 2 (^13^CO_2_ as ocean-driving gas) **A)** ^1^H NMR indicates splitting for formyl peak into a doublet. In the rest of our study, quantification of formate was conducted by comparing the integration of the formyl singlet against the integration of an acetone internal standard. Because the ^13^C labeling splits the formyl singlet into a doublet, such a comparison would be unreliable in this case. This analysis therefore offers only identification of the H^13^COO^−^ product, but not quantification. **B)** ^13^C NMR spectrum.

**Figure S6.**
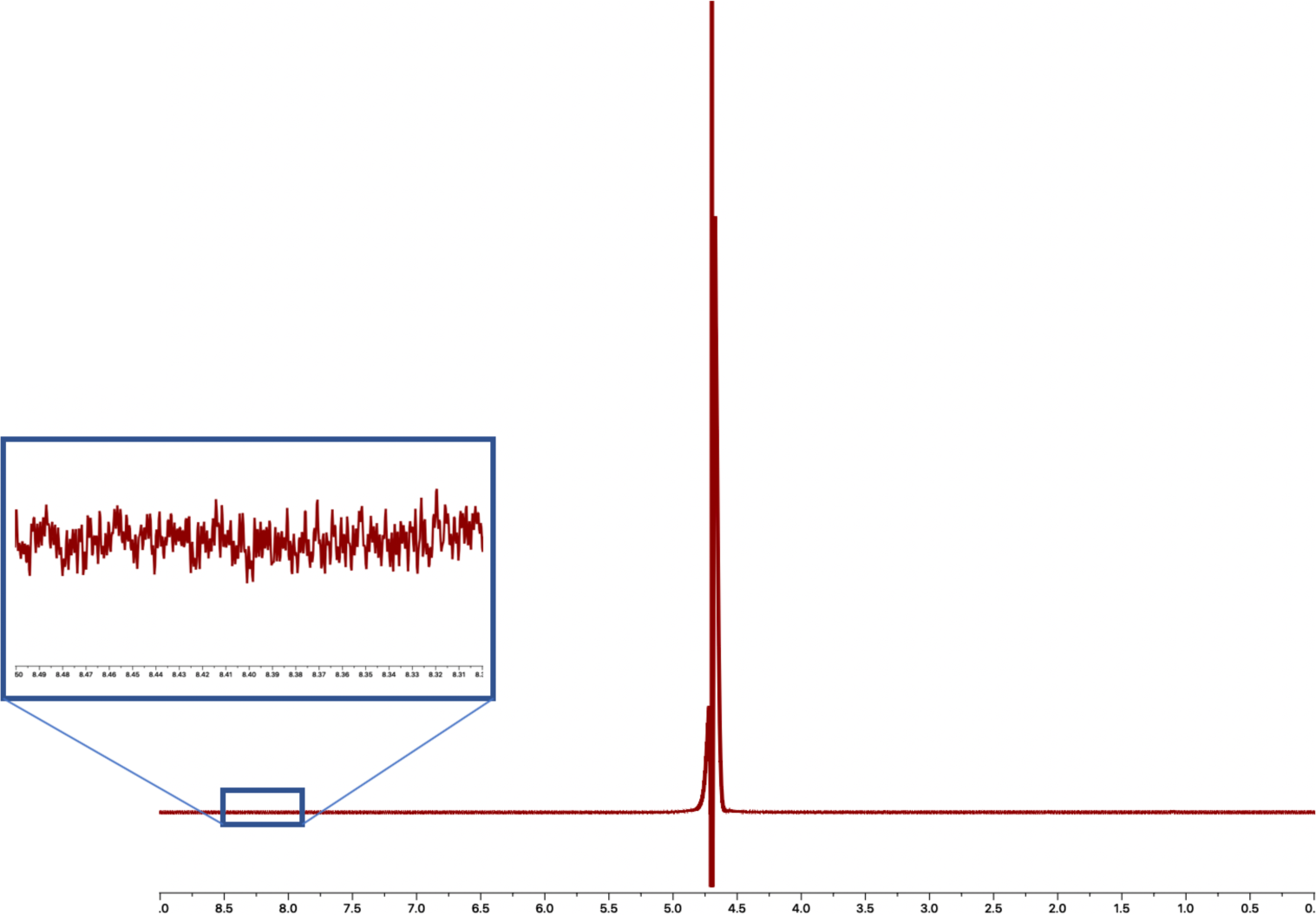
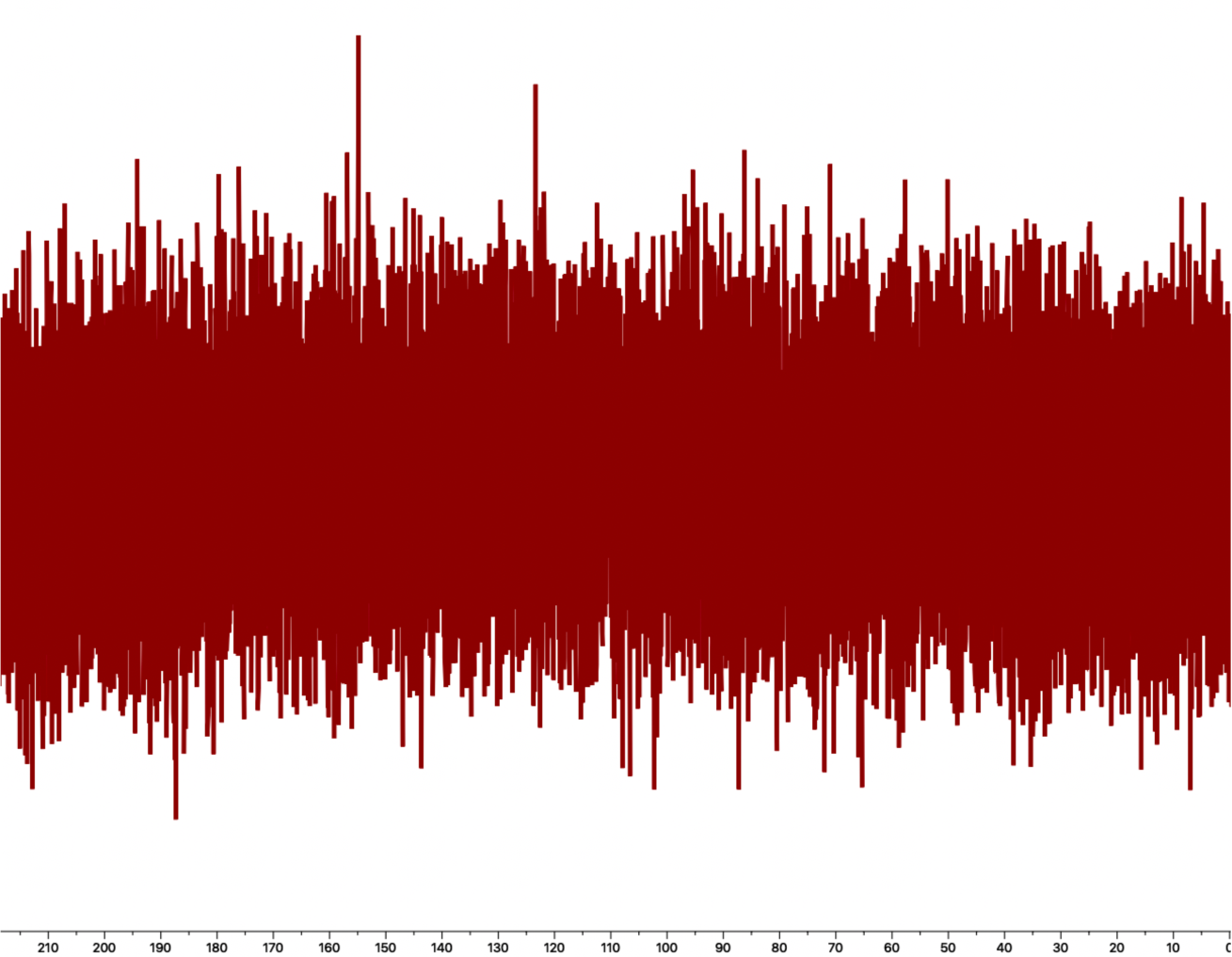
^1^H and ^13^C NMR spectra of Experiment 3 (N_2_ as vent-driving gas) **A)** The absence of any relevant peak (other than water) in the ^1^H NMR spectrum indicates that no product was formed. **B)** No relevant peak in the ^13^C NMR spectrum indicates that no product was formed appreciably.

**Figure S7.**
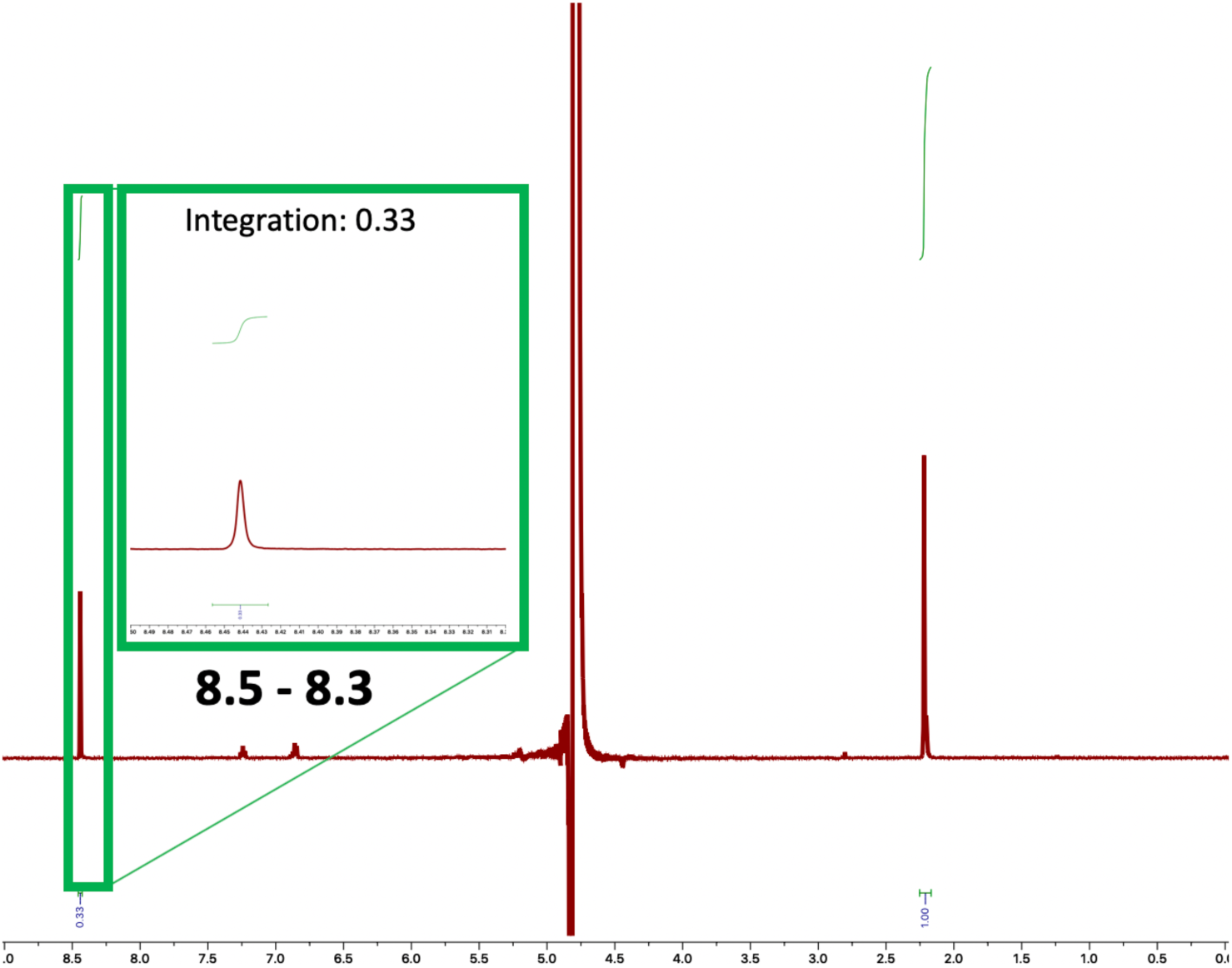
^1^H spectrum of Experiment 4 (D_2_ as vent-driving gas) The formyl singlet indicates the presence of HCOO^−^ rather than DCOO^−^ (which would offer a doublet). Integration of the acetone peak (1.00; 6 protons) relative to the formyl peak (0.33; 1 proton), indicates a 1.2 µM concentration of formic acid.

**Figure S8.**
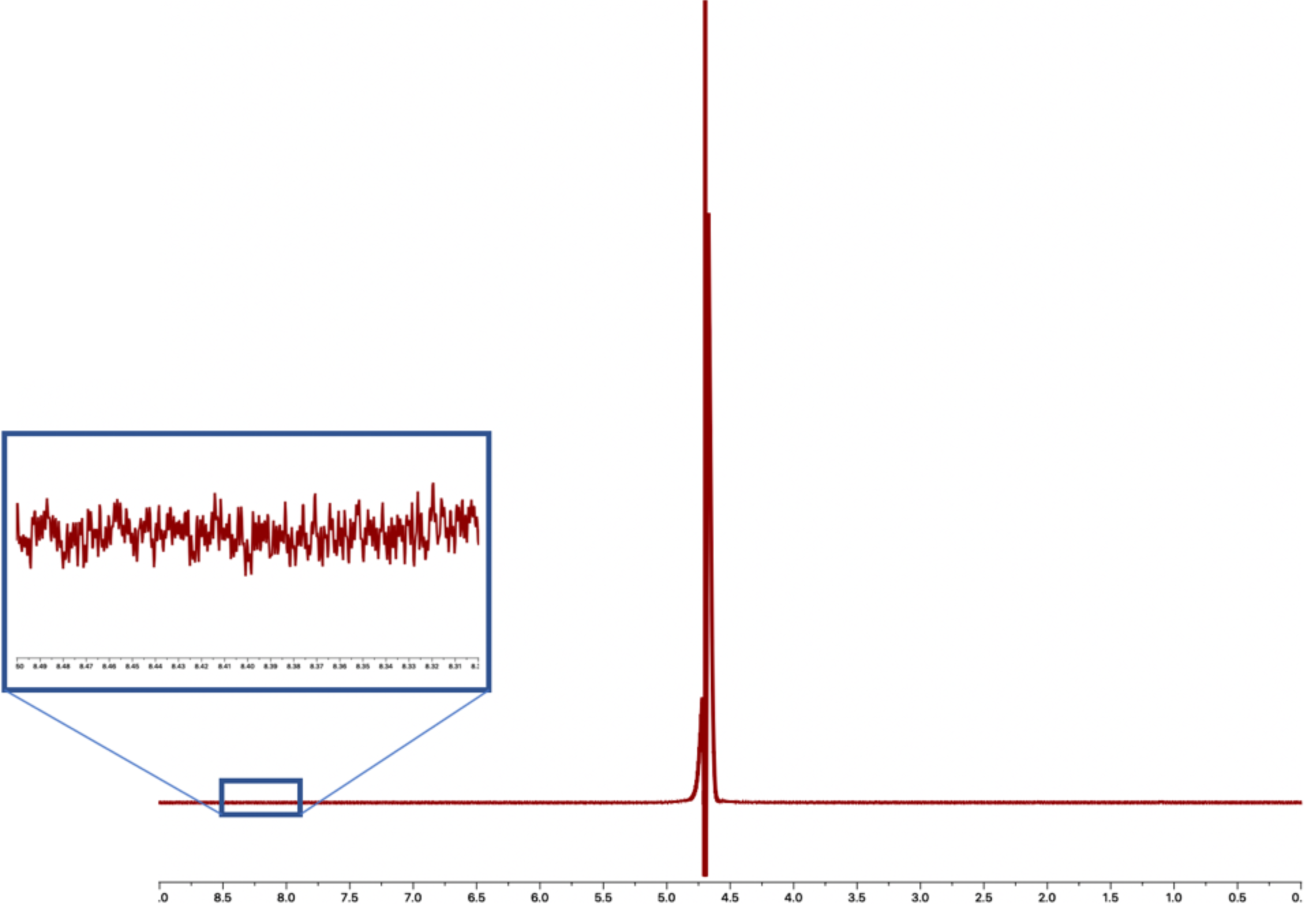
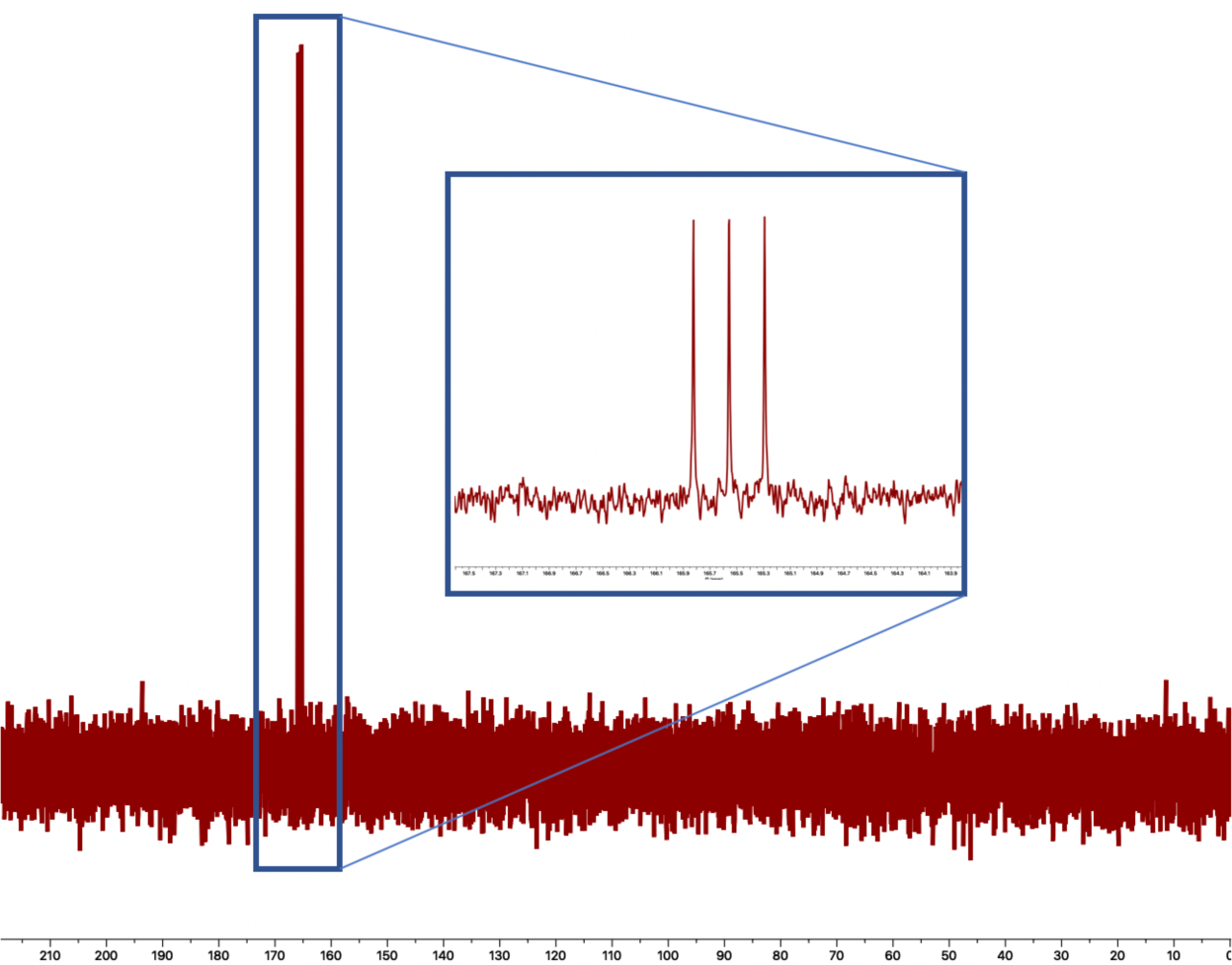
^1^H and ^13^C NMR spectra of Experiment 5 (D_2_O as ocean solvent) **A)** ^1^H NMR indicates no formyl ^1^H, which could mean either that no formate was produced at all, or instead, that the product is deuterium-labeled formate (DCOO^−^)—as is indeed shown by the ^13^C spectrum in panel B). In the rest of our study, quantification of formate was conducted by comparing the integration of the formyl singlet against the integration of an acetone internal standard. Because there is no formate peak, such quantification is not possible in this case. This analysis therefore offers only identification of the DCOO^−^ product, but not quantification. **B)** ^13^C NMR spectrum indicates splitting of the singlet into a triplet—consistent with D-C coupling.

**Figure S9.**
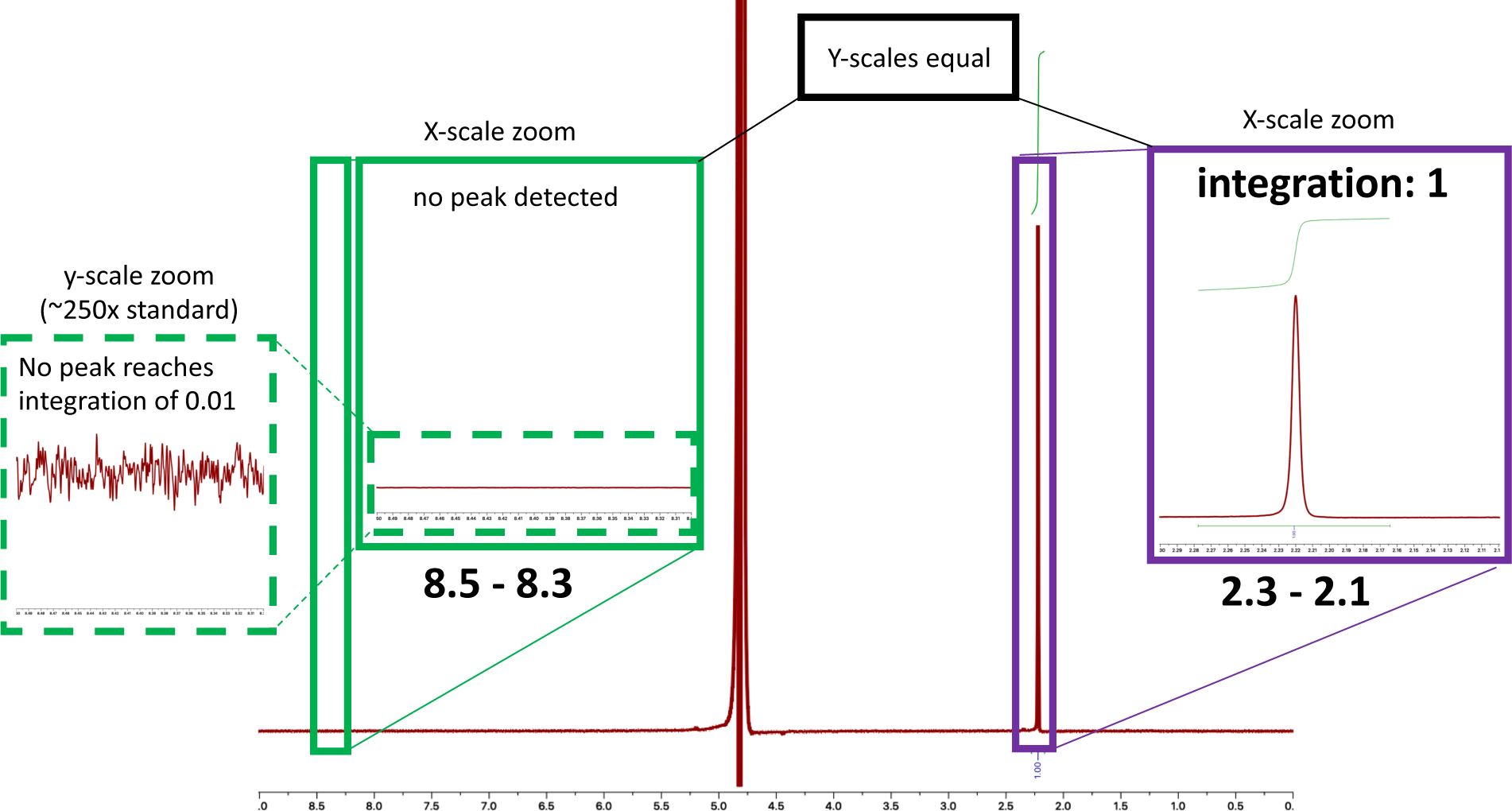
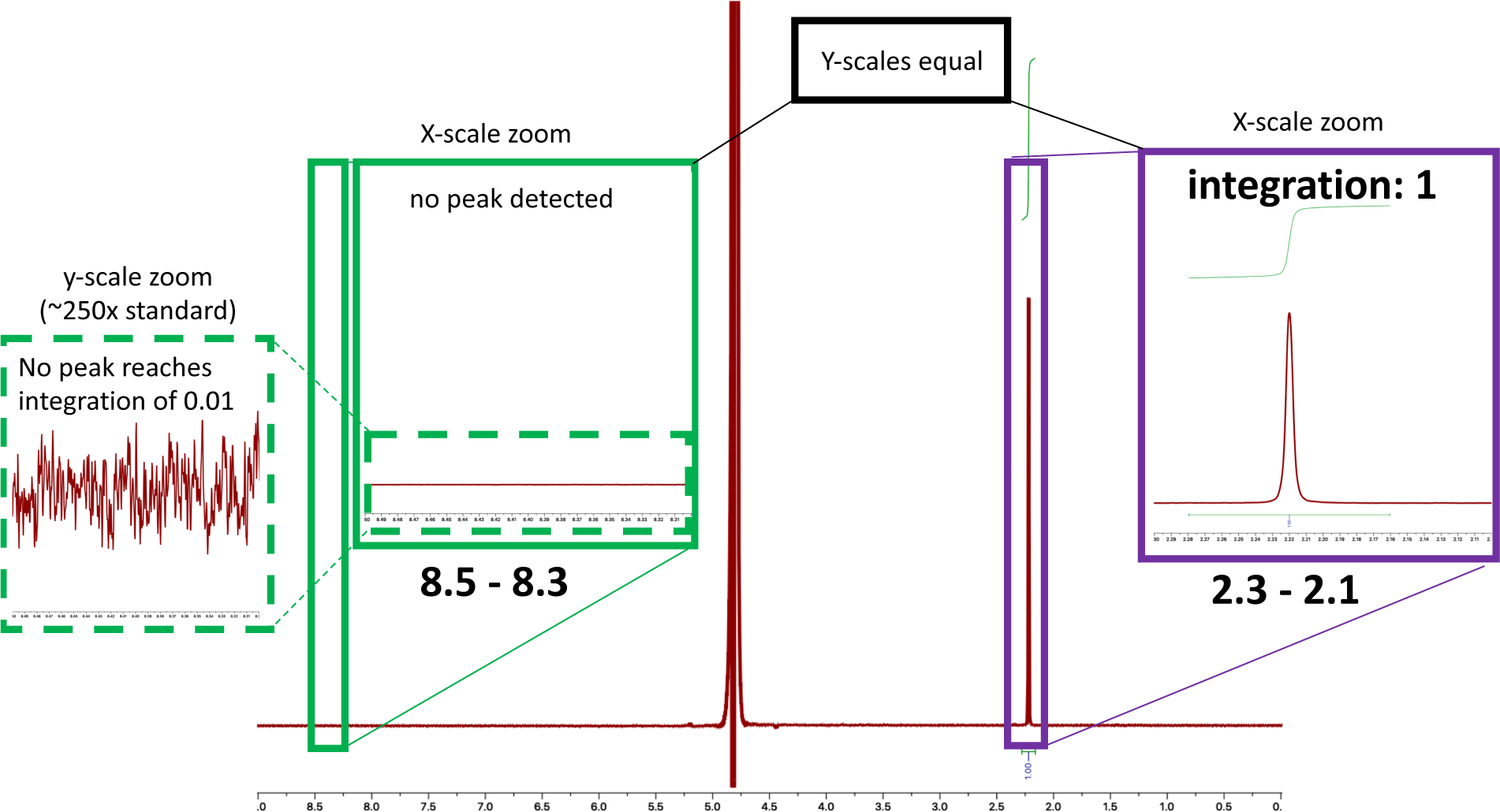
^1^H NMR spectra of Experiment 6 (no solutes on vent-side) Acetone 0.6 µM was added as an internal standard. **A)** No formate peak was detected. **B)** Duplicate. No formate peak was detected.

**Figure S10.**
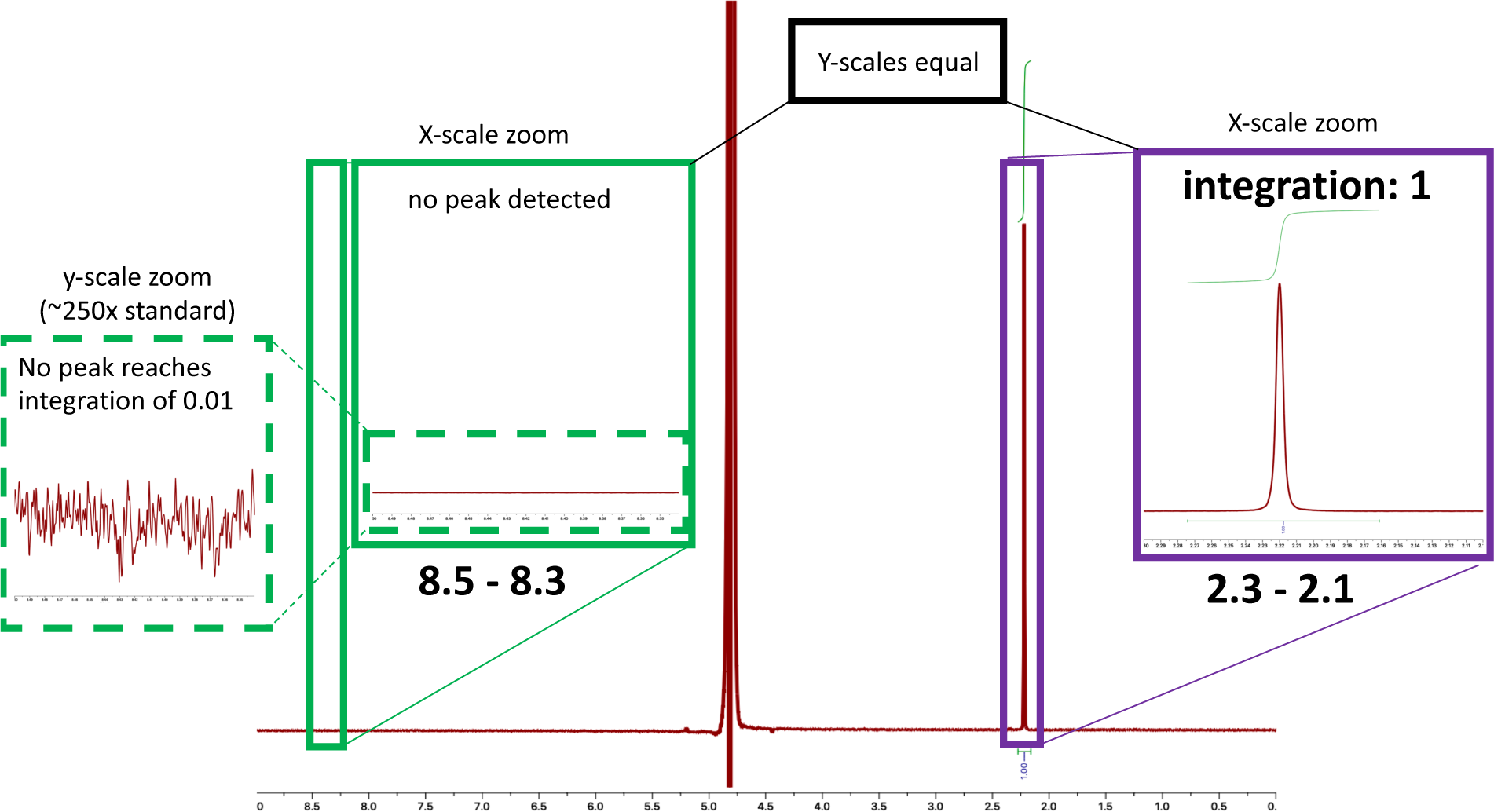
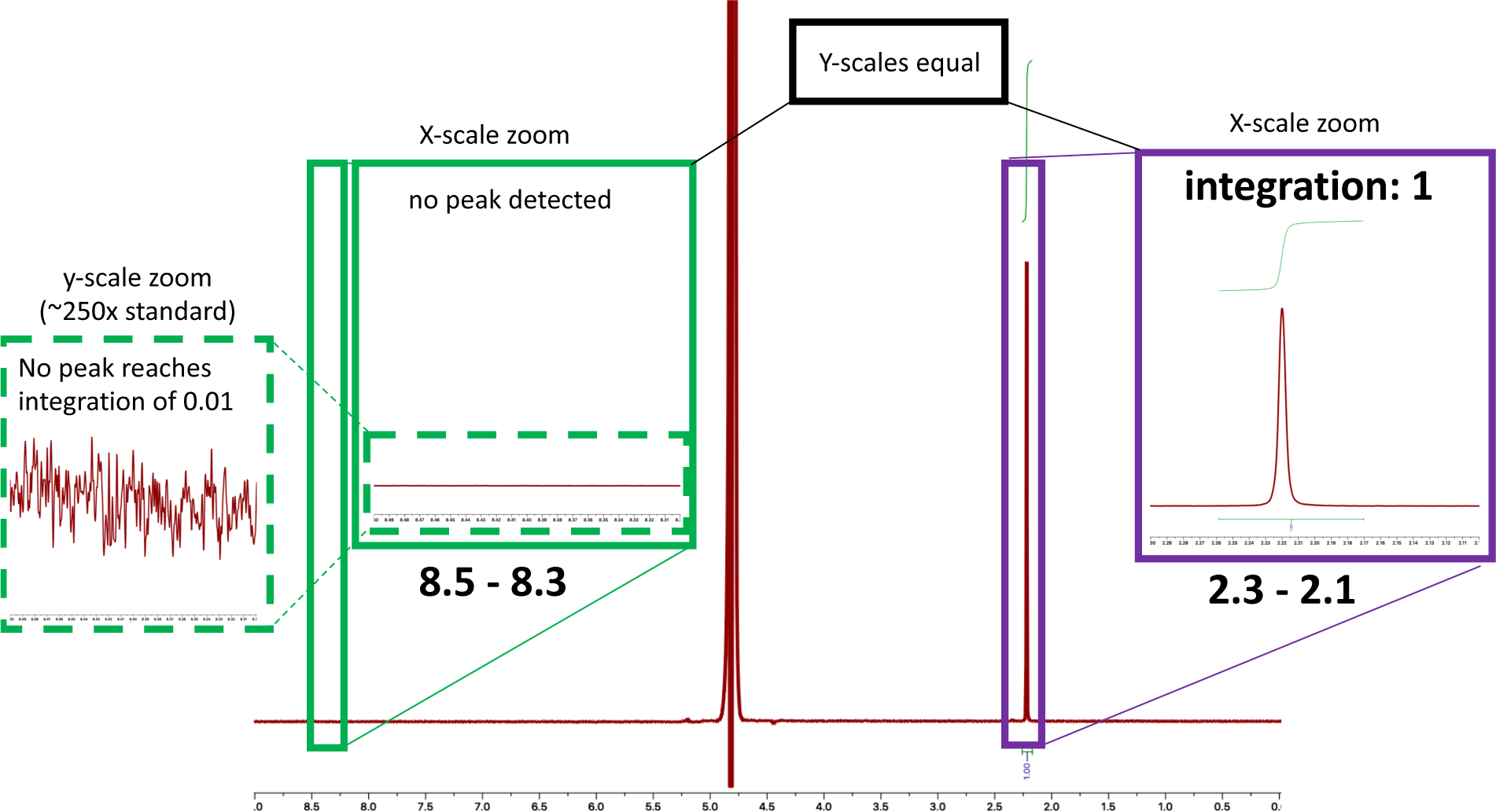
^1^H NMR spectra of Experiment 7 (vent-side titrated to pH 7.0) Acetone 0.6 µM was added as an internal standard. **A)** No formate peak was detected. **B)** Duplicate. No formate peak was detected.

**Figure S11.**
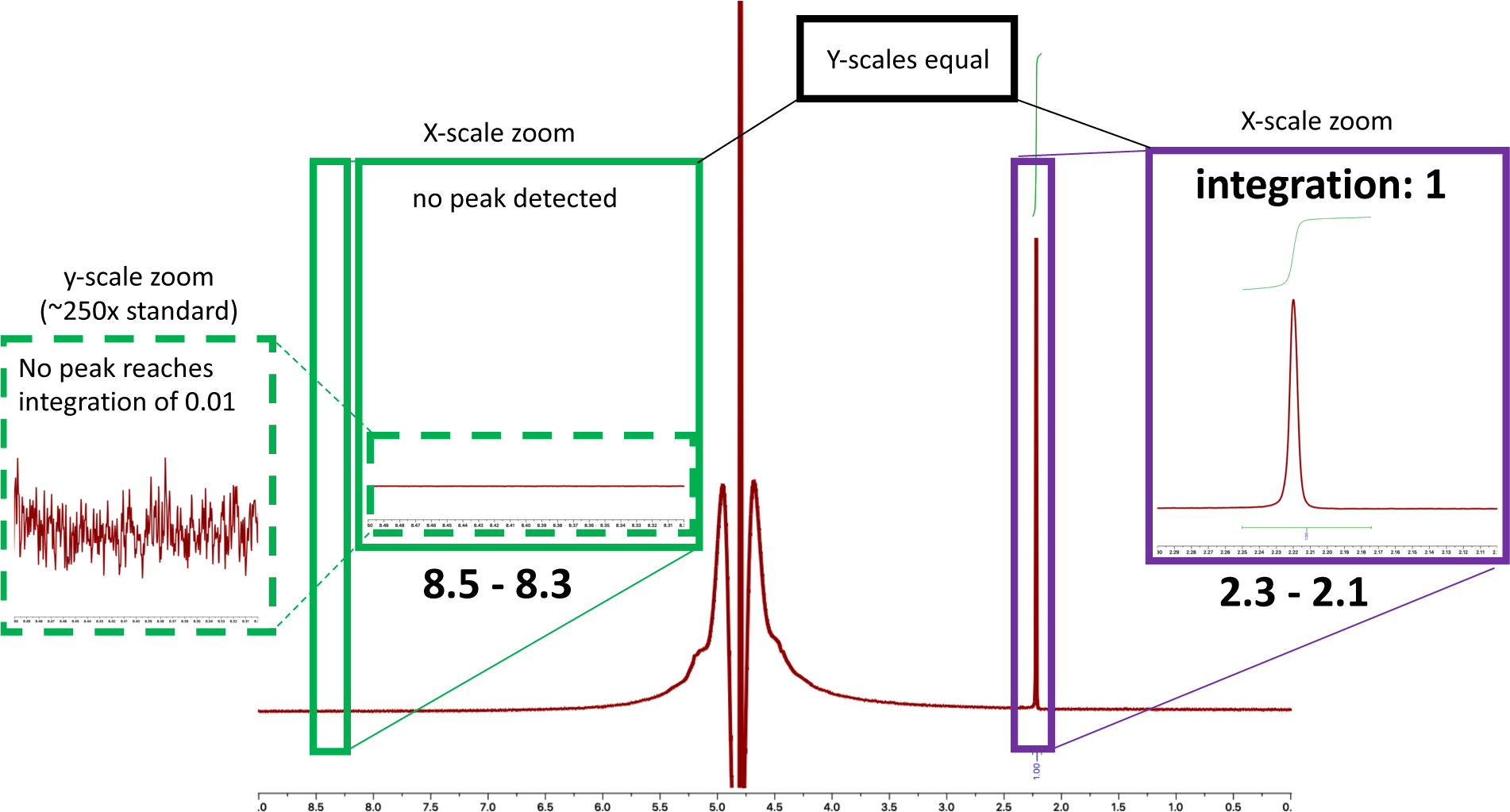
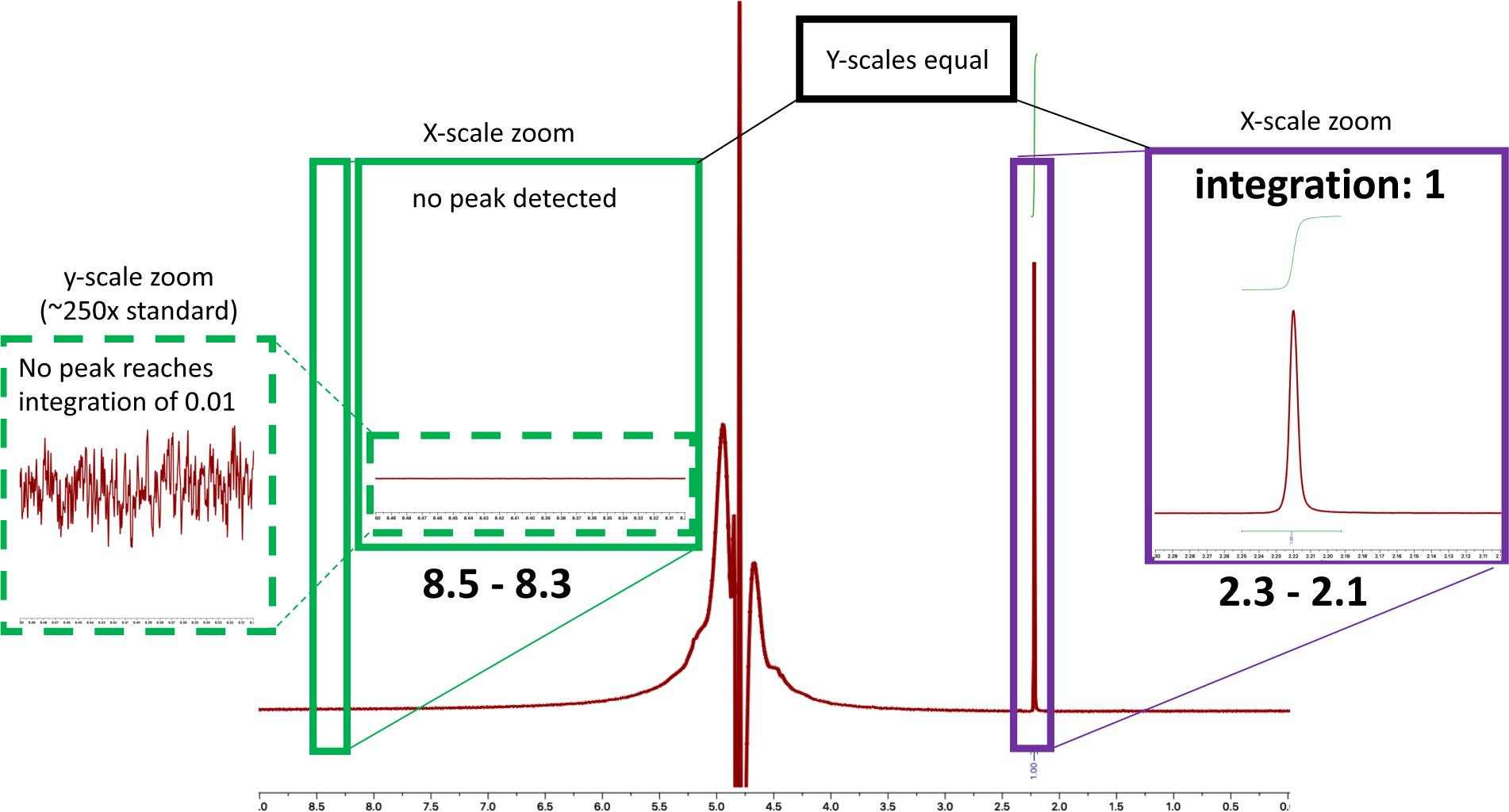
^1^H NMR spectra of Experiment 8 (vent-side titrated to pH 3.9) Acetone 0.6 µM was added as an internal standard. **A)** No formate peak was detected. **B)** Duplicate. No formate peak was detected.

**Figure S12.**
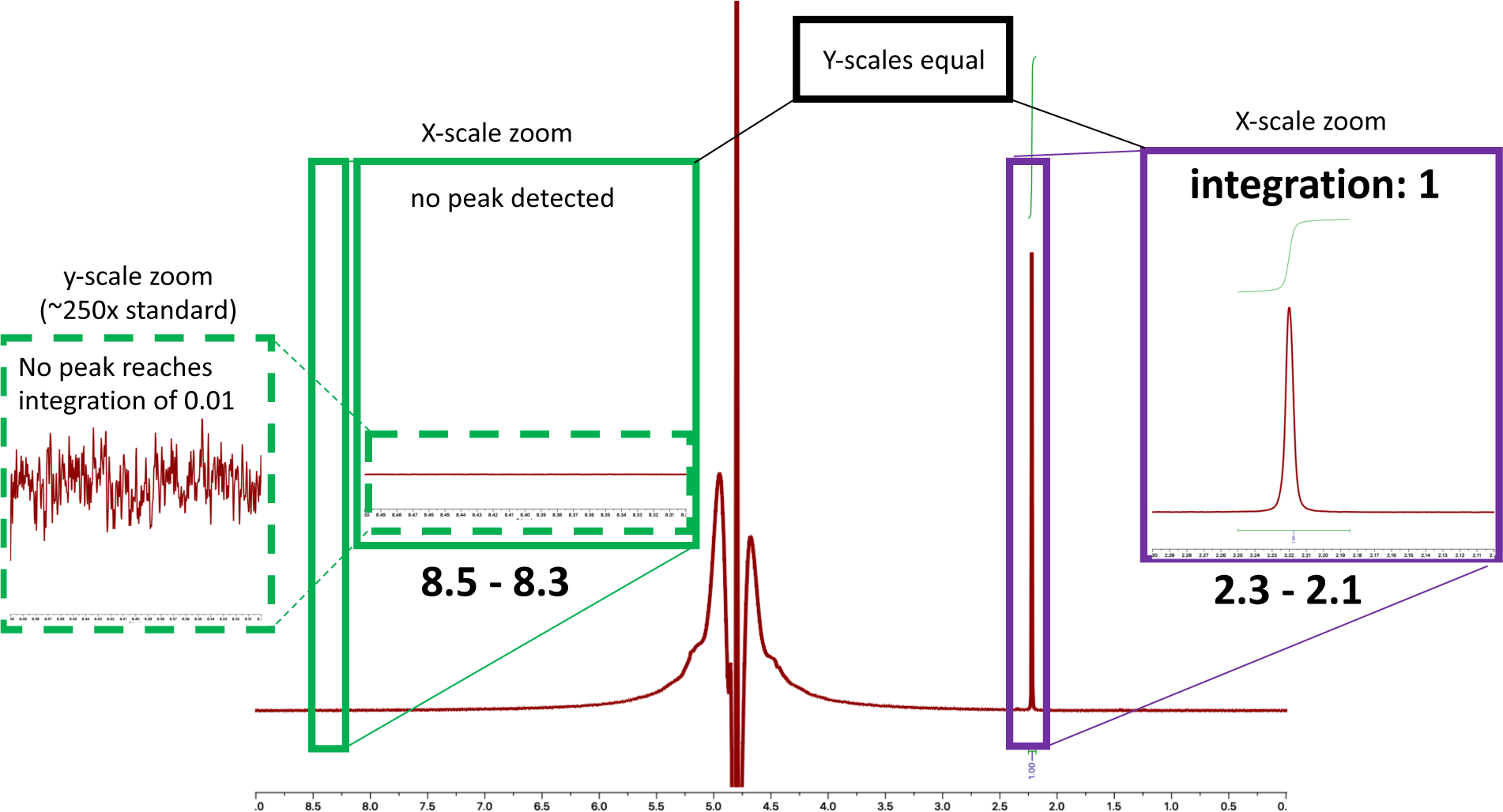
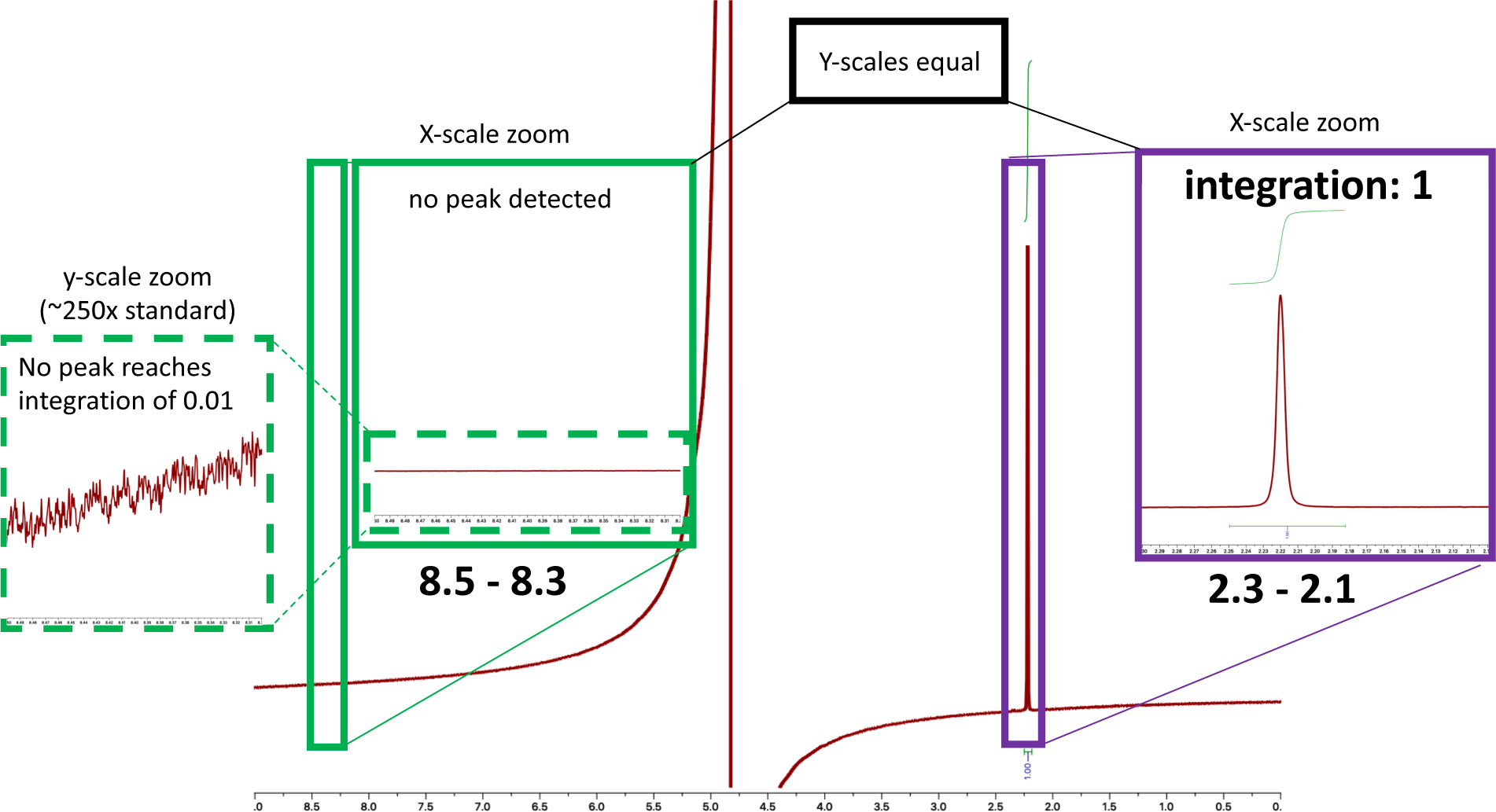
^1^H NMR spectra of Experiment 9 (Na_2_CO_3_ in ocean fluid) Acetone 0.6 µM was added as an internal standard. **A)** No formate peak was detected. **B)** Duplicate. No formate peak was detected.

**Figure S13.**
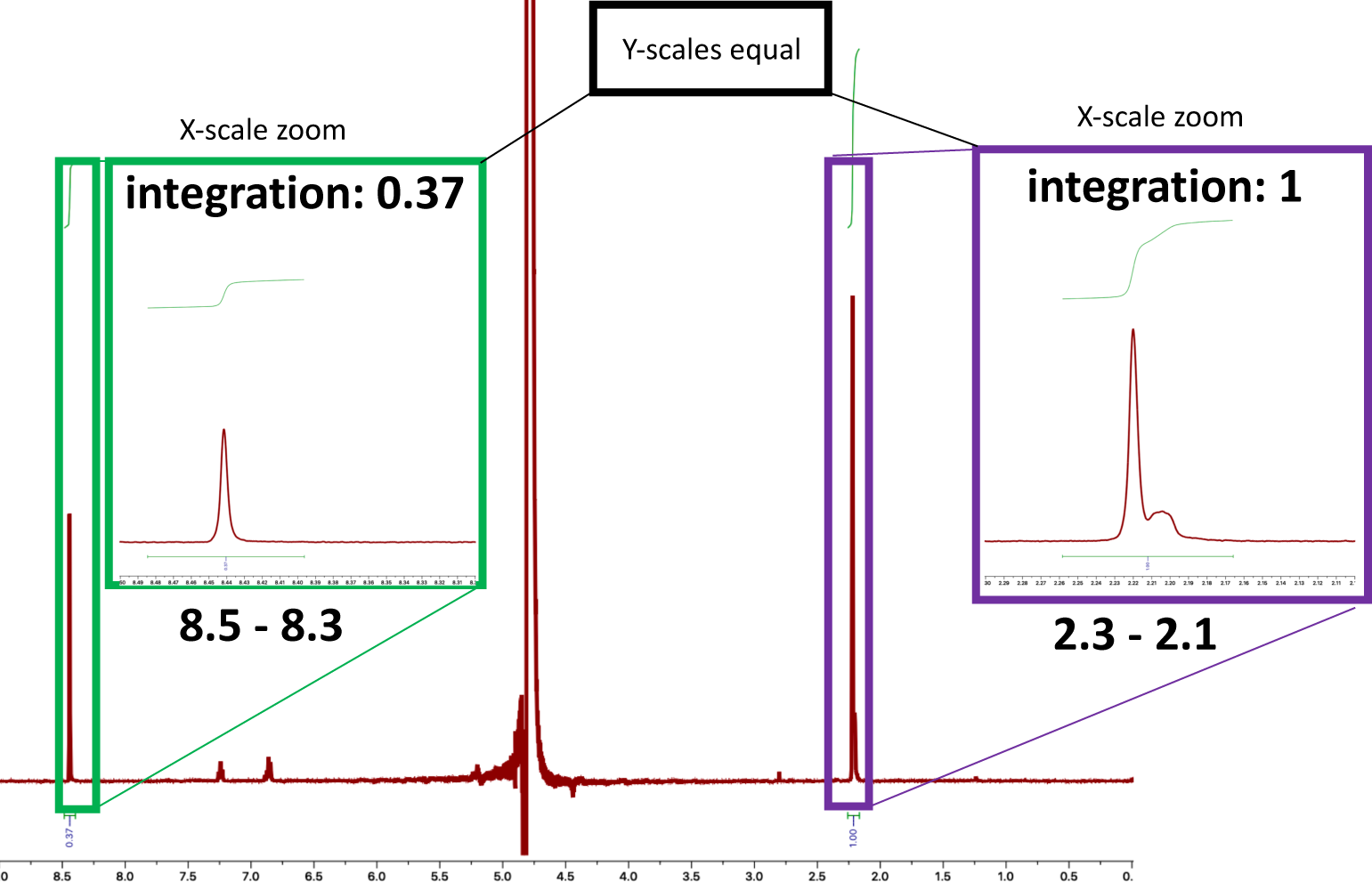
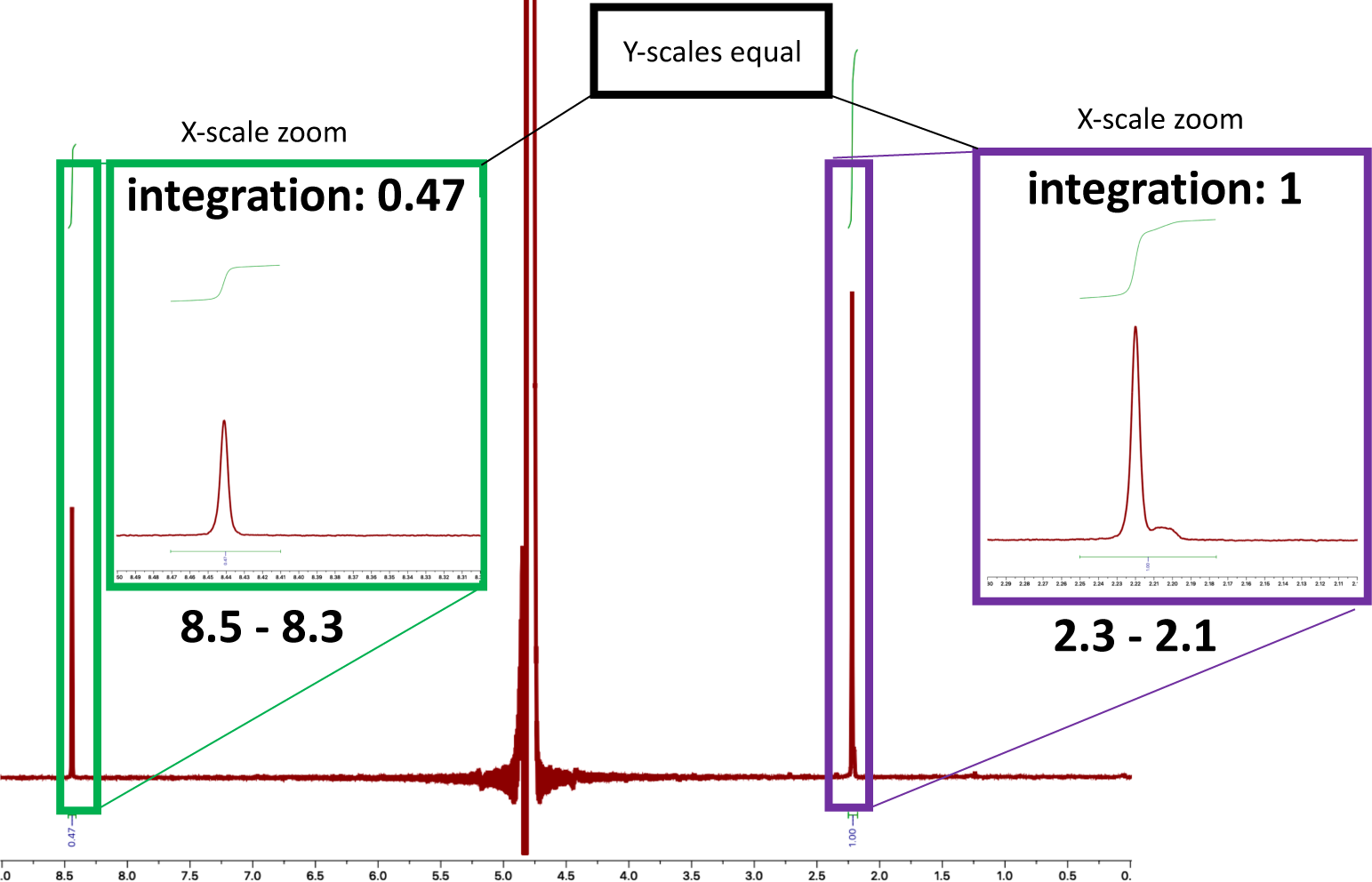
^1^H NMR spectra of Experiment 10 (no Na2Si3O7 in vent post-precipitation fluid) Acetone 0.6 µM was added as an internal standard. **A)** Integration of the acetone peak (1.00; 6 protons) relative to the formyl peak (0.37; 1 proton), indicates a 1.332 µM concentration of formic acid. **B)** Duplicate. Integration of the acetone peak (1.00; 6 protons) relative to the formyl peak (0.47; 1 proton), indicates a 1.692 µM concentration of formic acid. An average of the two samples indicates a 1.5 µM concentration of formic acid. A small broad peak in the alkyl region (2.19–2.21 ppm), observed also in Figure S14, will require further analysis.

**Figure S14.**
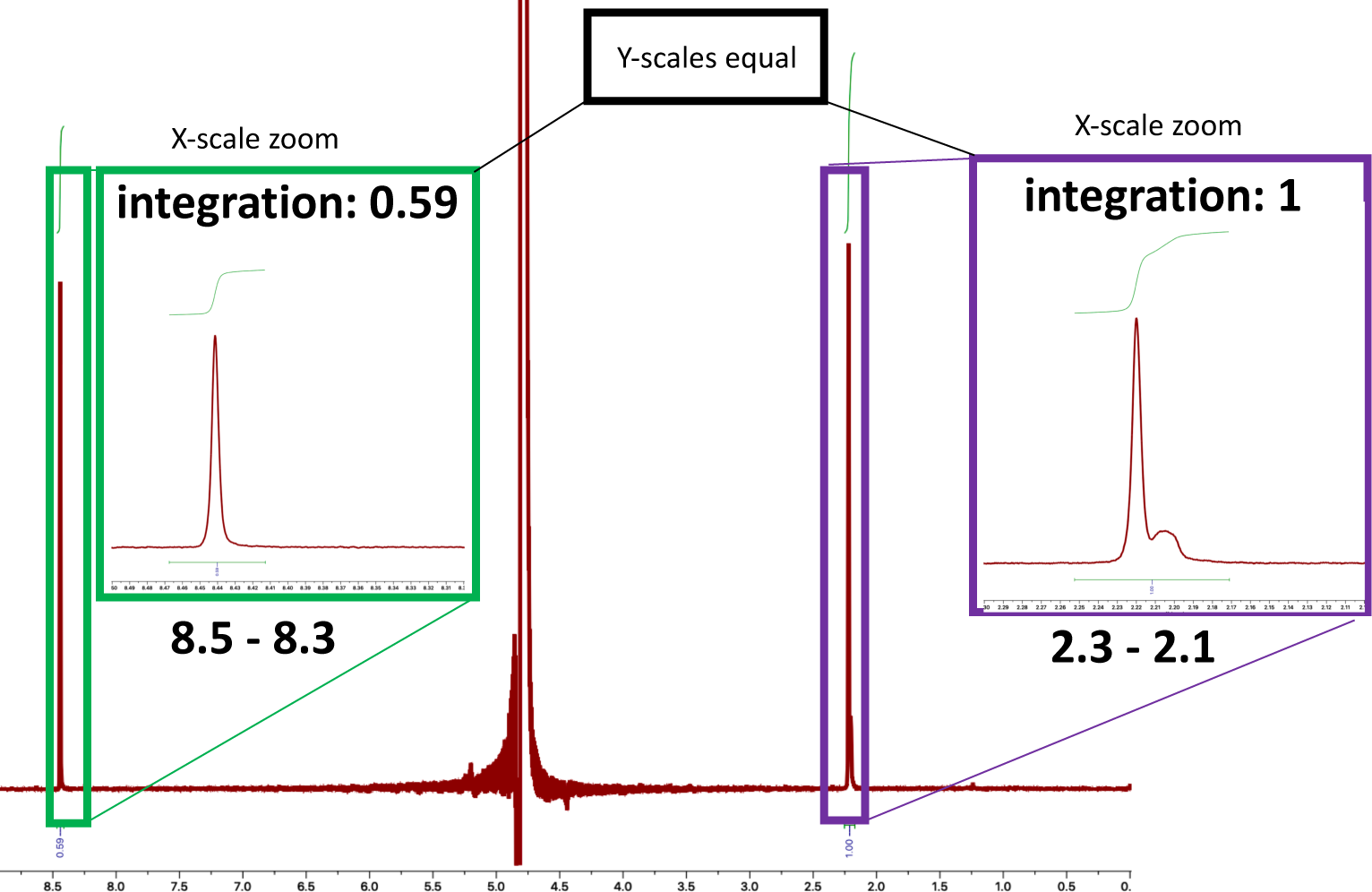
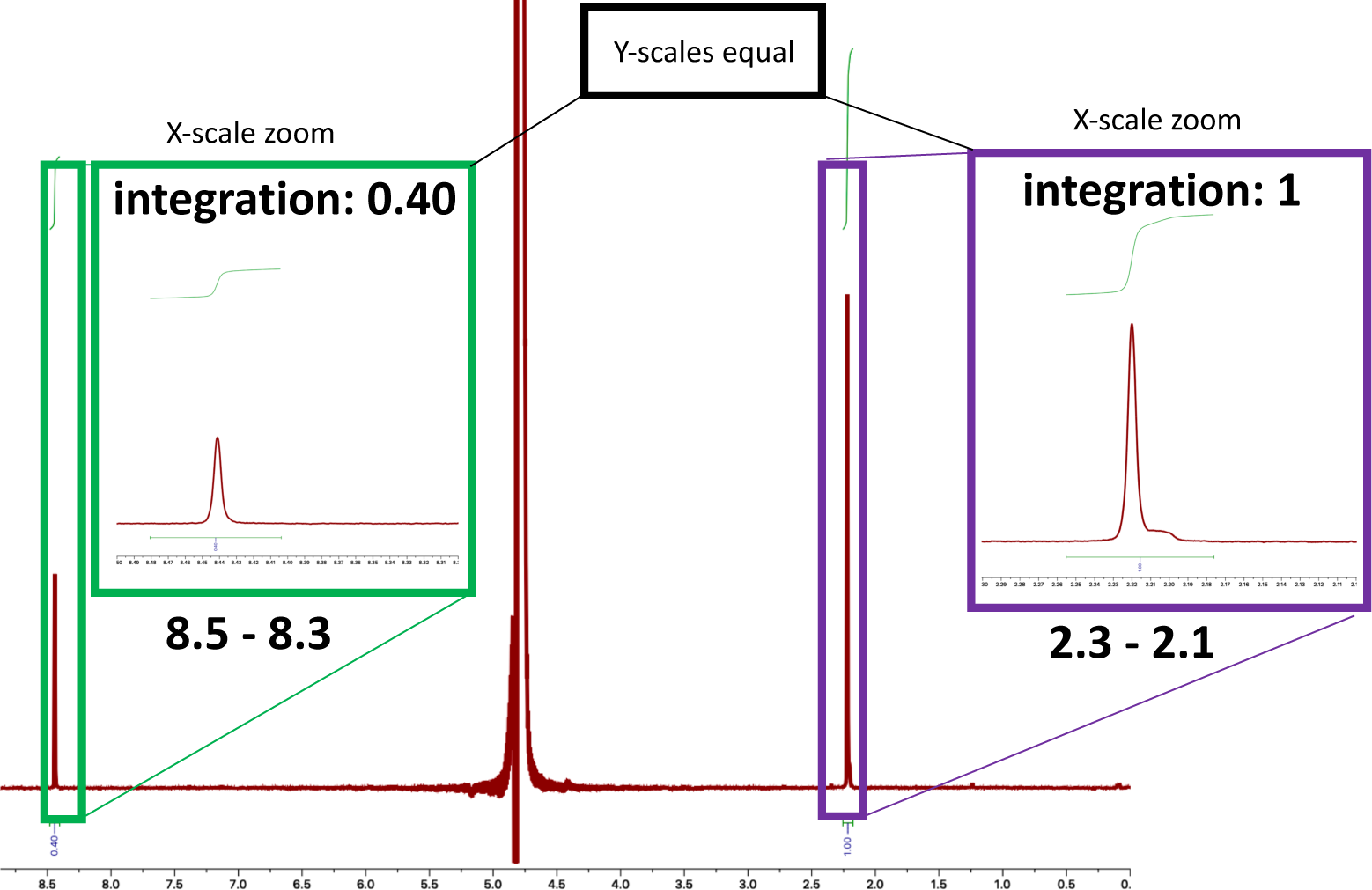
^1^H NMR spectra of Experiment 11 (only Na2S in vent post-precipitation fluid) Acetone 0.6 µM was added as an internal standard. **A)** Integration of the acetone peak (1.00; 6 protons) relative to the formyl peak (0.59; 1 proton), indicates a 2.124 µM concentration of formic acid. **B)** Duplicate. Integration of the acetone peak (1.00; 6 protons) relative to the formyl peak (0.40; 1 proton), indicates a 1.440 µM concentration of formic acid. An average of the two samples indicates a 1.8 µM concentration of formic acid. A small broad peak in the alkyl region (2.19–2.21 ppm), observed also in Figure S13, will require further analysis.

**Figure S15.**
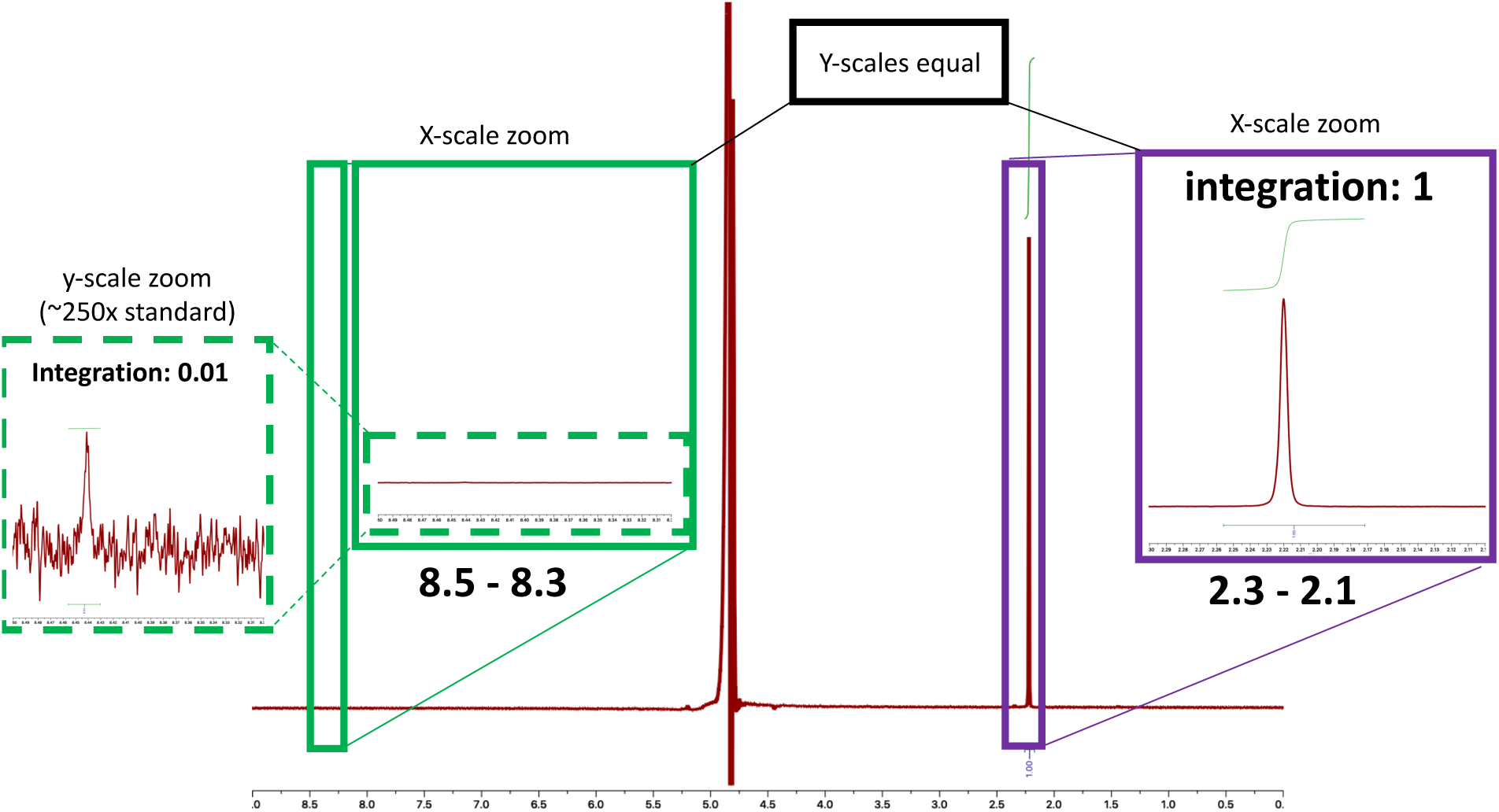
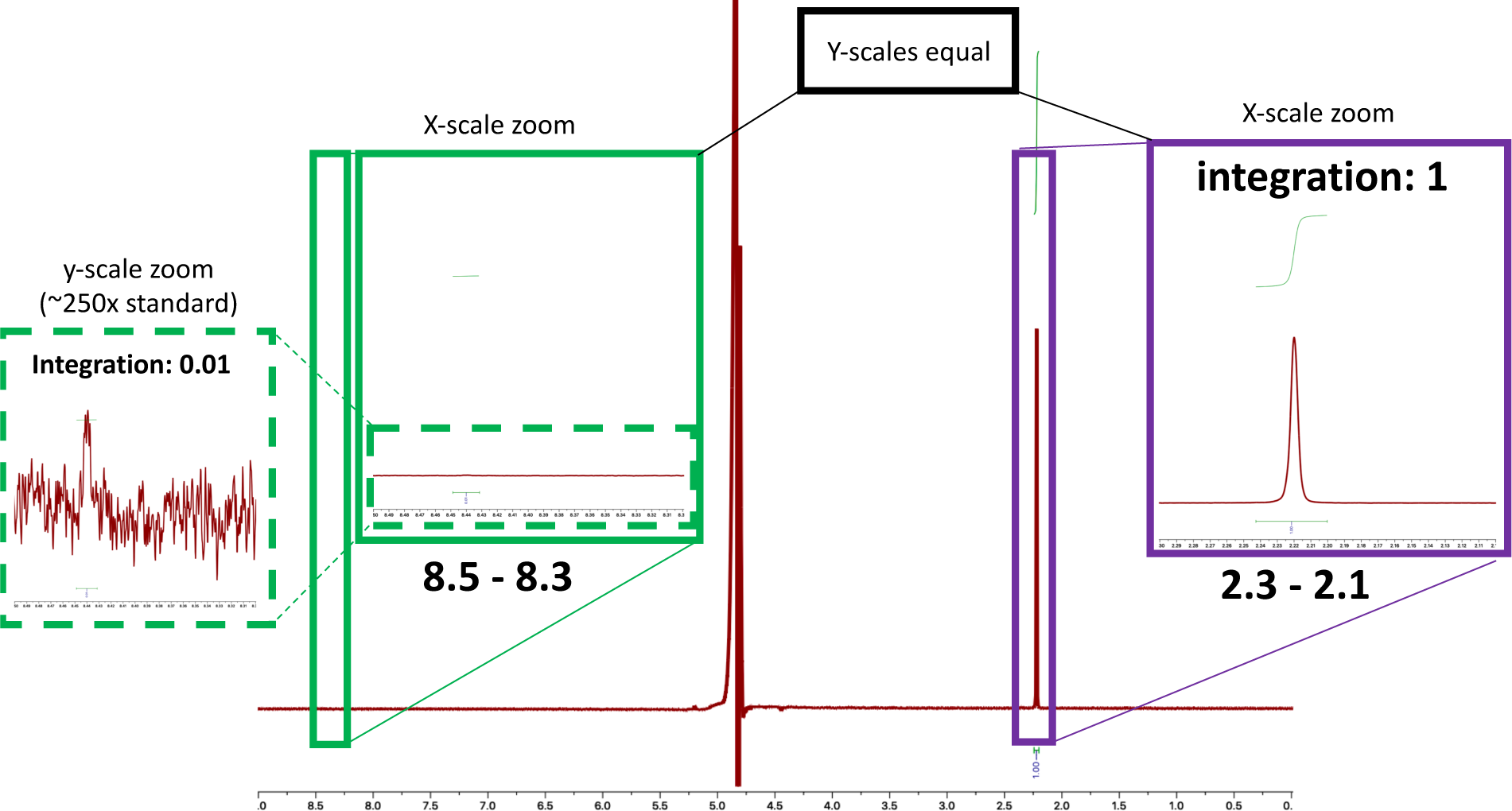
^1^H NMR spectra of Experiment 12 (only K2HPO4 in vent post-precipitation fluid) Acetone 0.6 µM was added as an internal standard. **A)** Integration of the acetone peak (1.00; 6 protons) relative to the formyl peak (0.01; 1 proton), indicates that the formate peak is lower than the limit of quantification (see section 5) **B)** Duplicate. Integration of the acetone peak (1.00; 6 protons) relative to the formyl peak (0.01; 1 proton), indicates that the formate peak is lower than the limit of quantification (see section 5). Both samples indicate formate above the detection limit, but below the limit of quantification (see section 5).

**Figure S16.**
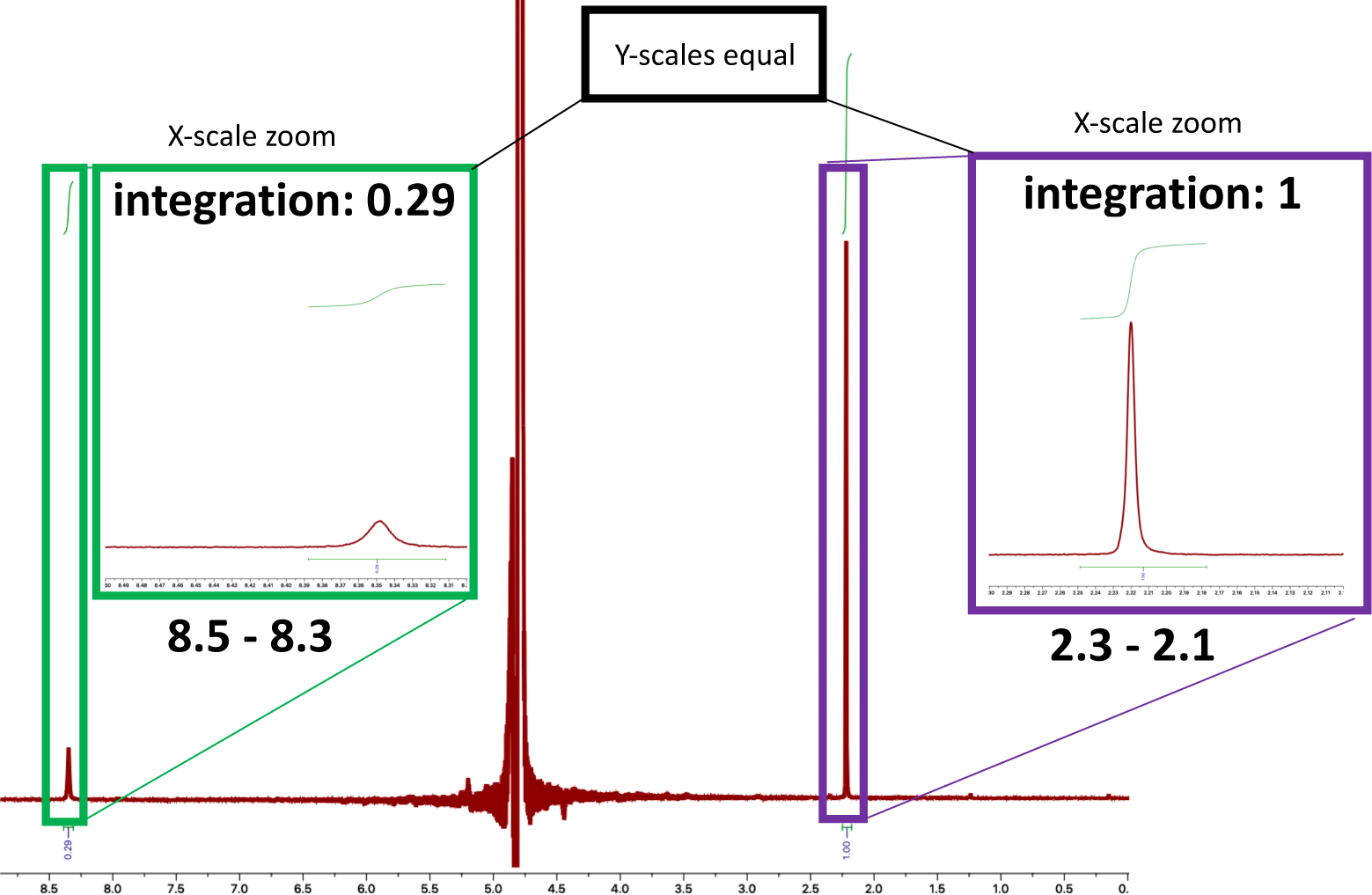
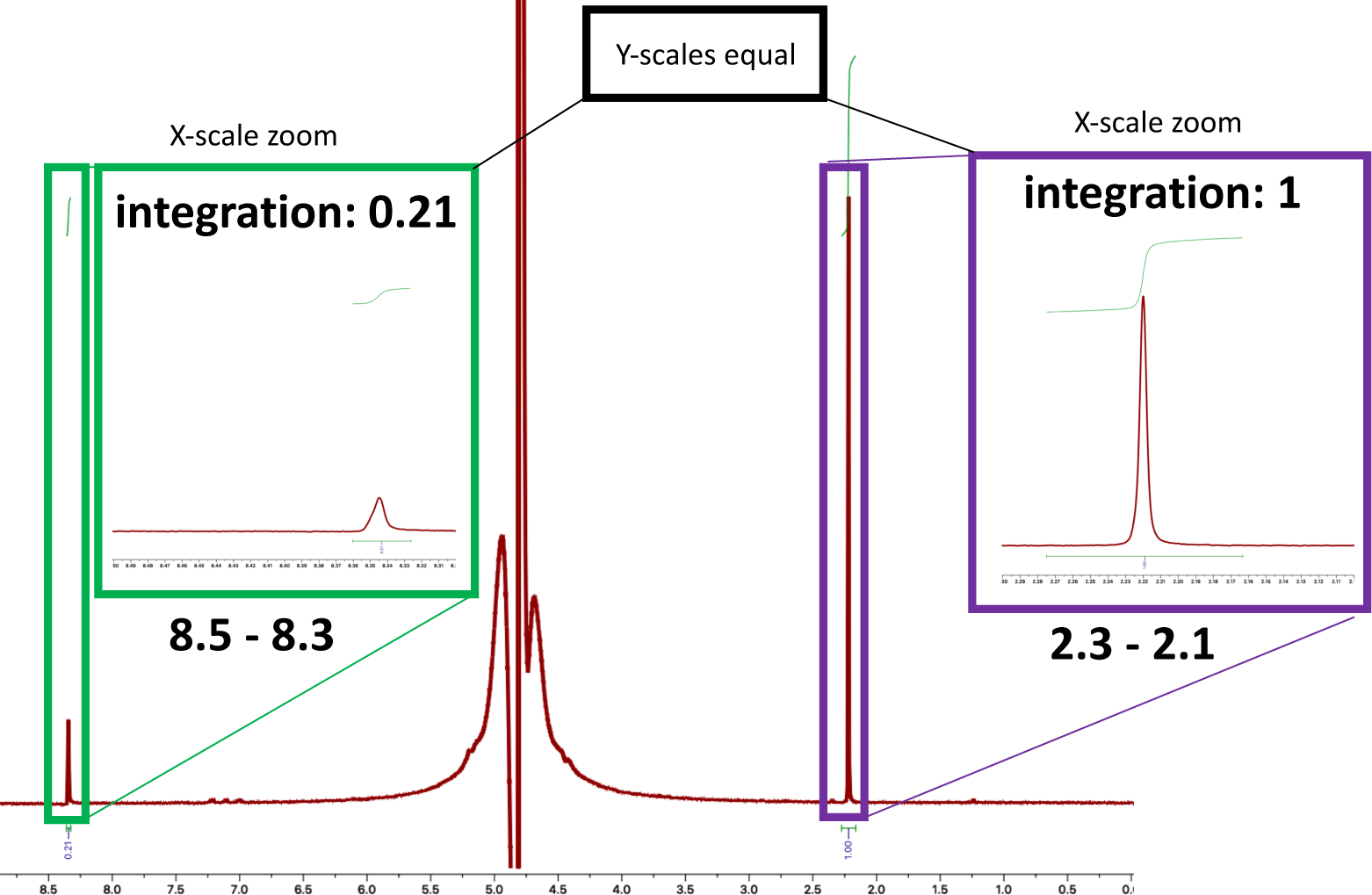
^1^H NMR spectra of Experiment 13 (only K3PO4 in vent post-precipitation fluid) Acetone 0.6 µM was added as an internal standard. **A)** Integration of the acetone peak (1.00; 6 protons) relative to the formyl peak (0.29; 1 proton), indicates a 1.044 µM concentration of formic acid. **B)** Duplicate. Integration of the acetone peak (1.00; 6 protons) relative to the formyl peak (0.21; 1 proton), indicates a 0.756 µM concentration of formic acid. An average of the two samples indicates a 0.90 µM concentration of formic acid.

**Figure S17.**
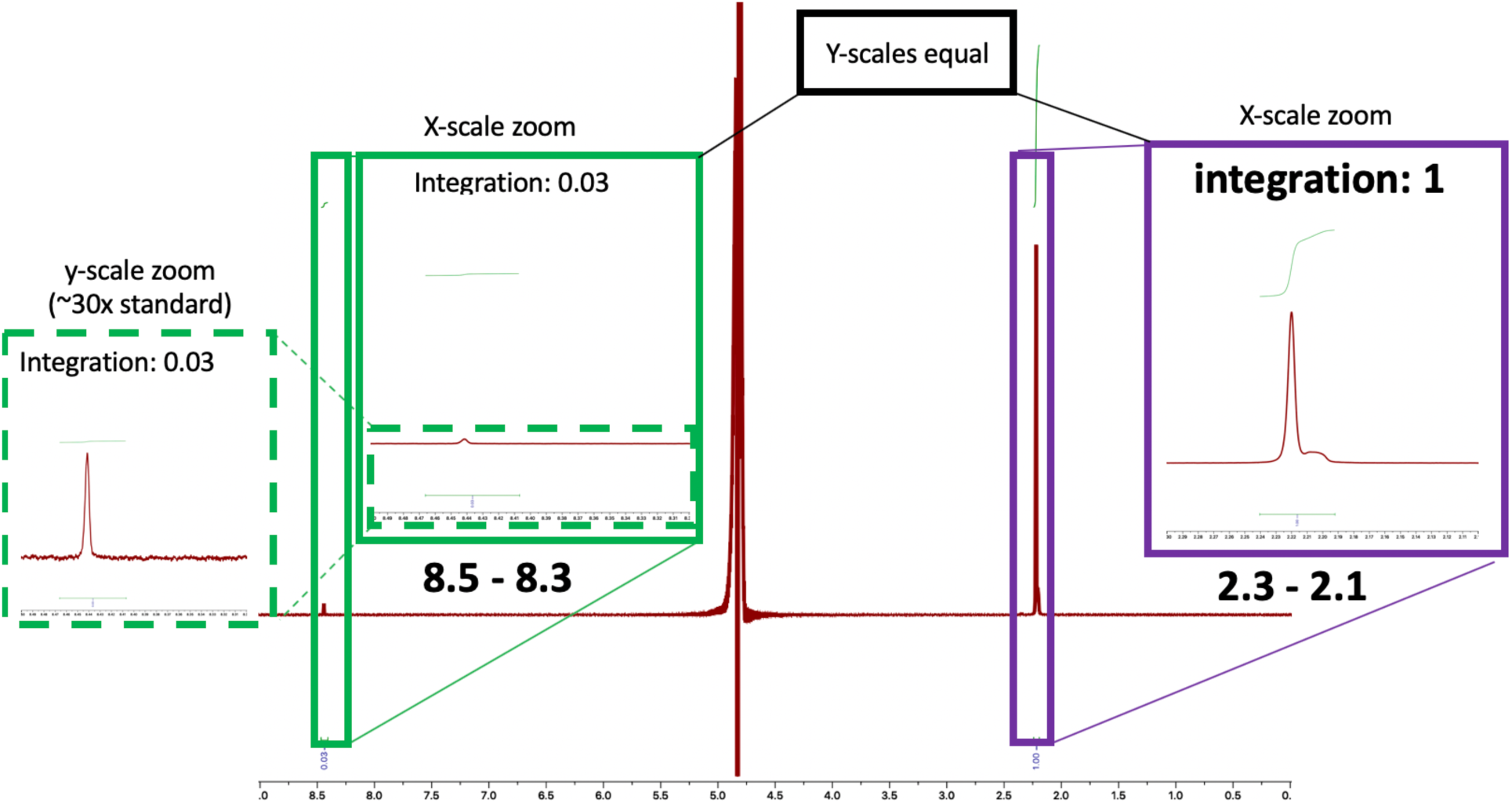
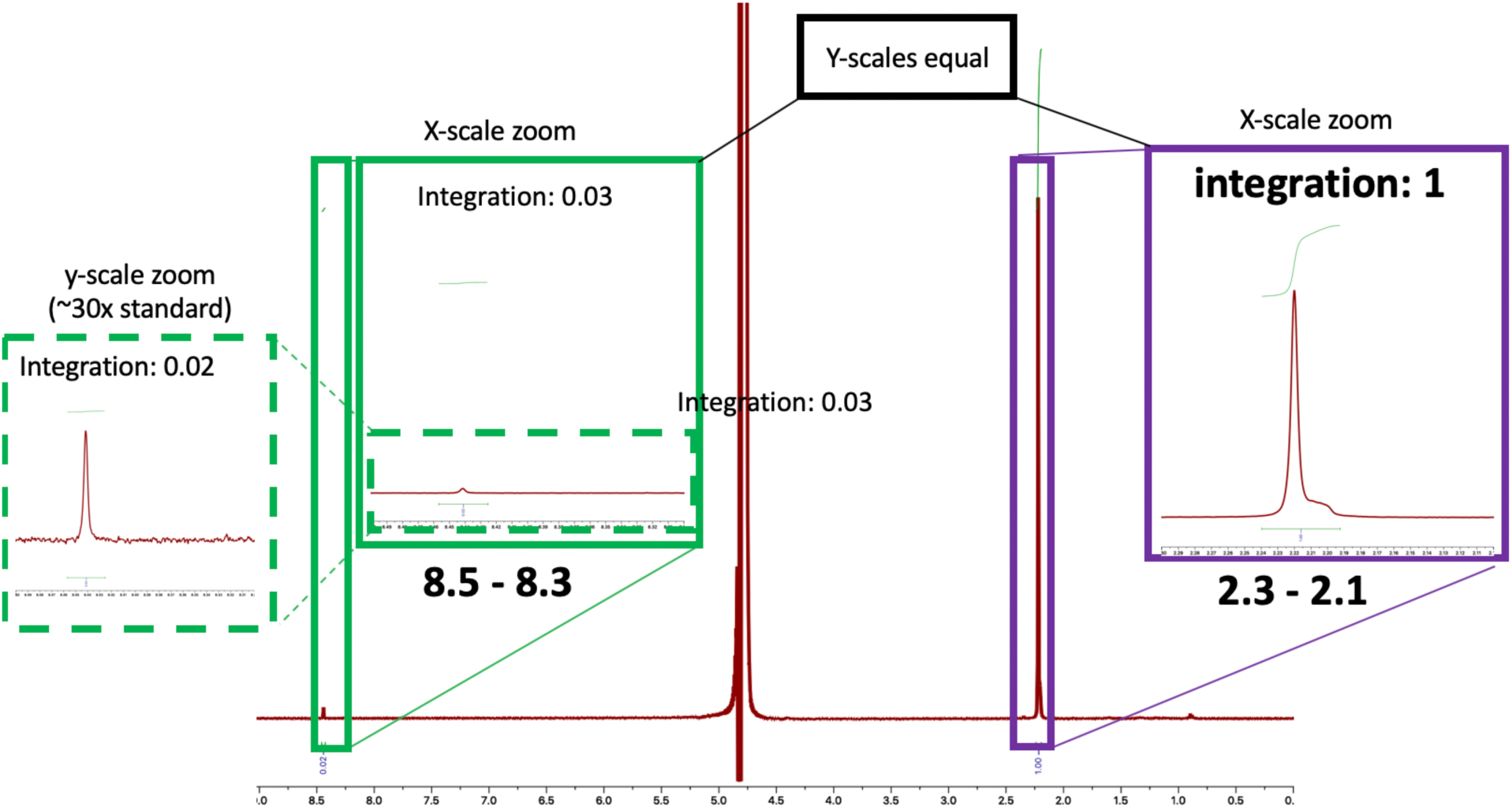
^1^H NMR spectra of Experiment 14 (only FeCl_2_ in ocean precipitation fluid) NiCl_2_ was removed from the ocean precipitation fluid. Acetone 0.6 µM was added as an internal standard. **A)** Integration of the acetone peak (1.00; 6 protons) relative to the formyl peak (0.03; 1 proton), indicates that the formate peak is lower than the limit of quantification (see section 5). **B)** Duplicate. Integration of the acetone peak (1.00; 6 protons) relative to the formyl peak (0.02; 1 proton), indicates that the formate peak is lower than the limit of quantification (see section 5). Both samples indicate formate above the detection limit, but below the limit of quantification (see section 5).

**Figure S18.**
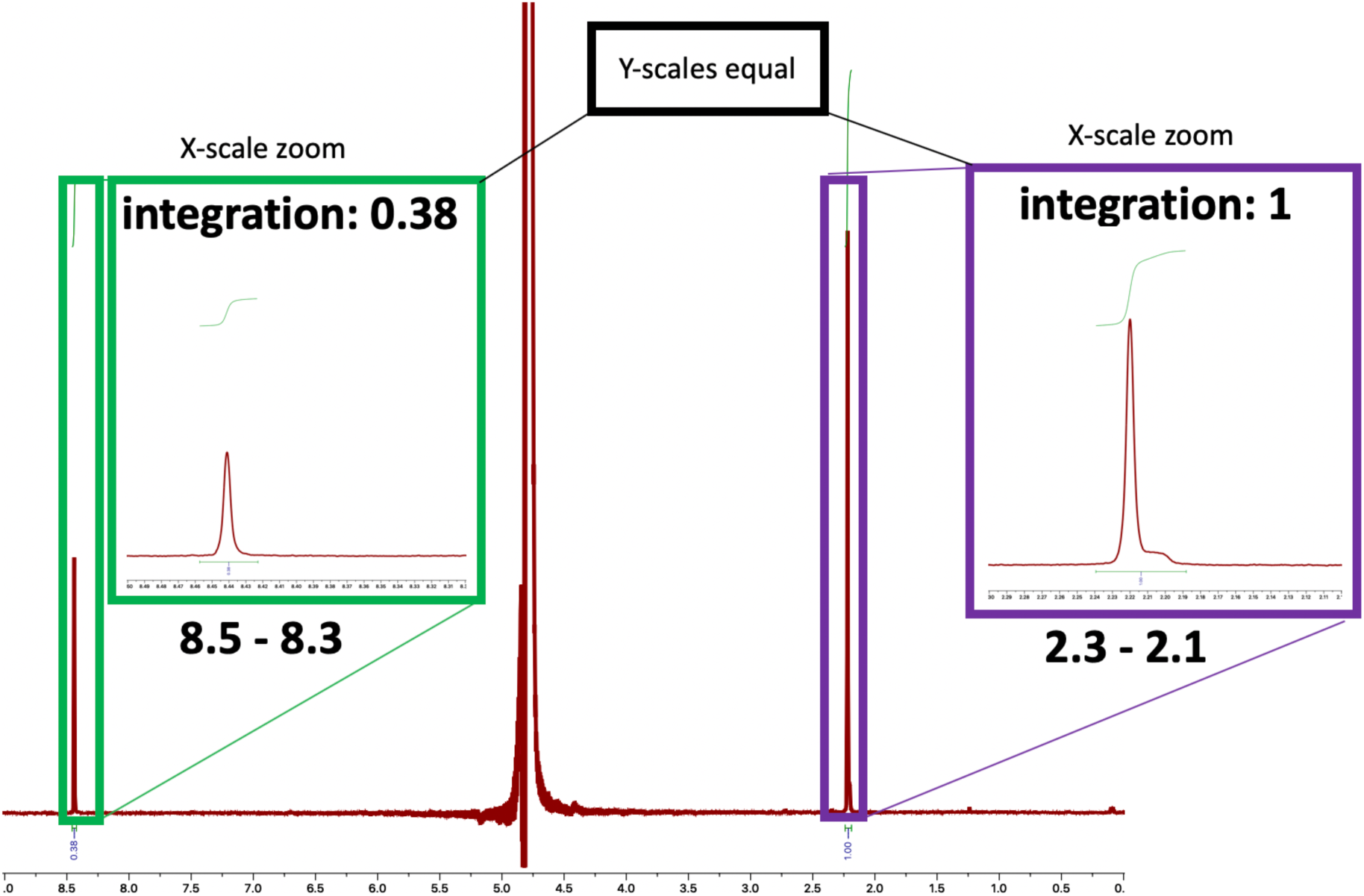
^1^H NMR spectra of Experiment 15 (only NiCl_2_ in ocean precipitation fluid) FeCl_2_ was removed from the ocean precipitation fluid. NiCl_2_ concentration was increased to 55 mM (up from 5 mM) to compensate for the missing 50 mM FeCl_2_. Acetone 0.6 µM was added as an internal standard. Integration of the acetone peak (1.00; 6 protons) relative to the formyl peak (0.38; 1 proton), indicates a formate concentration of 1.4 µM.

**Figure S19.**
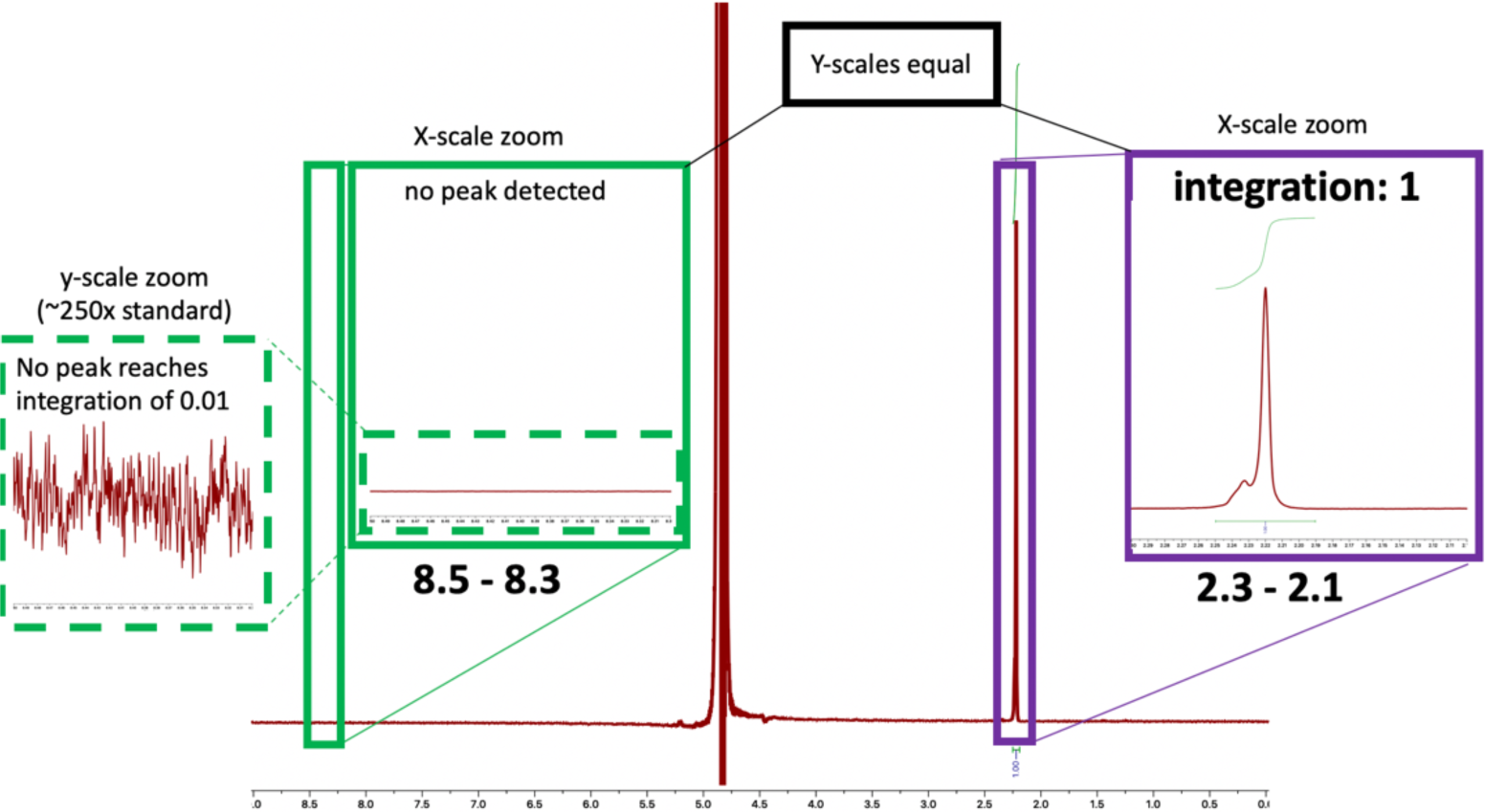
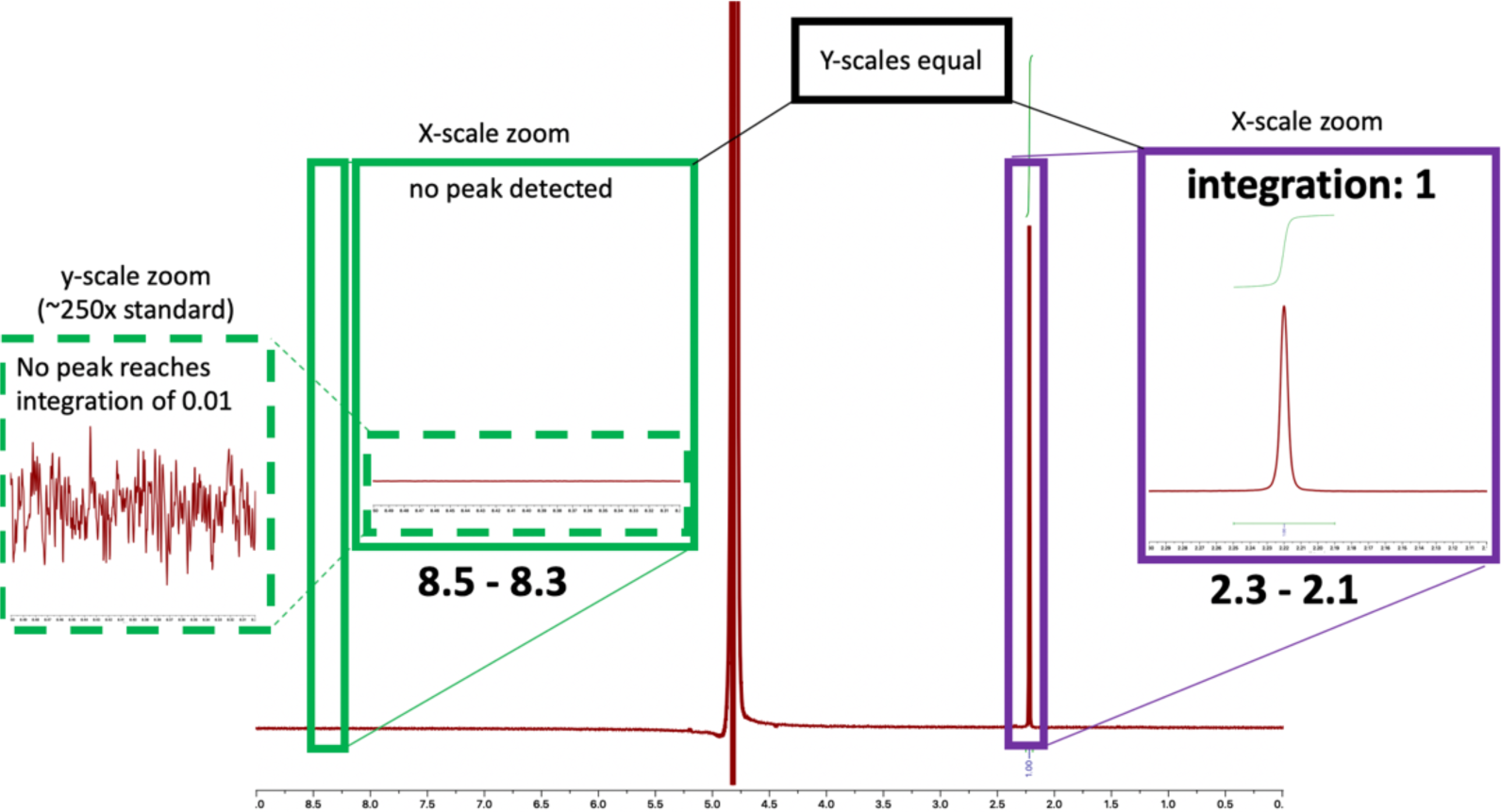
^1^H NMR spectra of Experiment 16 (neither FeCl_2_ nor NiCl_2_, i.e. no precipitate) By removing both FeCl_2_ and NiCl_2_ from the ocean precipitation fluid, no precipitate was formed. Acetone 0.6 µM was added as an internal standard. **A)** No formate peak was detected. **B)** Duplicate. No formate peak was detected.

### 5. ^1^H NMR determination of the Limit of Quantification (LoQ)

**Table S2.** **Standard curve for quantification of formate with acetone as internal standard** The Limit of Quantification (LoQ) was determined by generating a standard curve of formate relative to a constant concentration of acetone.

| [acetone]<br>($\mu\text{M}$) | acetone<br>integration | formate<br>integration | actual [formate]<br>($\mu\text{M}$) | Calc. [formate]<br>(relative to acetone<br>integration) | Quantifiable? |
| --- | --- | --- | --- | --- | --- |
| 0.6 | 1 | 3.33 | 12 match → | ← match 11.98 | >LoQ |
| 0.6 | 1 | 1.03 | 3.7 match → | ← match 3.71 | |
| 0.6 | 1 | 0.11 | 0.37 match → | ← match 0.39 | LoQ |
| 0.6 | 1 | 0.04 | 0.037 no match → | ← no match 0.14 | <LoQ |
| 0.6 | 1 | 0.01 | 0.0037 no match → | ← no match 0.036 | |

For the top 3 points, the calculated concentration of formate (relative to the acetone internal standard) matches the actual concentration of formate. For the bottom two points, the calculated concentration of formate no longer corresponds to the actual concentration of formate. The limit of quantification (LoQ) is therefore determined to be the most dilute solution of formate ( $0.37 \mu\text{M}$ ) where the concentration can be reliably calculated by comparison to the acetone internal standard.

### 6. Reactor chip and precipitate images

**Figure S20.**
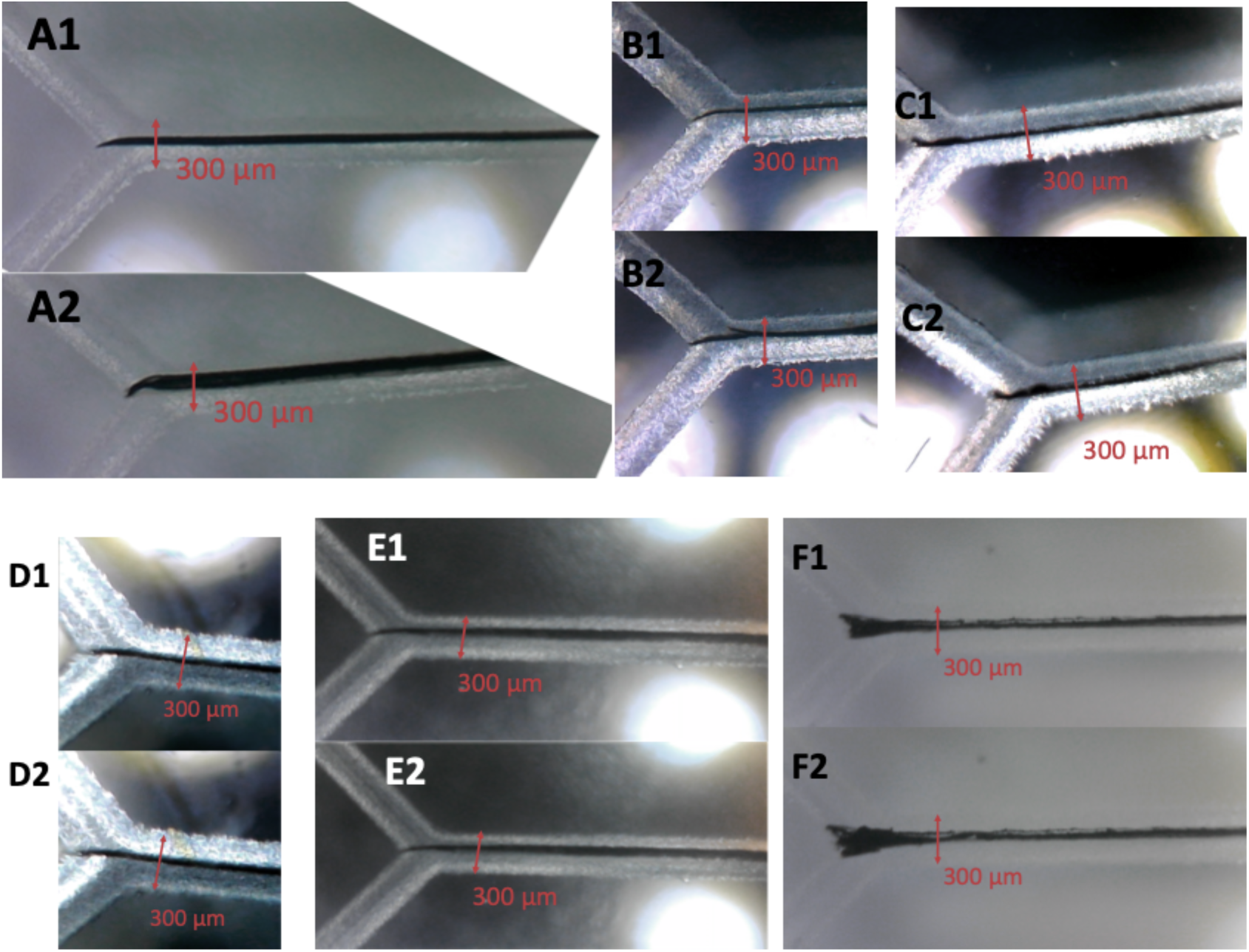
Reactor chip and precipitate images. **A)** Precipitate before (A1) and after (A2) the reaction in experiment #1 (standard reaction and precipitation conditions). **B)** Precipitate before (B1) and after (B2) the reaction in experiment #8 (standard precipitation conditions, but with the vent-side solution titrated to a pH of 3.9 during the post-precipitation conditions. **C)** Precipitate before (C1) and after (C2) the reaction for experiment #11 (standard precipitation conditions, but with Na_2_S as the only vent-side solute post-precipitation). **D)** Precipitate before (D1) and after (D2) the reaction in experiment #13 (standard precipitation conditions, but with K_3_PO_4_ as the only vent-side solute post-precipitation). **E)** Precipitate before (E1) and after (E2) the reaction for experiment #12 (standard precipitation conditions, but with K_2_HPO_4_ as the only vent-side solute during the reaction) **F)** Precipitate before (F1) and after (F2) the reaction for experiment #15 (precipitation conditions in which NiCl_2_ is the only ocean-side metal solute during the precipitation, and otherwise standard reaction solutes).

### 7. Depictions of plausible alternative CO_2_-reduction mechanisms

Multiple mechanisms are plausible for the H_2_-powered reduction of CO_2_ that we have observed here. In this following section we discuss a number of them and why we conclude that none of them are more likely than the electrochemical mechanism that we have proposed in Figure 1A of the main text.

**Figure S21.**
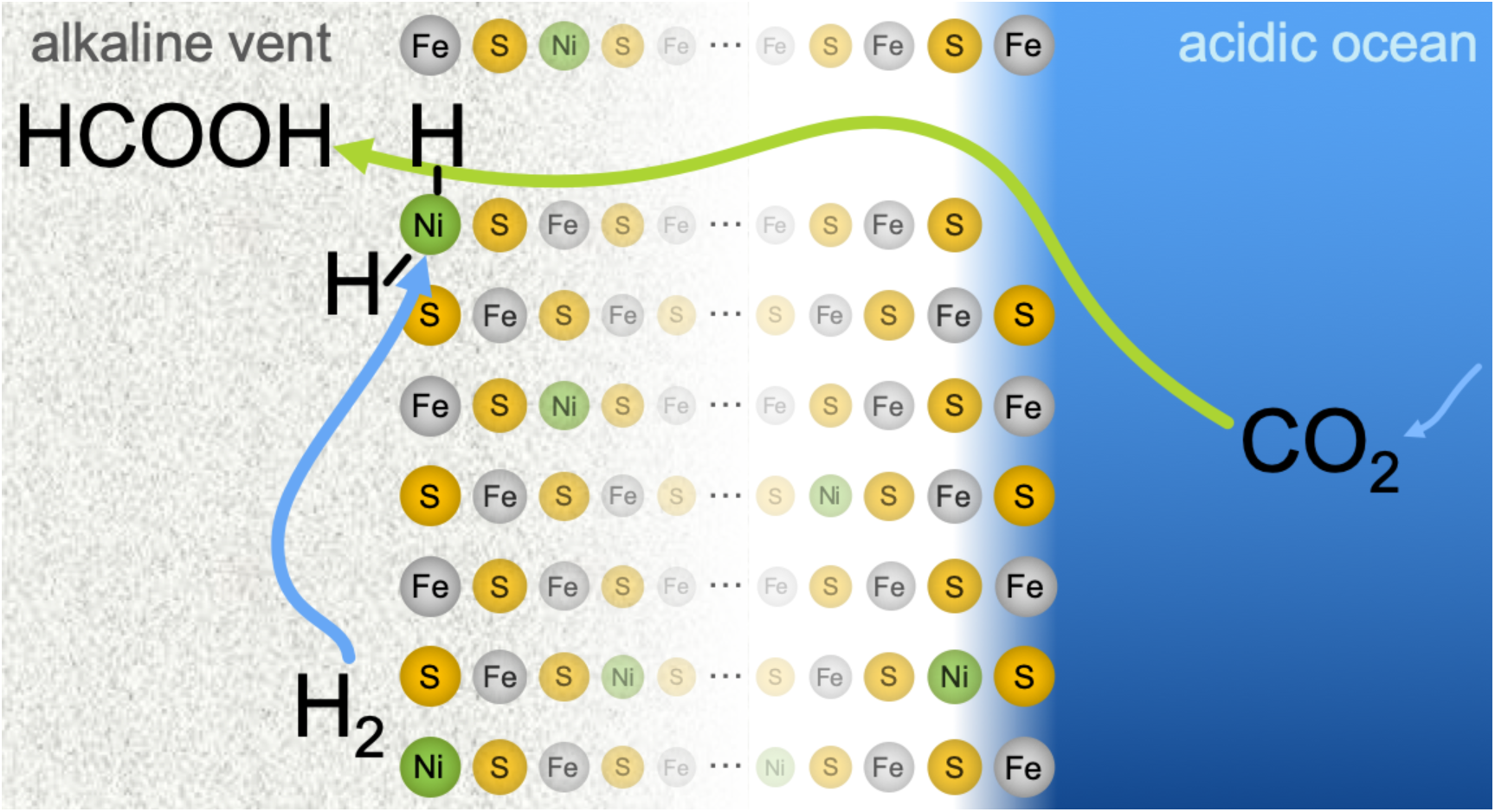
Classical hydrogenation – CO_2_ permeability.

In a system with high gas permeability, CO_2_ could permeate from the ocean side to the vent side. Upon reaching the vent side, it could interact with dissociatively adsorbed surface and subsurface atomic H (originating from the H_2_-rich vent fluids).

**Passing across the precipitate:** CO_2_ (from the ocean to the vent side, through a pore).

**Plausibility (given isotopic labeling results):** Highly unlikely. Our isotopic labeling experiments indicate that the formyl H on the produced formate derives not from the feed gas on the vent side (H_2_/D_2_), but instead specifically from the ocean-side water (H_2_O/D_2_O). Therefore, with our microfluidic system, such a mechanism is not possible.

#### Ionic Hydrogenation (alternative)

A putative ionic hydrogenation would proceed similarly to the above classical hydrogenation but, rather than dissociatively adsorbing on the precipitate as a pair of hydrogen atoms, the H_2_ would formally reduce the precipitate by generating a hydride at the surface. The hydride would then transfer to the incoming CO_2_.

**Plausibility (given isotopic labeling results):** Highly unlikely.

#### Possibility of H exchange with the solvent in direct hydrogenations

Adsorbed H· or H^−^ in a classical or ionic hydrogenation, respectively, could in principle exchange with the surrounding aqueous environment, so that we lose the original isotopic signal (reference (32) in the main text). However, this would inevitably imply considerable mixing of the two fluids, with a correspondingly mixed H/D signal in the product that we do not observe.

**Figure S22.**
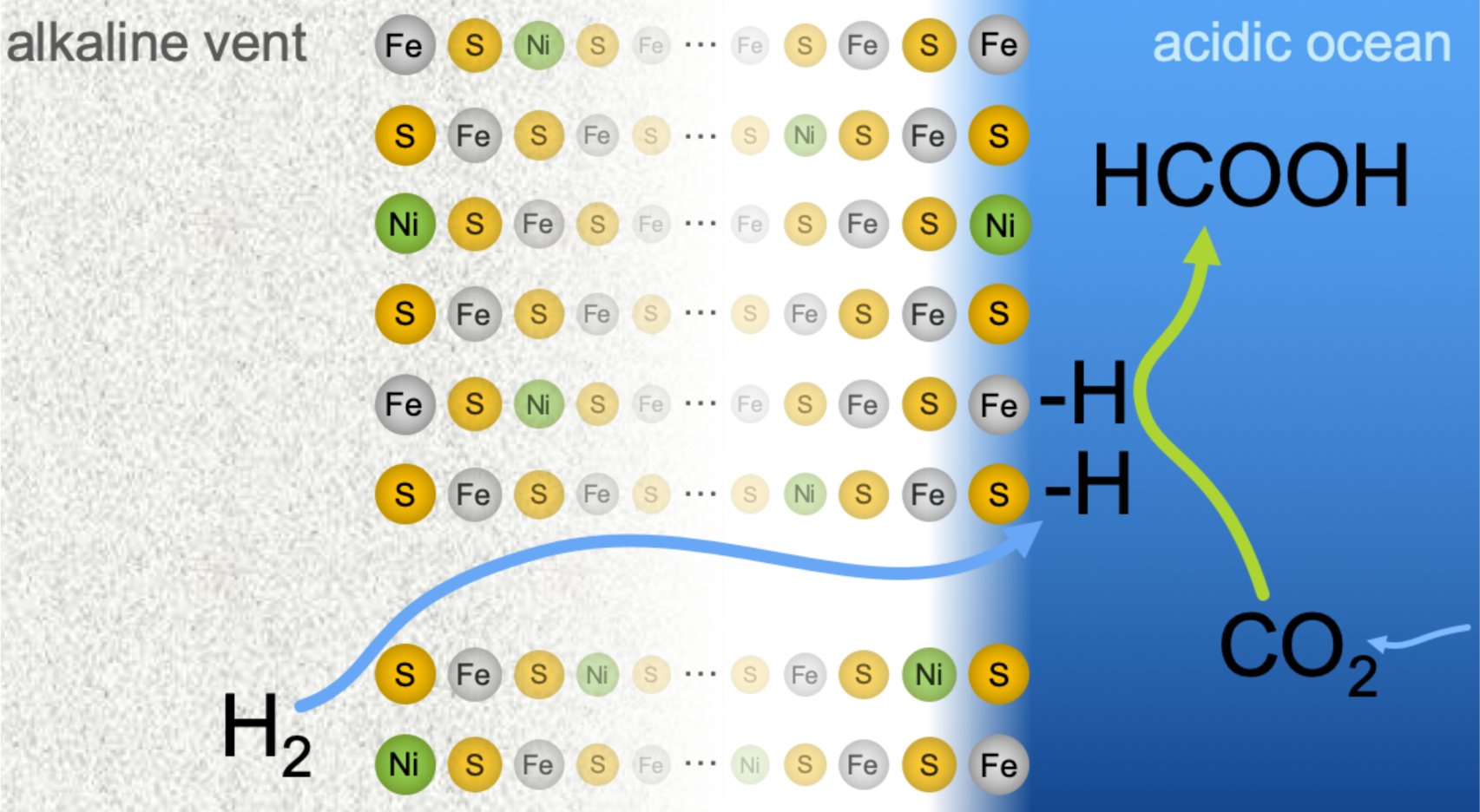
Classical hydrogenation – H_2_ permeability.

In a system with high gas permeability, H_2_ could permeate from the ocean side to the vent side. Upon reaching the vent side, it could interact with dissociatively adsorbed surface and subsurface atomic H (originating from the H_2_-rich vent fluids).

**Passing across the precipitate:** H_2_ (from the vent to the ocean side, through a pore).

**Plausibility (given isotopic labeling results):** Highly unlikely. As above, our isotopic labeling experiments indicate that the formyl H on the produced formate derives not from the feed gas (H_2_/D_2_), but instead specifically from the ocean-side water (H_2_O/D_2_O). Therefore, with our microfluidic system, such a mechanism seems impossible.

#### Ionic hydrogenation (alternative)

Following migration of H_2_ to the acidic side, an ionic-hydrogenation mechanism would proceed via adsorption of a hydride at the surface of the precipitate with release of a proton into the surrounding fluid (instead of a homolytic adsorption of a pair of neutral hydrogen atoms). The hydride could then potentially exchange with the surrounding aqueous environment, confounding our isotopic signal.

**Plausibility (given isotopic-labeling results):** Highly unlikely. If D_2_ in experiment #4 had migrated to the ocean side and then become absorbed into the precipitate, the bound deuteride could have exchanged with the local environment (H_2_O), giving the unlabeled signal that we observe. However, the migration of the dissolved hydrogen to the opposite side must have involved considerable mixing of fluids. So, if this were the mechanism, in experiment #5 we would have generated a mixture of H_2_O and D_2_O, with the corresponding DCOO^−^ and HCOO^−^ (representing a mixture of exchanged and unexchanged hydrides) in the product. Instead, we observe only DCOO^−^, suggesting that the fluids did not significantly mix prior to reaction.

**Figure S23.**
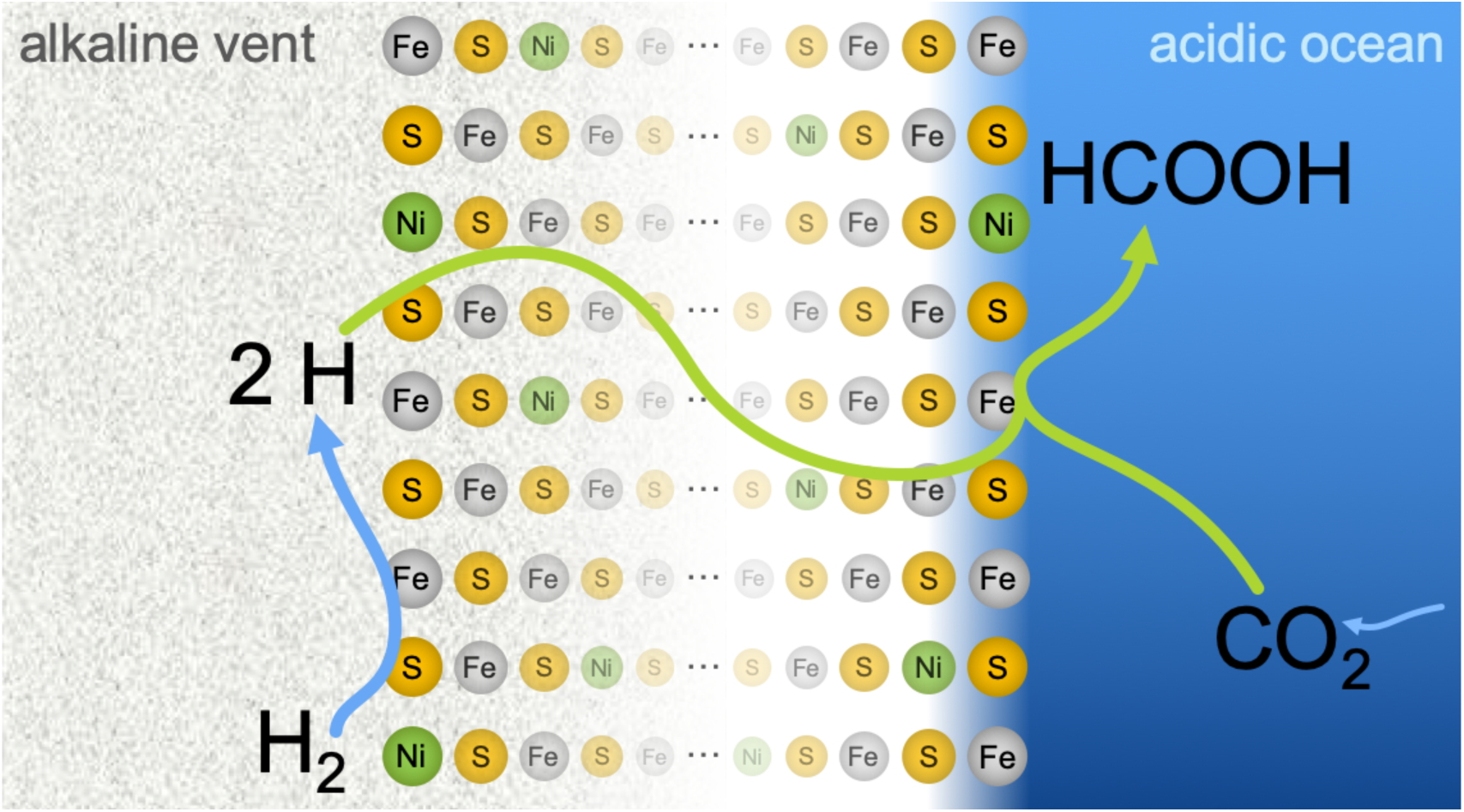
Classical hydrogenation – passage of dissociated atomic H.

In a system with low gas permeability, H_2_ could dissociatively adsorb on the vent side, and the atomic H could travel through the lattice of the precipitate to interact with CO_2_ on the ocean side, where it could reduce CO_2_ in classical hydrogenation.

**Passing across the precipitate:** Dissociated atomic H (from the vent to the ocean side, through the precipitate network itself).

**Plausibility (given isotopic labeling results):** Highly unlikely.

Once again, our isotopic labeling experiments indicate that the formyl H on the produced formate derives not from the feed gas (H_2_/D_2_), but instead specifically from the ocean-side water (H_2_O/D_2_O). Therefore, with our microfluidic system, such a mechanism seems impossible.

#### Ionic hydrogenation (alternative)

This proposed ionic hydrogenation would proceed similarly to the above classical hydrogenation but, rather than dissociatively adsorbing, the H_2_ could formally reduce the precipitate by generating a hydride bound at the surface. The hydride could then potentially migrate to the ocean side, where it would exchange with the surrounding aqueous environment.

**Plausibility (given isotopic labeling results):** Highly unlikely. Again, hydride exchange with the local environment (D_2_O on the ocean side) could explain the coupling of the formyl H/D signal with that of the ocean side water (DCOO when ocean side water is D_2_O), but this would likely generate a mixture of DCOO^−^ and HCOO^−^ (representing a mixture of exchanged and unexchanged hydrides from the considerable mixing of H_2_O and D_2_O in experiment #5).

**Figure S24.**
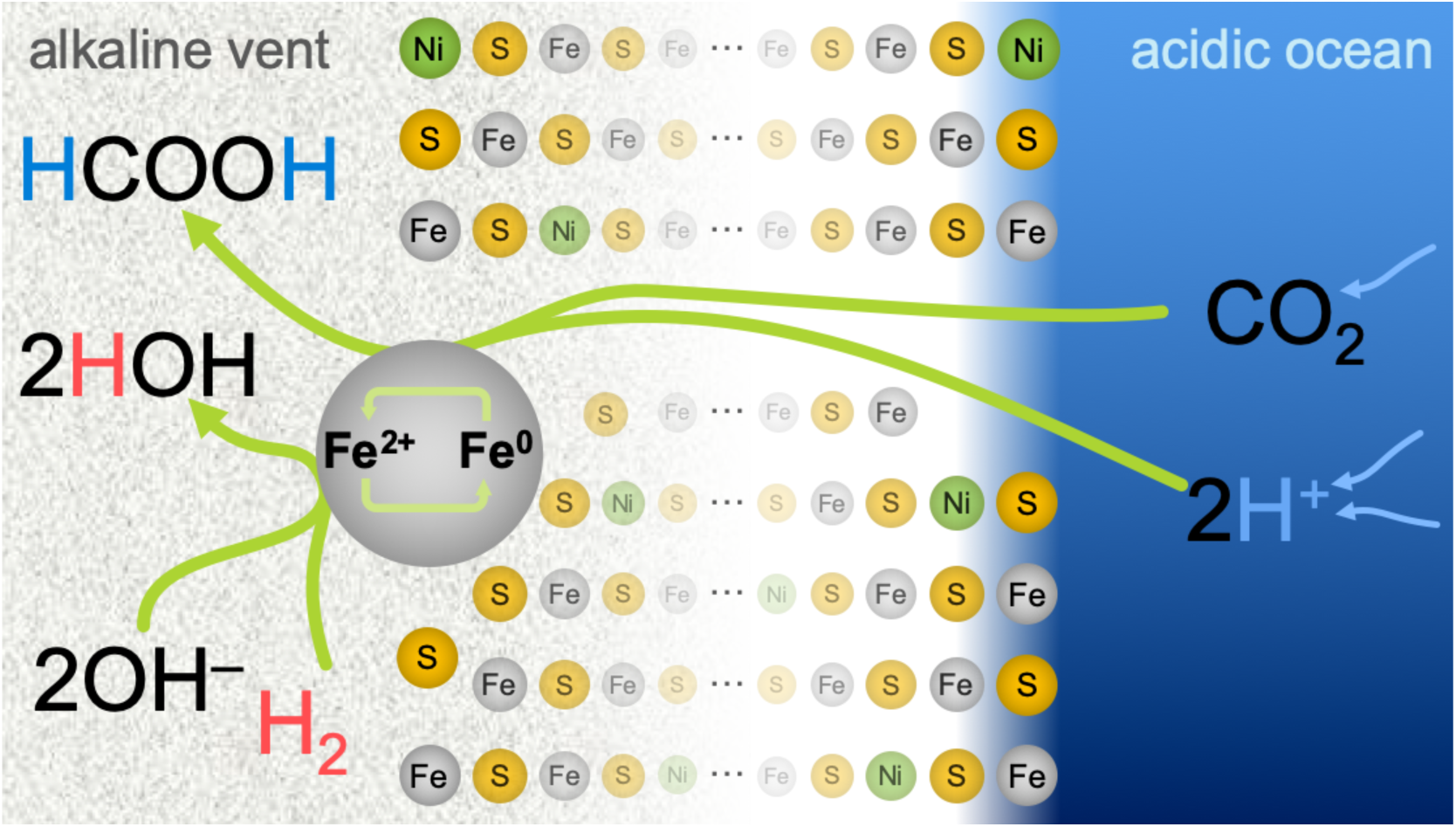
Localized redox cycling – CO_2_ and H^+^ permeability.

This mechanism relies on redox cycling of corner or edge Ni or Fe atoms (M^2+^ ⇄ M^0^) on the vent side wherein H_2_ oxidation is decoupled from CO_2_ reduction. In such a scenario, the ‘H’ incorporated into the formate would derive from the acidic ocean side rather than from H_2_. In this scenario, we consider H^+^ passing from the acidic ocean side to the alkaline vent side through hydrated microchannels.

**Passing across the precipitate:** CO_2_ and H^+^ (from the ocean to the vent side, both through a pore). Passage of H^+^ through a hydrated microchannel would involve rapid equilibration with the surrounding water. The resulting H^+^ ions should thus take on more of the deuterated isotopic make-up of the vent-side fluid as they got closer to the vent side in our experiments with D_2_ as a driver gas for the vent fluid. This mechanism should therefore result in at least a mixture of HCOO^−^ and DCOO^−^, which we do not observe (we observe pure HCOO^−^).

As mentioned in the main text, we never added acids to the ocean side; the acidic pH (typically 3.9) was achieved solely by dissolution of CO_2_ in water. Thus, in the experiments with D_2_O as ocean-side solvent, all ocean-side protons must derive from the dissociation of carbonic acid via:

D_2_O + CO_2_ ⇌ D_2_CO_3_ ⇌ **D^+^** + DCO_3_^−^

**Plausibility (given isotopic labeling results):** Highly unlikely. We would have observed a mixed D/H signal (or in fact mostly H since H_2_O was the prevalent fluid in the vent side) in experiment #5.

**Figure S25.**
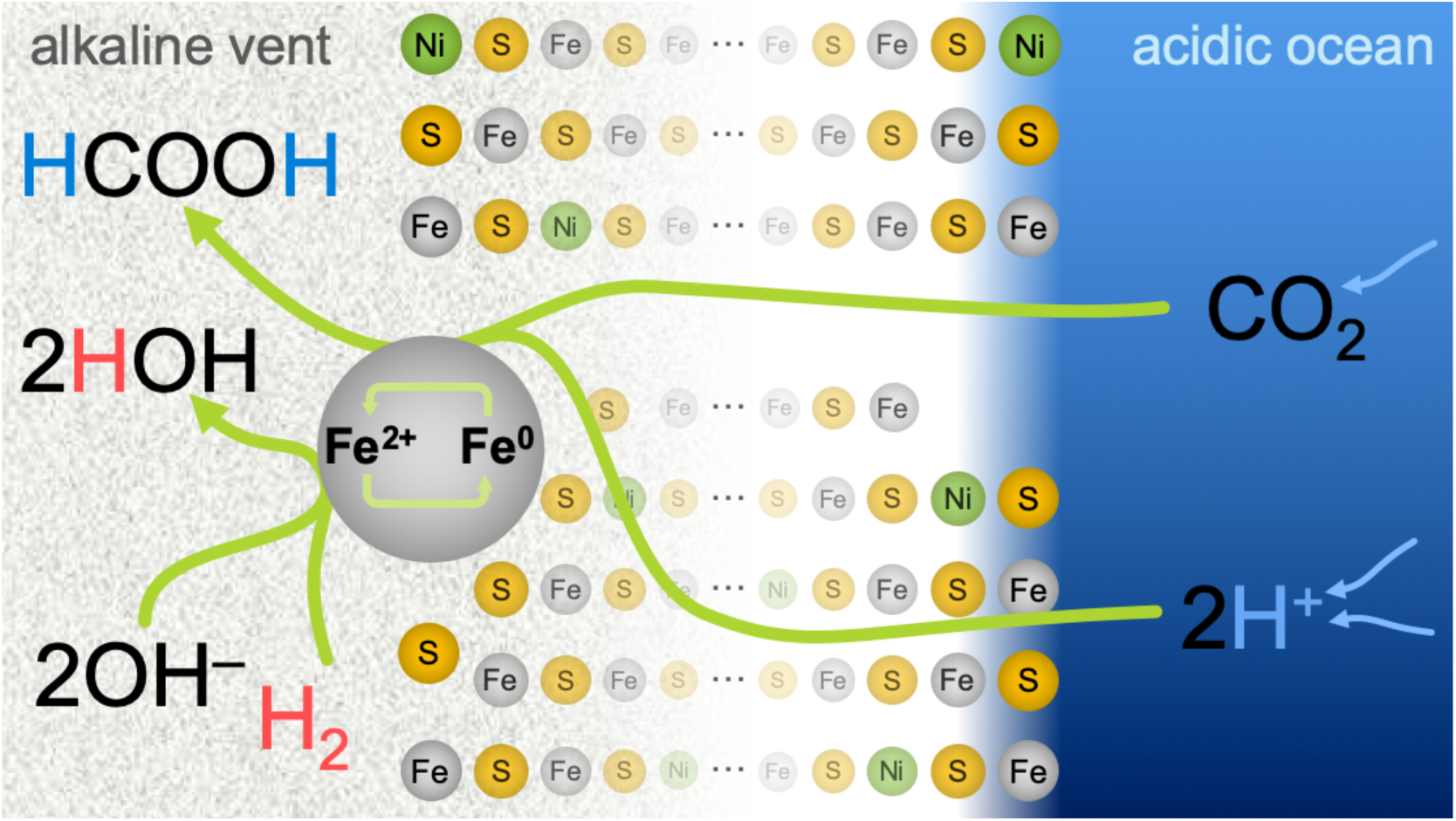
Localized redox cycling – CO_2_ permeability and H^+^ conductivity.

As in Figure S24, this mechanism relies on redox cycling of edge Ni or Fe atoms (M^2+^ ⇄ M^0^) on the vent side, with H_2_ oxidation decoupled from CO_2_ reduction. In such a scenario the ‘H’ incorporated into formate derives from the ocean side rather than from the H^+^ generated by H_2_ oxidation on the vent side. This would account for our observed deuterated product with D_2_O as the solvent in the acidic fluid (and H_2_O in the vent fluid). In contrast with Figure S24, in this scenario we consider H^+^ passing from the acidic ocean side to the alkaline vent side via anhydrous proton conduction through the precipitate lattice.

**Passing across the precipitate:** CO_2_ (from the ocean to the vent side, through a hydrated pore). H^+^ (from the ocean to the vent side, anhydrously through the precipitate network itself).

In an anhydrous proton-conduction mode, protons would not equilibrate with the surrounding aqueous environment, allowing for the observed HCOO^−^ in the experiments with D_2_ as vent driver. However, it seems highly unlikely that our precipitates could be aqueous and porous enough to permit dissolved CO_2_ to migrate to the alkaline side, but simultaneously have anhydrous enough sections to allow for dry proton conduction without equilibration to surrounding D_2_O (in the experiments with D_2_O as vent solvent).

Separately, our results indicate the need for a large pH gradient between the two semi-reactions. Keeping such a gradient at the nano-scale seems unlikely, but remains an open question (see the discussion in reference (25) in the main article).

**Plausibility (given isotopic labeling results and need for pH gradient):** Possible, but unlikely.

**Figure S26.**
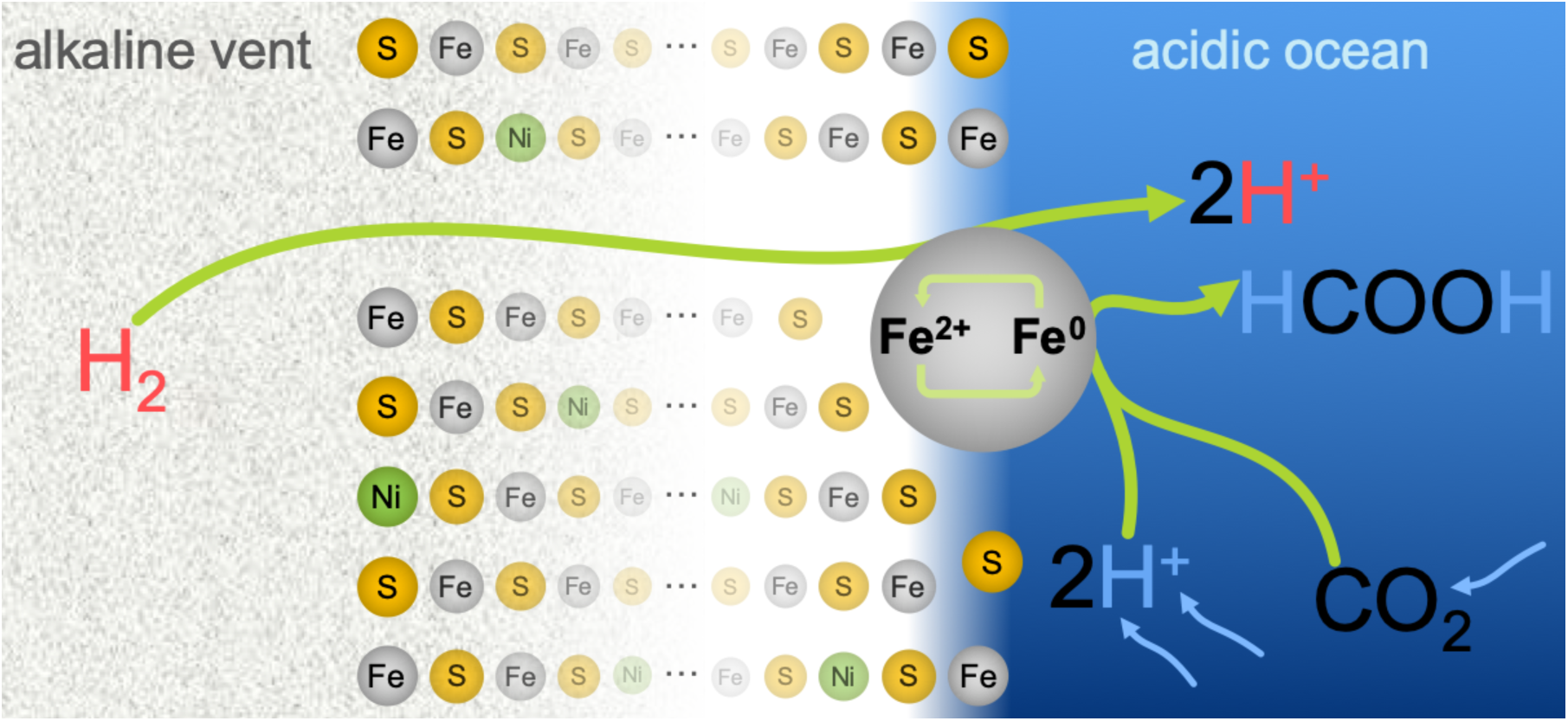
Localized redox cycling – H_2_ permeability.

This mechanism relies on a redox cycling of corner or edge Ni or Fe atoms (M^II^ßà M^0^) on the ocean side wherein H_2_ oxidation is decoupled from CO_2_ reduction. In such a scenario the ‘H’ incorporated into the formate would derive from the acidic ocean side rather than the H^+^ generated from H_2_ oxidation.

**Passing across the precipitate:** H_2_ (from the vent to the ocean side).

This mechanism would agree with our isotopic results, assuming the pair of protons (H^+^) released from the initial H_2_ oxidation fully diffuse away before the CO_2_ reacts with a fresh pair taken from the surroundings (otherwise we should have detected deuterated product when pushing the vent fluid with D_2_). However, this mechanism would require H_2_ oxidation on the ocean side, which is far less favorable than on the vent side, so we deem this process unlikely in our system.

**Plausibility (given isotopic labeling results and need for pH gradient):** Possible, but unlikely.

In conclusion, an electrochemical mechanism (main text, Figure 1A) seems the most likely mechanistic scenario, avoiding the implausibilities of the various classical and transfer hydrogenation alternatives described above.

### 8. Finite-element computer simulation of Venturi flow in a hydrothermal pore

We modeled a microfluidic pore narrowing from 2 mm into a channel 300 µm wide, before expanding again. The narrow section is connected to a perpendicular side pore 200 µm in diameter, which is itself assumed to lead into a reservoir (such as the ocean in our case). The narrowing at the bottom causes a Bernoulli drop in pressure, which leads to Venturi pull from the side reservoir into the hydrothermal channel system (Figure S27).

**Figure S27.**
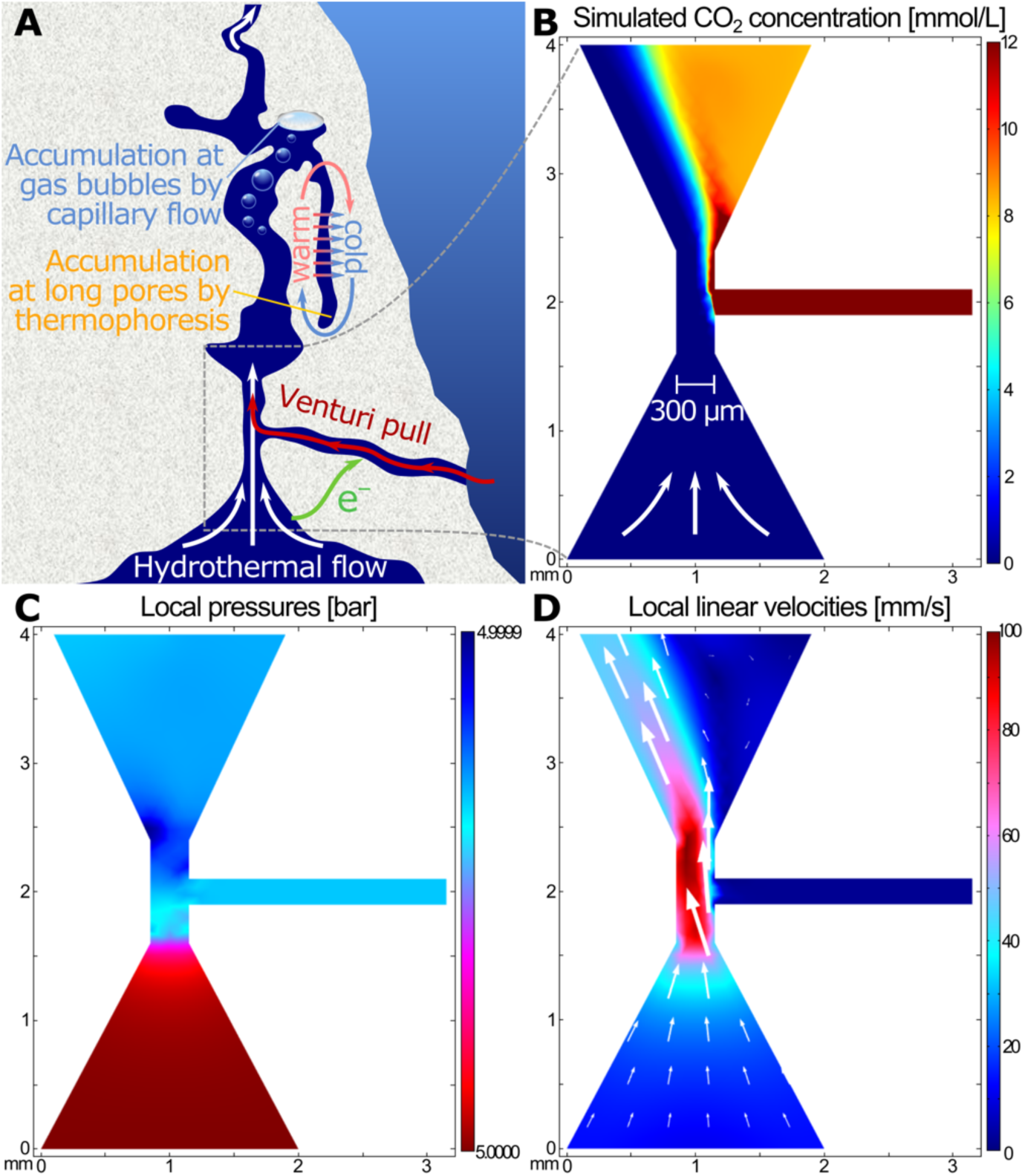
Simulation of hydrothermal-driven Venturi flow. **(A)** Narrowing of hydrothermal flow entering from the bottom causes a Bernoulli-principle drop in pressure, leading to Venturi pull from an open reservoir (e.g. ocean) connected to the side via a separate pore. Reagents (such as Fe^2+^, CO_2_ and H^+^) could enter the microporous hydrothermal system in this way, and once inside react either with electrons (e^−^) transferred through the electroconductive mineral, or upon meeting the hydrothermal flow at the center. Further up, thermal gradients at gas bubbles (41) and elongated pores (39, 40) can lead to concentration increases of any organics produced. **(B)** Steady-state finite-element simulation of concentration changes of CO_2_ dissolved in a fluid entering from a side pore leading to a reservoir on the right (purported ocean). **(C)** Local pressures in the system. Even the minuscule 0.01% drop in pressure in the simulation here produces enough drag to pull fluids into the system, as shown in (B). **(D)** Local velocities in the system confirm the Bernoulli-principle behavior that leads to pull of fluids from the side pore.

As the H_2_ dissolved in the hydrothermal fluid permeates upward through the vent, a decrease in pressure would lead to the spontaneous production of gas bubbles. A similar situation would occur as the CO_2_ dissolved in the ocean was heated upon contact with the warmer vent fluid, releasing further gas bubbles. Flowing up, the bubbles would collect inside some of the vent’s microchambers (Figure S27A, top center). At the gas-liquid interfaces, heated by the hydrothermal flow on one side and cooled by the ocean’s proximity on the other, continuous capillary flow would facilitate vast increases in concentration and promotion of complex reactivity (41). Separately, the heat gradients would also cause thermal convection and diffusion at elongated pores, further increasing concentrations of organics by thermophoresis (39, 40) (Figure S27A, top right).

The finite-element computational simulation was performed in COMSOL Multiphysics v4.4 including the Laminar Flow and Transport of Diluted Species modules, with slip walls under a stationary solver with default options. The liquid was defined as incompressible. Temperature was defined uniformly as 75 °C (323.15 K). Water-solvent and substance properties were set to COMSOL defaults. Pressure at the base of the hydrothermal system was defined as 5 bar (equivalent to ∼40 m under sea level at present). To study the effect of Venturi pull, the pressures of the side pore and the top outlet were defined equal to each other, as 0.01% lower than the pressure of the hydrothermal inflow at the base.

The diffusion coefficient of CO_2_ was calculated as described in (45) using:

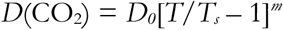

where *D_0_* = 13.942·10^−9^ m^2^/s; *T_s_* = 227.0 K; and *m* = 1.7094.

